# Integrating morphological and molecular approaches to assess photosynthetic species diversity in the Venice Lagoon

**DOI:** 10.64898/2026.10.12.756211

**Authors:** Filippo Drigo, Riccardo Trentin, Chiara Frasson, Chiara Stefanelli, Emanuela Moschin, Davide De Battisti, Cristiano De Pittà, Isabella Moro

## Abstract

The Venice Lagoon is the largest transitional environment in the Mediterranean Sea and an important biodiversity hotspot. However, it is under increasing climate and anthropogenic pressures, highlighting the need for long-term monitoring of its biodiversity.

Three monitoring campaigns were conducted between 2023 and 2025, sampling seaweeds, seagrasses and halophytes. Samples were identified using an integrated morphological and molecular approach, with morphological traits guiding the selection of the most appropriate DNA-barcoding markers. High-quality sequences were obtained for 50% of the samples and compared with reference databases. To improve identification reliability, phylogenetic trees were reconstructed for each major taxonomic group to assess clustering patterns and sequence relationships. Two independent species-delimitation methods, ASAP and GMYC, were also applied.

Overall, 81 taxa were identified, 13 belonging to Chlorophyta, 11 to Heterokontophyta, 28 to Rhodophyta, and 29 to Tracheophyta. We further conducted a targeted survey on *Sporobolus* species using the *trnL-trnF* marker and genome-size estimates to distinguish between native and non-indigenous ones. The presence of *S. x townsendii* was confirmed in the central and southern lagoon, while the non-indigenous *S. anglicus* was more prevalent than the native *S. maritimus*.

**Highlights:**

1. An integrated approach assessed photosynthetic diversity in the Venice Lagoon.
2. Eighty-one photosynthetic species and genera were identified across major taxa.
3. *Vaucheria litorea* was recorded for the first time.
4. *Sporobolus x townsendii* was confirmed in the Venice Lagoon.
5. Non-indigenous *S. anglicus* was more widespread than native *S. maritimus*.

## 1. Introduction

Reliable species identification is crucial for monitoring species occurrence and assessing changes in biodiversity over time. However, expert taxonomists are decreasing, hampering the effectiveness of monitoring studies based on morphological identification, especially for problematic taxa, like seaweeds (Robuchon *et al*., 2015; Kowalska *et al*., 2019; Bartolo *et al*., 2020).

Molecular tools can support species identification and enable non-specialists to classify samples, thereby improving knowledge of biodiversity. However, limited taxonomic expertise may result in the misidentification of sequences deposited in public databases (Robuchon *et al*., 2015). Integrating molecular analyses with morphology-based identification using taxonomic keys can therefore improve accuracy, particularly for cryptic species and organisms with morphologically distinct life stages. This combined approach is also valuable for the early detection and management of non-indigenous species (NIS) (Kowalska *et al*., 2019).

Nowadays, different molecular tools are available, like the DNA barcoding (Bartolo *et al*., 2020; CBOL, 2009; Kowalska *et al*., 2019), which is frequently applied with a single-*locus* approach (Bartolo *et al*., 2020; Magoga *et al*., 2021; Robuchon *et al*., 2015). To overcome the limitations of one *locus* (e.g., evolutionary differences between nuclear and organelle genomes; retention of ancestral polymorphism and incomplete lineage sorting) (Bartolo *et al*., 2020; Leliaert *et al*., 2014), the use of phylogenetic reconstruction (Elias *et al*., 2007) and species delimitation approaches on the same dataset might increase the confidence of species identification (Magoga *et al*., 2021), overcoming dubious alignments and wrong identifications based only on similarity searches (e.g., BLAST alignment with GenBank database). This integrated approach is particularly important for macrophytes, whose identification is often complicated by phenotypic plasticity, cryptic diversity, and the limited availability of distinctive morphological traits.

Seaweeds, seagrasses and halophytes (referred as photosynthetic species from now on) are primary producers and provide various ecosystem services; two examples are nursery areas crated by canopy forming species and sediment stabilization by seagrasses and halophytes (Macreadie *et al*., 2017; Sfriso *et al*., 2017).

In this context, species assessment studies of photosynthetic species are fundamental for the understanding of the wellbeing and stability of an ecosystem. Firstly, the reduction or the disappearance of a native species can be an early warning signal of the worsening of environmental conditions of an ecosystem (e.g., loss of ecosystem functions). Secondly, the appearance of a Non-Indigenous Species (NIS) requires targeted studies to assess the potential impacts of the organism. NIS pose a dual threat: they can lead to the extinction of native, vulnerable species, while also disrupting entire ecosystems by altering biogeochemical cycles and trophic chains, with consequential impacts on human well-being (Katsanevakis *et al*., 2014; Gallardo *et al*., 2016; Flood *et al*., 2020; Cuthbert *et al*., 2021; Sfriso *et al*., 2023).

These aspects are of high importance in delicate ecosystems as Transitional Water Systems (TWS), which are highly vulnerable to global climate change and human activities that can further induce biodiversity loss (Cuthbert *et al*., 2021). A primary example of this is the Venice Lagoon, which supports a high diversity of photosynthetic species but is also recognized as a major hotspot for biological invasions in the Mediterranean (Drigo *et al*., 2024). The genus *Sporobolus* provides a particularly relevant example. This genus is represented by the native species *S. maritimus* (Curtis) P.M. Peterson & Saarela, the NIS *S. anglicus* (C.E. Hubb.) P.M. Peterson & Saarela (Cazzin *et al*., 2009) and there are dubious reports of another NIS, *S. x townsendii* (H. Groves & J. Groves) P.M. Peterson & Saarela (Wong *et al*., 2018). Due to their complex evolutionary history, morphological traits alone are often insufficient for reliable identification, particularly when distinguishing *S. anglicus* from *S. x townsendii*. Molecular markers, such as *trnL*–*trnF*, can therefore be combined with genome-size estimates. This is especially informative because *S. anglicus* has approximately twice the genome size of the related taxa considered here (Chelaifa *et al*., 2010).

Within this framework, we assessed the diversity and distribution of photosynthetic species across the southern, central, and northern sectors of the Venice Lagoon. Morphological identification was integrated with molecular analyses to improve taxonomic confidence and evaluate the occurrence of non-indigenous species.

A targeted survey was also conducted to clarify the distribution of *Sporobolus* species, whose identification remains uncertain because of their morphological similarity (Wong *et al*., 2018). Since morphology alone cannot reliably distinguish these taxa, DNA barcoding was combined with nuclear genome-size analysis.

## 2. Materials and Methods

### 2.1 Sample collection and species identification

The Venice Lagoon is connected to the Adriatic Sea through three main inlets: Chioggia; Malamocco; Lido, arranged from the south to the north. The location and width of these inlets shape water circulation, allowing the lagoon to be divided into three main sectors: Southern, Central, and Northern.

During 2023 a preliminary sampling campaign was conducted in the Southern and Central parts of the lagoon on salt marshes and wood poles for species sampling and identification. Sampled sites are indicated in Figure 1 and latitude and longitude are reported in Supplementary Table 1. Collected organisms were brought to the laboratory for preliminary taxonomic identification.

**Figure 1.**
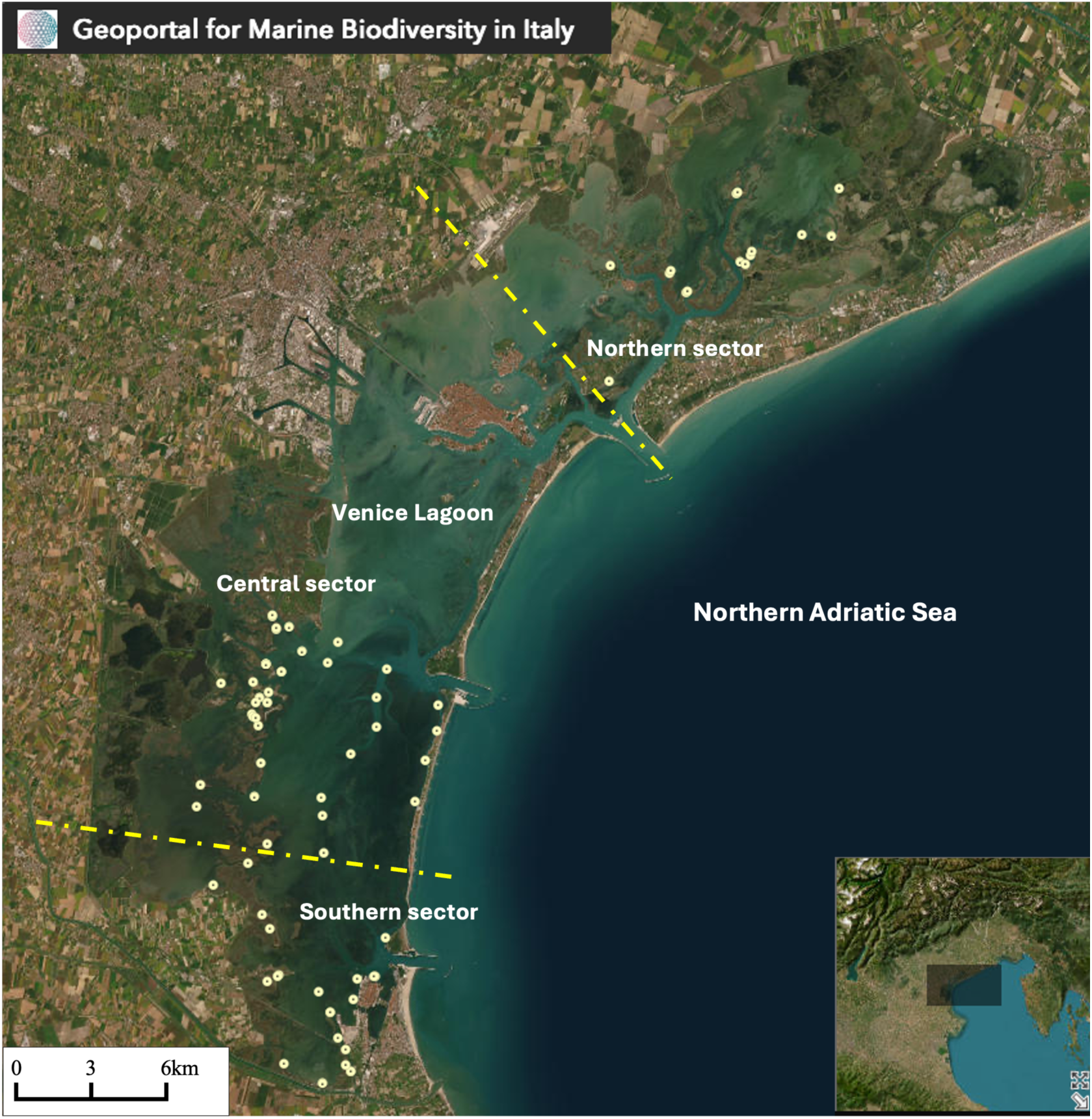
Sampling sites monitored during the three field campaigns. Dots indicate the wooden-pole and salt marsh sites surveyed in 2023 and 2024, as well as the sites included in the 2025 Sporobolus campaign. Dashed lines separate the Southern, Central, and Northern sectors of the Venice Lagoon. The map is available through the National Biodiversity Future Center’s Geoportal for Marine Biodiversity in Italy, hosted on the Biodiversity Gateway (https://www.biodiversitygateway.it/soluzioni-per-la-biodiversita/geoportale-biodiversita-marina-specie-habitat/). Sampling years, site labels, and geographic coordinates are provided in Supplementary Tables 1-3.

In 2024, a more comprehensive monitoring campaign was conducted, with equal numbers of salt marshes and wooden poles sampled in each lagoon sector (Southern, Central, and Northern) during spring, summer, and autumn. Sampled sites are indicated in Figure 1 and latitude and longitude are reported in Supplementary Table 2. Species assessment was performed along 30m lines, oriented parallel to the shoreline and located at increasing distances from the shore. High plant species presence was recorded along each line, and specimens requiring further confirmation were collected for identification in the laboratory. Wooden poles were sampled by scuba divers along their entire submerged length. Collected seaweeds were brought to the laboratory for preliminary taxonomic identification.

In the laboratory, all specimens were initially identified using taxonomic keys based on morphological traits (Pignatti, 1982; Wolf & Moro, 2013). Specimens of uncertain identity were preserved in plastic sterile bags containing silica gel for subsequent genomic DNA (gDNA) extraction. Selected samples were also fixed in 1.5 mL tubes containing a 5% glutaraldehyde in 0.1 M sodium cacodylate buffer (pH 6.9) and stored for subsequent cytological examination (Moro *et al*. 2020).

To characterize the biodiversity of the *Sporobolus* genus, during July 2025 we sampled an equal number of salt marshes for each sector (Figure 1 and coordinates are reported in Supplementary Table 3). To characterize the diversity of the genus *Sporobolus*, plants were transferred to the laboratory, carefully cleaned to remove salt marsh sediment, and potted in perlite. The pots were maintained in a growth chamber (POL-EKO, Poland) under a 16:8 h light/dark photoperiod, with a light intensity of 250 µmol photons m□² s□¹ and day/night temperatures of 25/23 °C. Each pot was irrigated daily with 100 mL of quarter-strength Hoagland’s solution (Hoagland & Arnon, 1950). This step was fundamental to correctly evaluate the genome size of collected specimen, since the analysis must be performed on fresh material (Baumel *et al*., 2003).

### 2.2 Sample preparation for light microscopic examination

After the fixation, samples were washed 2-4 times with 0.1 M sodium cacodylate buffer (pH 6.9) to remove the excess of glutaraldehyde. Subsequently, samples were postfixed in 1% OsO_4_ for 2 h and dehydrated in a graded series of ethyl alcohol and propylene oxide and embedded in araldite according to Moro *et al*. (2020). Thin sections (1 μm thick) were cut with an Ultracut Reichert-Jung ultramicrotome, stained with basic toluidine blue (1% toluidine blue and 1% Na tetraborate, 1:1 by volume), washed with distilled water, dried, and observed through a DMR Leica microscope (Wetzlar, Germany) equipped with a digital image acquisition system.

### 2.3 Genomic DNA extraction

Genomic DNA (gDNA) was extracted using different protocols for high plants and macroalgae. For each sample, 20 mg of dried biomass was frozen in liquid nitrogen and homogenized for 30 s at 35 Hz using a TissueLyser LT (Qiagen, Germany).

For vascular plant, including *Sporobolus* specimens, gDNA was extracted using the DNeasy Plant Pro Kit (Qiagen, Germany) according to the manufacturer’s instruction. Macroalgal gDNA was extracted using either the FastPure Plant DNA Isolation Mini Kit (Vazyme, China) or, when DNA quality was low, a CTAB-based protocol (unpublished protocol). For all methods, gDNA was eluted in nuclease-free water, and its concentration and purity were assessed using a NanoDrop UV spectrophotometer (Thermo Fisher Scientific, USA), while integrity was evaluated by electrophoresis on a 0.7% agarose gel. Samples were subsequently stored at -20 °C.

### 2.4 Amplification, Purification and Sequencing of gDNA markers

Organism identification based on molecular markers was carried out using the DNA barcoding technique. Different molecular markers were amplified for each specimen according to its taxonomic group: (1) the *ribulose-1,5-bisphosphate carboxylase/oxygenase large subunit (rbc*L) for Tracheophyta; (2) the *5’-region of the cytochrome c oxidase subunit I gene* (*COI-5P*) for Rhodophyta and Heterokontophyta; (3) the *plastid elongation factor Tu gene* (*tuf*A) for Chlorophyta; (4) intergenic spacer region *trnL-trnF* for *Sporobolus* species collected during the 2025 campaign. For selected genera, a second molecular marker was used to resolve taxonomic uncertainties: (1) *rbc*L for *Scytosiphon, Chondria, Vaucheria* and *Centroceras*; (2) maturase K (*mat*K) for *Atriplex*; *18S* ribosomal RNA gene (*18S* rRNA) for *Chaetomorpha*; (3) universal plastid amplicon (*UPA*) for *Antithamnion*; (4) internal transcribed spacer region (*ITS* rRNA) for *Limonium* and *Thinopyrum*; (5) intergenic spacer region *psbA-trnH* for *Salicornia*.

PCR primers and thermal profiles used for each marker are reported in Supplementary Table 4. In particular, primers targeting the *trnL-trnF* marker in *Sporobolus* species were designed using Primer-BLAST (Ye *et al*., 2012).

PCRs were performed in a final volume of 50 µL using 10 ng of gDNA, the appropriate reaction buffer, 1 µL of dNTP mix (10 mM each), 1 µL of each primer (10 µM), and 0.25 µL of Taq DNA polymerase (Promega, USA, or Vazyme, China). Amplifications were conducted using Mastercycler X40 (Eppendorf) or Applied Biosystems Veriti 96-Well and 2720 thermal cyclers. PCR products were verified by electrophoresis on 1% agarose gels, purified using the QIAquick PCR Purification Kit (Qiagen) and quantified by NanoDrop spectrophotometry. The purified amplicons were subsequently sequenced by Eurofins Genomics using the Mix2Seq service and the corresponding forward primer. For subsequent analysis, sequences were manually checked to remove low-quality portions.

### 2.5 Sequence Alignment and Species Delimitation Procedures

After quality assessment, sequences were preliminarily identified through similarity searches using BLASTn. Taxonomic assignments were further supported by phylogenetic analyses performed with the command-line version of IQ-TREE (Minh *et al*., 2020; Nguyen *et al*., 2015; Trifinopolus *et al*., 2016; Wong *et al*., 2026) with default settings. Branch support was assessed using three methods: (1) Ultrafast Bootstrap (UFBoot) with 1000 alignments (Minh *et al*., 2013; Hoang *et al*., 2018); (2) SH-aLRT (Shimodaira-Hasegawa-like approximate likelihood ratio test) branch test with 1000 replicates; (3) the Approximate Bayes test. Maximum-likelihood (ML) phylogenies were inferred using the best-fitting evolutionary model selected for each dataset by ModelFinder (Kalyaanamoorthy *et al*., 2017), which is reported in Supplementary Table 5 for each tree. Species boundaries were further evaluated using two independent delimitation methods: Assemble Species by Automatic Partitioning (ASAP) and the Generalized Mixed Yule–Coalescent (GMYC) model.

To apply these methods, sequences of closely related species and outgroups were downloaded from GenBank and iBOLD databases. Gene marker and accession number are reported in Supplementary Table 6. These sequences, along with those obtained from the Venice Lagoon, were aligned using MEGA desktop app (v.12) (Kumar *et al*., 2024) with the MUSCLE algorithm with default setting. ASAP analysis was performed using the desktop application with the Kimura two-parameter (K2P/K80) substitution model, while all other settings were left at their default values. For the GMYC analysis, the input file for BEAST v.2.7.7 was prepared using BEAUti v.2. The sequence alignment was imported, and the Hasegawa–Kishino–Yano model with gamma-distributed rate variation among sites (HKY+G) was selected Michonneau (2017). The substitution model was set to HKY, with four gamma-rate categories. Posterior distributions were estimated in BEAST using Markov chain Monte Carlo (MCMC) analyses with 10 million generations, sampling every 1,000 generations. Convergence and statistical reliability were assessed in Tracer v.1.7.2, requiring effective sample size (ESS) values above 200; otherwise, the MCMC settings were adjusted. Posterior trees were summarized using TreeAnnotator v.2.7.7, and GMYC analyses were performed in R v.4.5.1 following Michonneau (2017). Phylogenetic trees were visualized and annotated using iTOL (Letunic & Bork, 2024).

For Limonium ITS sequences, the secondary structure of the ITS2 region was additionally evaluated using the ITS2 Database (https://its2.bioapps.biozentrum.uni-wuerzburg.de).

### 2.6 Genome size comparison of *Sporobolus* species

Genome size was estimated by flow cytometry using approximately 100 mg of fresh leaf tissue per analysis. Based on the molecular marker identity, a leaf pool was prepared for each sampling site for uncertain specimens using a previously identified *S. anglicus* specimen as reference. All sample preparation was performed at 4 °C. Each pool was placed in a Petri dish on ice and supplemented with 1 mL of cold nuclei-extraction buffer containing 0.1 M Tris-HCl, 2 mM MgCl, 0.1 M NaCl, and 0.5% Triton X-100 at pH 7 (Baumel *et al*., 2003). The tissue was finely chopped with sterile scalpels, and the resulting suspension was filtered through a 50 µm CellTrics^®^ filter (Sysmex Partec, Germany) into a 1.5 mL tube maintained on ice. RNase A was added to a final concentration of 50 µg/mL, followed by incubation at room temperature for 1 h. Propidium iodide (PI) was then added to a final concentration of 50 µg/mL, and the samples were incubated on ice in the dark for 15 min.

Flow-cytometric analyses were performed using a CytoFLEX cytometer and CytExpert software (Beckman Coulter, USA). Samples were excited at 488 nm, and PI fluorescence was detected using a 610/20 nm filter. Events were initially gated based on PI fluorescence and side scatter, after which fluorescence intensity was evaluated using PI histograms. Data acquisition continued until 100,000 events had been recorded per sample. The same leaves used for gDNA extraction and PCR amplification were used for genome size comparison.

## 3. Results

### 3.1 Large-scale monitoring of photosynthetic species diversity: 2023–2024 surveys

Species surveys conducted in salt marshes and on wooden poles focused on halophytes, seagrasses, and seaweeds. During the 2023 and 2024 campaigns, photosynthetic organisms were identified using an integrated morphological and molecular approach. Of the 347 gDNA samples analysed, 197 (∼52%) produced high-quality DNA sequences that could be used for BLAST comparisons and for species delimitation with the ASAP and GMYC algorithms and phylogenetic analysis. Only thirteen sequences (∼5% of the total) were classified at the genus level, while the majority (206, ∼95% of the total) were classified at the species level. DNA barcoding results were inconsistent with the initial morphological identification for 44.64% of macroalgal samples and 29.41% of vascular plant samples. A total of 81 taxa were identified, including 20 at the genus level and 61 at the species level, as listed in Table 1, with detailed distribution reported in Supplementary Table 7. We identified 13 Chlorophyta (Figure 2), 11 Heterokontophyta (1 Xantophyceae and 10 Phaeophyceae) (Figure 3 and Figure 4), 28 Rhodophyta (Figure 5) and 29 Tracheophyta (Figure 6).

**Figure 2.**
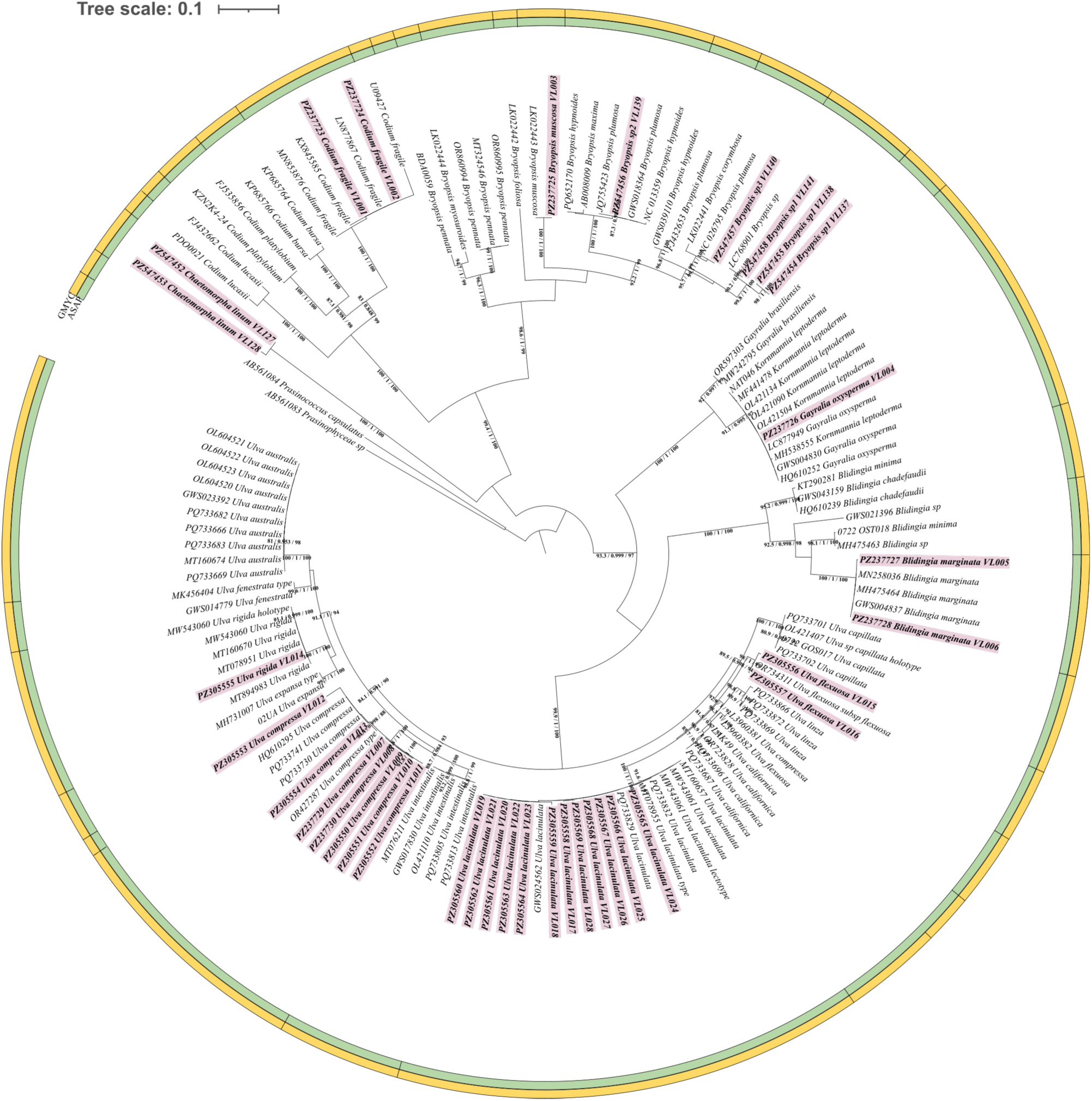
Phylogenetic reconstruction of Chlorophyta from the Venice Lagoon based on *tuf*A sequences. The tree was inferred using IQ-TREE. Node values indicate SH-aLRT/aBayes/UFBoot support, with SH-aLRT values shown only when ≥80%. ASAP and GMYC delimitation results are represented by green and yellow outer circles, respectively. Survey samples are shown in bold and highlighted in light magenta, while GenBank sequences are labelled with accession numbers and species names (Supplementary Table 6).

**Figure 3.**
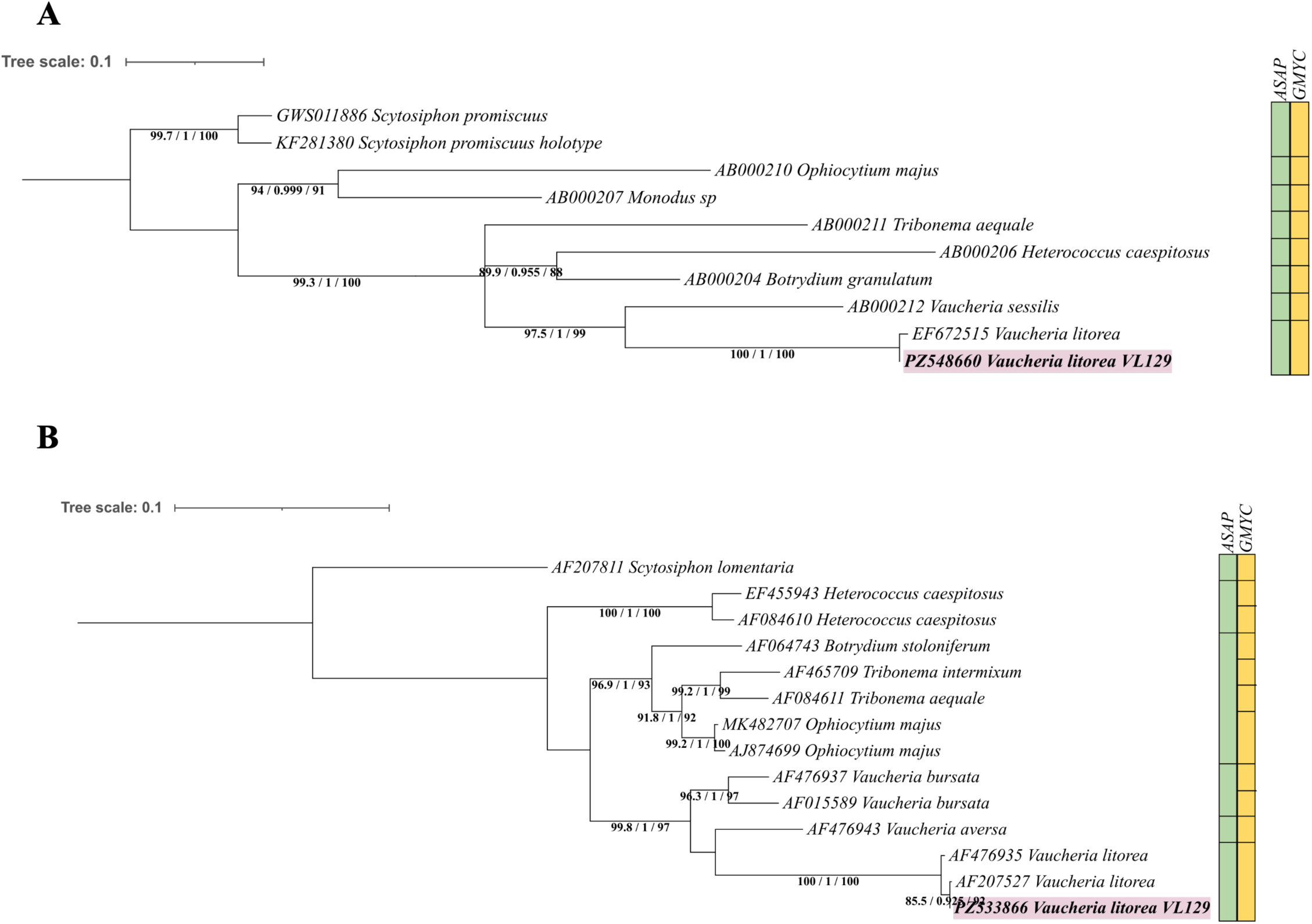
Phylogenetic reconstruction of Xanthophyceae from the Venice Lagoon based on *COI-5P* (A) and *rbc*L (B) sequences. Trees were inferred using IQ-TREE. Node values indicate SH-aLRT/aBayes/UFBoot support, with SH-aLRT values shown only when ≥80%. ASAP and GMYC species-delimitation results are represented by green and yellow boxes, respectively. Survey samples are shown in bold and highlighted in light magenta, while GenBank sequences are labelled with accession numbers and species names (Supplementary Table 6).

**Figure 4.**
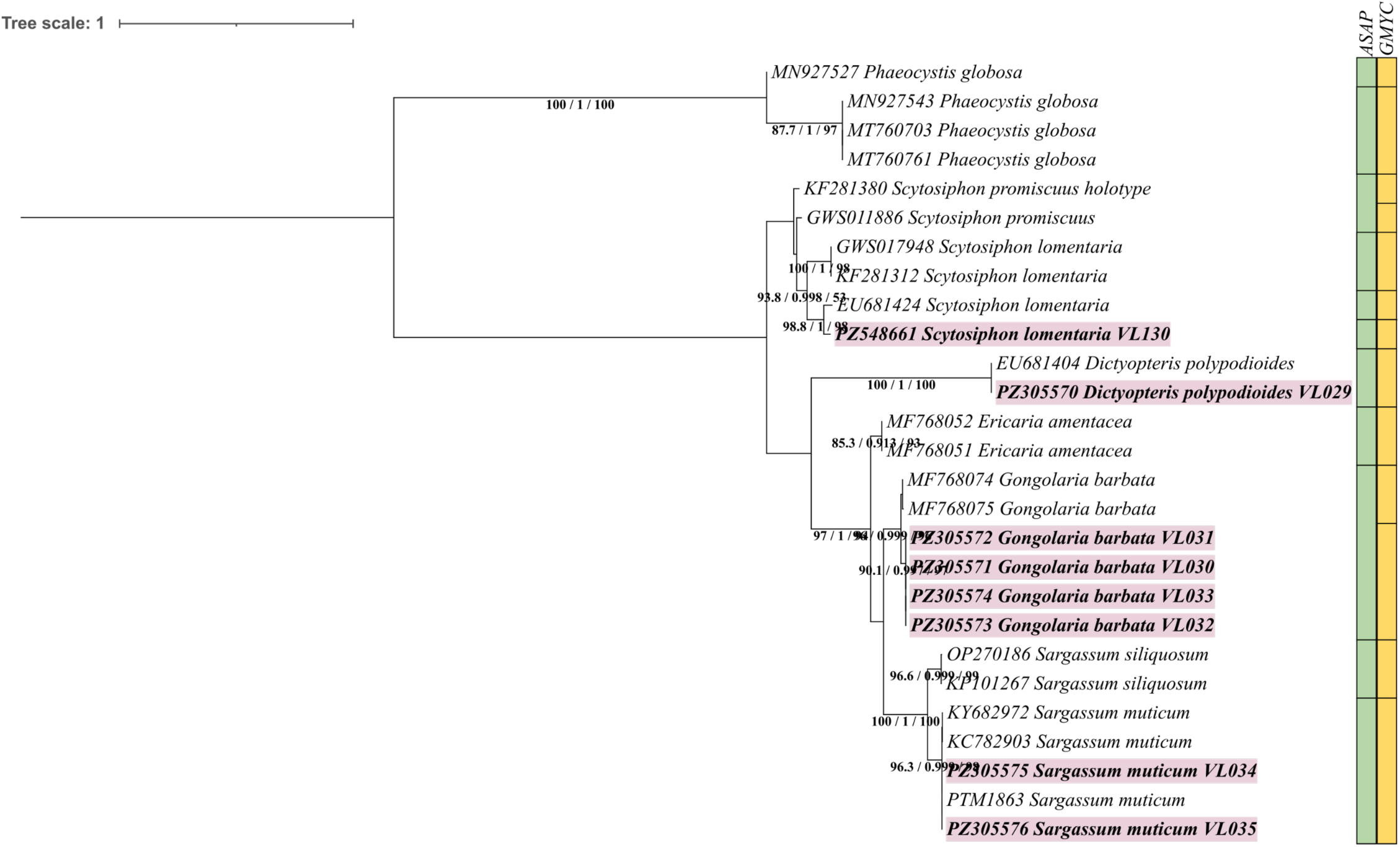
Phylogenetic reconstruction of Phaeophyceae from the Venice Lagoon based on *COI-5P* sequences. The tree was inferred using IQ-TREE. Node values indicate SH-aLRT/aBayes/UFBoot support, with SH-aLRT values shown only when ≥80%. ASAP and GMYC species-delimitation results are represented by green and yellow boxes, respectively. Survey samples are shown in bold and highlighted in light magenta, while GenBank sequences are labelled with accession numbers and species names (Supplementary Table 6).

**Figure 5.**
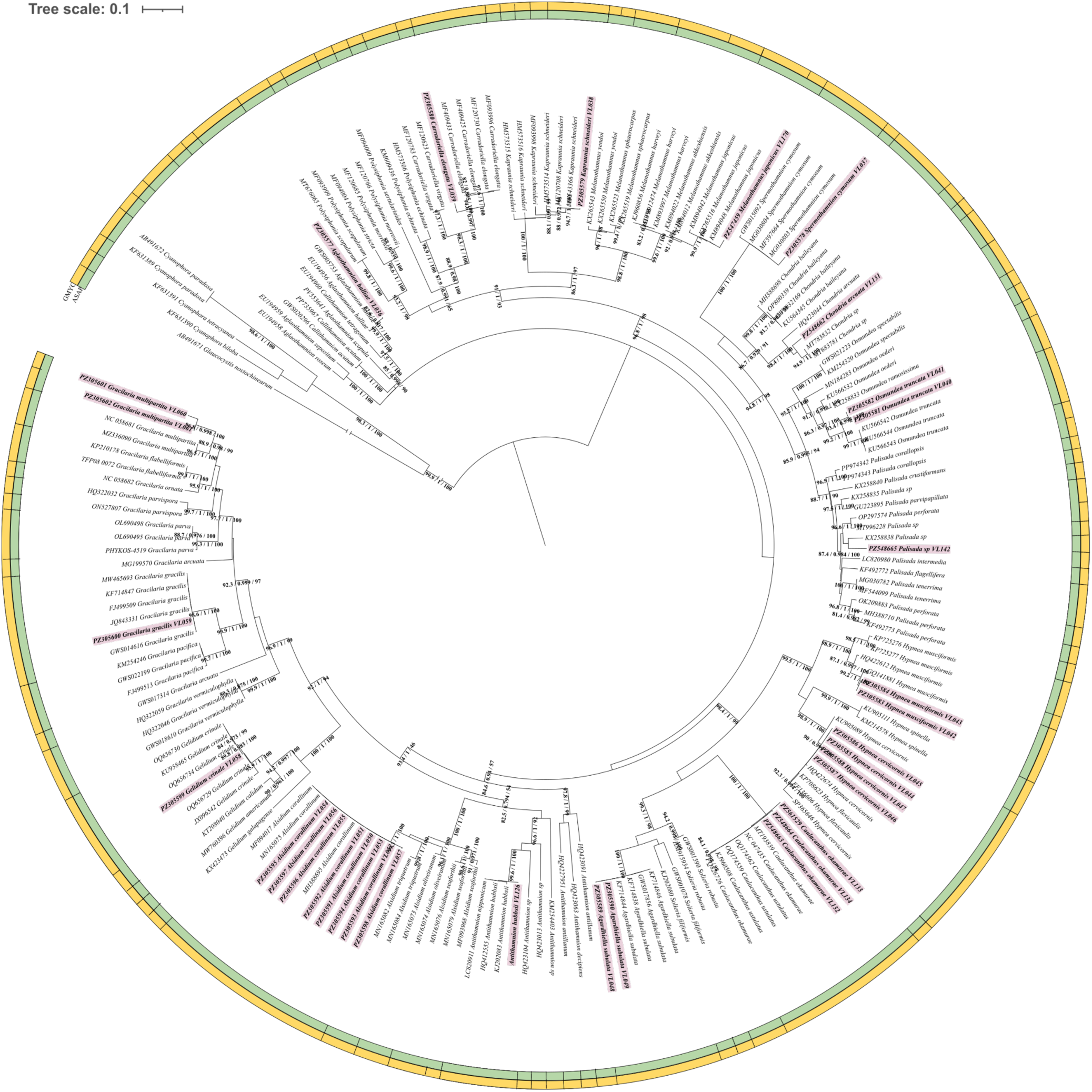
Phylogenetic reconstruction of Rhodophyta from the Venice Lagoon based on *COI-5P* sequences. The tree was inferred using IQ-TREE. Node values indicate SH-aLRT/aBayes/UFBoot support, with SH-aLRT values shown only when ≥80%. ASAP and GMYC delimitation results are represented by green and yellow outer circles, respectively. Survey samples are shown in bold and highlighted in light magenta, while GenBank sequences are labelled with accession numbers and species name (Supplementary Table 6).

**Figure 6.**
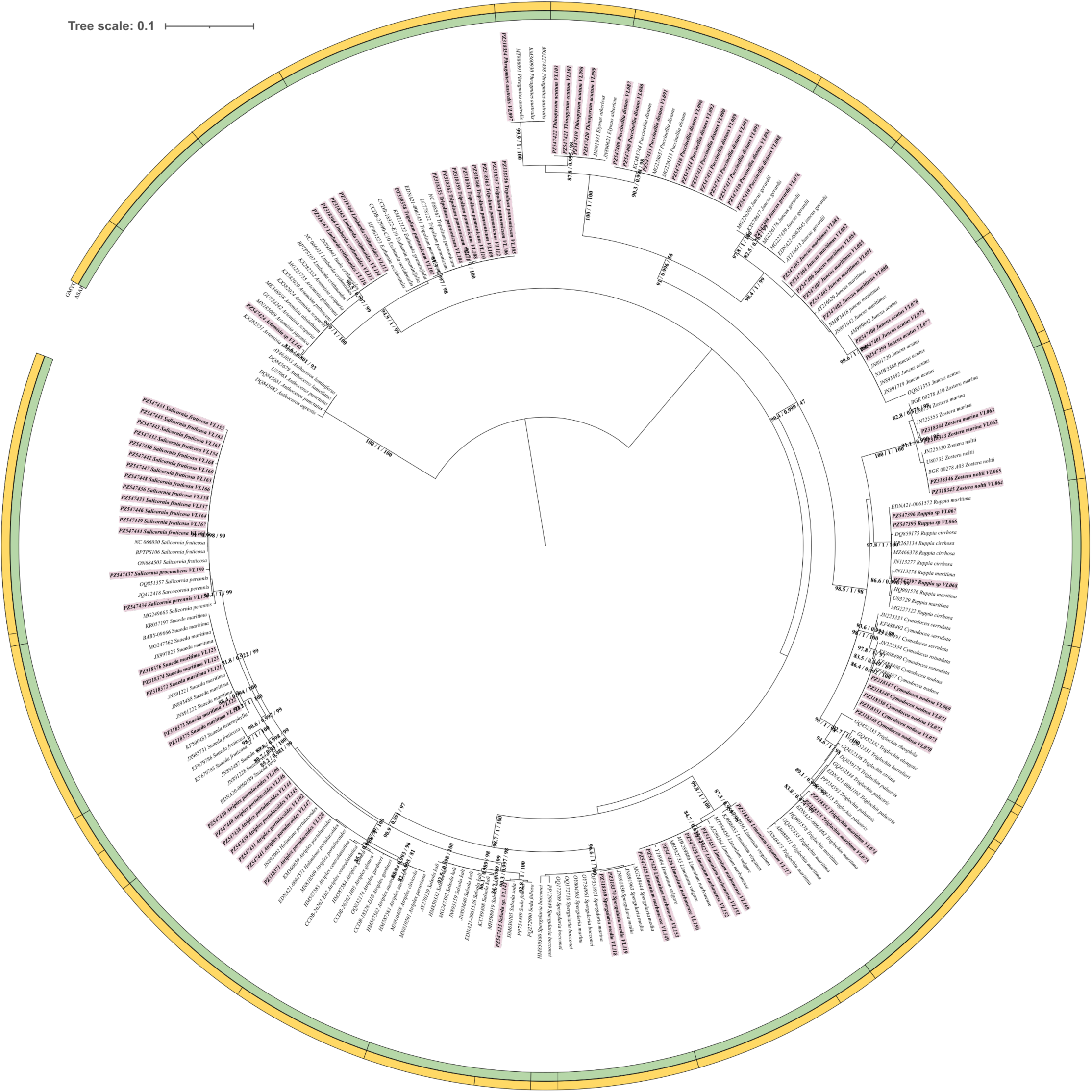
Phylogenetic reconstruction of Tracheophyta from the Venice Lagoon based on *rbc*L sequences. The tree was inferred using IQ-TREE. Node values indicate SH-aLRT/aBayes/UFBoot support, with SH-aLRT values shown only when ≥80%. ASAP and GMYC delimitation results are represented by green and yellow outer circles, respectively. Survey samples are shown in bold and highlighted in light magenta, while GenBank sequences are labelled with accession numbers and species names (Supplementary Table 6).

**Table 1.** Species identified in the Venice Lagoon. For each species, the group (C = Chlorophyta, H = Halopytes, P = Phaeophyceae, R = Rhodophyta, S = Seagrasses, X = Xantophyceae), the identification method and the corresponding field campaign are reported.

| Species | Group | Morphology | Molecular analysis | 2023 | 2024 | 2025 |
| --- | --- | --- | --- | --- | --- | --- |
| *Agardhiella subulata* | R |  | x |  | x |  |
| *Aglaothamnion halliae* | R |  | x |  | x |  |
| *Aglaothamnion sp.* | R | x |  |  | x |  |
| *Alsidium corallinum* | R |  | x |  | x |  |
| *Antithamnion hubbsii* | R |  | x |  | x |  |
| *Antithamnion sp.* | R | x |  |  | x |  |
| *Artemisia sp.* | H |  | x |  | x |  |
| *Atriplex portulacoides* | H |  | x | x | x |  |
| *Blidingia marginata* | C |  | x |  | x |  |
| *Bryopsis muscosa* | C |  | x |  | x |  |
| *Bryopsis sp.* | C | x | x | x | x |  |
| *Carradoriella elongata* | R |  | x | x |  |  |
| *Caulacanthus okamurae* | R |  | x |  | x |  |
| *Centroceras gasparrinii* | R |  | x |  | x |  |
| *Chaetomorpha linum* | C |  | x |  | x |  |
| *Chaetomorpha sp.* | C | x |  | x |  |  |
| *Chondria arcuata* | R |  | x | x |  |  |
| *Chondria sp.* | R | x |  |  | x |  |
| *Cladophora sp.* | C | x |  |  | x |  |
| *Codium fragile* | C |  | x |  | x |  |
| *Cymodocea nodosa* | S |  | x | x | x |  |
| *Cystoseira sp.* | P | x |  |  | x |  |
| *Dasya sp.* | P | x |  |  | x |  |
| *Dictyopteris polypodioides* | P | x | x |  | x |  |
| *Dictyota dichotoma* | P | x |  |  | x |  |
| *Dictyota fasciola* | P | x |  |  | x |  |
| *Dictyota sp.* | P | x |  | x |  |  |
| *Ectocarpus siliculosus* | P | x |  |  | x |  |
| *Fucus virsoides* | P | x |  | x | x |  |
| *Gayralia oxysperma* | C |  | x |  | x |  |
| *Gelidium crinale* | R |  | x | x |  |  |
| *Gongolaria barbata* | P | x | x | x | x |  |
| *Gracilaria gracilis* | R |  | x |  | x |  |
| *Gracilaria multipartita* | R |  | x |  | x |  |
| *Heterosiphonia sp.* | R | x |  |  | x |  |
| *Hypnea cervicornis* | R |  | x |  | x |  |
| *Hypnea musciformis* | R |  | x |  | x |  |
| *Juncus acutus* | H |  | x |  | x |  |
| *Juncus gerardii* | H |  | x |  | x |  |
| *Juncus maritimus* | H |  | x |  | x |  |
| *Kapraunia schneideri* | R | x | x |  | x |  |
| *Limbarda crithmoides* | H | x | x | x | x |  |
| *Limonium sp.* | H | x |  |  | x |  |
| *Limonium narbonense* | H |  | x | x | x |  |
| *Limonium virgatum* | H |  | x |  | x |  |
| *Melanothamnus japonicus* | R |  | x |  | x |  |
| *Nitophyllum punctatum* | R | x |  |  | x |  |
| *Osmundea truncata* | R |  | x | x | x |  |
| *Palisada sp.* | R | x | x |  | x |  |
| *Phragmites australis* | H |  | x |  | x |  |
| *Phyllophora sicula* | R | x |  |  | x |  |
| *Polysiphonia sp.* | R | x |  |  | x |  |
| *Puccinellia distans* | H | x | x |  | x |  |
| *Pyropia sp.* | R |  | x |  | x |  |
| *Radicilingua sp.* | R | x |  |  | x |  |
| *Ruppia sp.* | S |  | x |  | x |  |
| *Salicornia perennis* | H |  | x |  | x |  |
| *Salicornia procumbens* | H |  | x |  | x |  |
| *Salicornia fruticosa* | H |  | x | x | x |  |
| *Salicornia sp.* | H | x |  |  | x |  |
| *Salsola sp.* | H |  | x |  | x |  |
| *Sargassum muticum* | P |  | x |  | x |  |
| *Scytosiphon lomentaria* | P |  | x |  | x |  |
| *Spergularia media* | H |  | x | x | x |  |
| *Spermothamnion cymosum* | R |  | x |  | x |  |
| *Sporobolus sp.* | H | x | x | x | x |  |
| *Sporobolus anglicus* | H |  | x |  |  | x |
| *Sporobolus maritimus* | H |  | x |  |  | x |
| *Sporobolus x townsendii* | H |  | x |  |  | x |
| *Suaeda maritima* | H |  | x |  | x |  |
| *Thinopyrum acutum* | H |  | x |  | x |  |
| *Triglochin maritima* | H |  | x |  | x |  |
| *Tripolium pannonicum* | H |  | x |  | x |  |
| *Ulva compressa* | C |  | x | x | x |  |
| *Ulva lacinulata* | C |  | x | x | x |  |
| *Ulva rigida* | C |  | x |  | x |  |
| *Ulva flexuosa* | C |  | x | x | x |  |
| *Ulva sp.* | C | x |  | x | x |  |
| *Vaucheria litorea* | X |  | x | x |  |  |
| *Zostera marina* | S | x | x | x | x |  |
| *Zostera noltii* | S |  | x | x | x |  |

For samples that we could not identify at the species level with one marker, a second molecular marker was amplified to improve taxonomic resolution. The resulting phylogenetic reconstructions and species-delimitation analyses are presented in Supplementary Figure 1 for Chlorophyta (*18S* rRNA) and Rhodophyta (*UPA* and *rbc*L), and in Supplementary Figure 2 for Tracheophyta (*ITS* rRNA, *mat*K, and *psbA-trnH*).

Among the species recorded in 2023, *Vaucheria litorea*, *Gelidium crinale*, *Chondria arcuata* and *Carradoriella elongata* were not detected during the 2024 campaign. No specimens of *Ulva australis* were detected (Figure 2).

Only one representative of the Xantophyceae phylum was recorded, *Vaucheria litorea*. Whilst two canopy-forming brown algal species (Heterokontophyta) were identified (Figure 4). *Fucus virsoides* was recorded in the Southern sector during spring and in the Central sector during summer. *Gongolaria barbata* occurred in the Central sector during spring and summer and in the Northern sector during autumn. Additional specimens assigned to *Cystoseira* sp. were found in the southern and Central sectors during spring.

Regarding seagrasses, *Cymodocea nodosa* and *Zostera marina* were found in all three lagoon sectors across all seasons. In contrast, *Zostera noltii* was recorded in every sector during spring and only in the Northern sector during autumn, while it was not detected during summer. As shown in Figure 6, we identified samples of the genus *Ruppia*, however a definitive classification in *R. maritima* or *R. cirrhosa* could not be completed. Nevertheless, this genus is less common, occurring exclusively in the Central sector during summer and in the Southern sector during autumn. For terrestrial plants, most species were recorded in the Southern sector of the lagoon, while the fewest were found in the Northern sector. Several species were exclusively detected in the Southern sector, including *Artemisia sp.*, *Phragmites australis*, *Salsola sp.*, *Thinopyrum acutum* and *Triglochin maritima*.

Overall, the integration of the morphological and molecular approach clarified some cases allowing the identification at the species level (e.g., *Antithamnion, Chaetomorpha, Chondria, Gracilaria, Osmundea, Sargassum, Scytosiphon, Ulva*), while others were still not clear (e.g., *Artemisia, Palisada, Ruppia*). A total of 81 species were identified, including 9 of the 35 non-indigenous species previously reported in the Venice Lagoon, the canopy-forming brown algae *Fucus virsoides* and *Gongolaria barbata*, and all seagrass species known to occur in the lagoon. All sequences generated in this study were deposited in GenBank; accession numbers are provided in Supplementary Table 8.

### 3.2 Occurrence of *Sporobolus* species in the Venice Lagoon

To clarify the occurrence of *S. x townsendii* in the Venice Lagoon, we integrated morphological observations, DNA barcoding, and genome-size measurements. A total of 105 plants were collected: 34 from the Southern sector, 31 from the Central sector and 40 from the Northern sector. The specimens recorded at each sampling site (about 5 per site) are listed in Table 2. Because of their evolutionary history, *S. anglicus* and *S. x townsendii* share the same chloroplast *trnL-trnF* marker. However, *S. x townsendii* has a genome size comparable to that of *S. maritimus* and approximately half that of *S. anglicus*. These characteristics allow us to distinguish among *S. maritimus*, *S. x townsendii*, and *S. anglicus*. Combining DNA barcoding with flow-cytometric genome-size measurements therefore enabled the three species to be distinguished: *S. maritimus* was characterized by the *S. maritimus* chloroplast haplotype and the smaller genome size; *S. x townsendii* by the *S. alterniflorus/S. anglicus* chloroplast haplotype and the smaller genome size; and *S. anglicus* by the same chloroplast haplotype but an approximately doubled genome size. A schematic overview of this approach is provided in Supplementary Figure 3, while the combined barcoding and flow-cytometry results are presented in Table 2 and detailed in Supplementary Table 9. Using this integrated approach, *S. x townsendii* was detected in the Southern sector at site LS 5 and in the Central sector at sites LC 7, LC 8, and LC 11, but not in the Northern sector. Two of these salt marshes were artificial (LS 5 and LC 7), whereas LC 8 and LC 11 were natural. *S. anglicus* was recorded at all surveyed sites, while the native *S. maritimus* was not detected at three sites: LS 1, LC 7, and LC 10. These findings confirm the occurrence of *S. x townsendii* in the Venice Lagoon, particularly in its Southern and Central sectors. They also reveal the widespread presence of the non-indigenous *S. anglicus* and the local absence of *S. maritimus* from three sampling sites. All sequences generated in this study were deposited in GenBank, and their accession numbers are provided in Supplementary Table 8. Representative herbarium specimens were also deposited in the Botanical Garden collection of the University of Padua (Supplementary Figure 4 to 12).

**Table 2.** Monitoring sites and identification of Sporobolus species. . The table reports site ID (LS, southern lagoon; LC, central lagoon; LN, northern lagoon), geographic coordinates (latitude and longitude), and number of *S. maritimus* and *S. anglicus* specimens initially identified based on morphology. Final taxonomic assignments were obtained by integrating DNA barcoding with flow-cytometric genome-size analysis (see Supplementary Figure 3 for details).

| Site ID | Latitude (°N) | Longitude (°E) | Morphologically identified as *S. maritimus* | Morphologically identified as *S. anglicus* | Final taxonomic assignments |  |  |
| --- | --- | --- | --- | --- | --- | --- | --- |
|  |  |  |  |  | *S. maritimus* | *S. anglicus* | *S. x townsendii* |
| LS 1 | 45.185389° | 12.253889° | - | 2 | - | 2 | - |
| LS 2 | 45.2238° | 12.2244° | 3 | 3 | 3 | 3 | - |
| LS 3 | 45.2258° | 12.2301° | 3 | 3 | - | 6 | - |
| LS 4 | 45.2488° | 12.2219° | 2 | 5 | 2 | 5 | - |
| LS 5 | 45.2683° | 12.2144° | 3 | 3 | 2 | 3 | 1 |
| LS 6 | 45.2602° | 12.1955° | 3 | 4 | 3 | 4 | - |
| LC 7 | 45.2894° | 12.1872° | - | 3 | - | 1 | 2 |
| LC 8 | 45.3228° | 12.2187° | 4 | 3 | 2 | 3 | 1 |
| LC 9 | 45.32866° | 12.22448° | 2 | 3 | 2 | 3 | - |
| LC10 | 45.342933° | 12.223733° | - | 3 | - | 3 | - |
| LC 11 | 45.3613° | 12.2275° | 3 | 4 | 2 | 4 | 1 |
| LC 12 | 45.3569° | 12.2363° | 3 | 3 | 2 | 4 | - |
| LN 13 | 45.4494° | 12.4072° | 2 | 4 | 2 | 4 | - |
| LN 14 | 45.4925° | 12.4077° | 3 | 4 | 3 | 4 | - |
| LN 15 | 45.490806° | 12.439889° | 3 | 2 | 3 | 2 | - |
| LN 16 | 45.483139° | 12.449194° | 3 | 3 | 3 | 3 | - |
| LN 17 | 45.492972° | 12.479917° | 4 | 6 | 4 | 6 | - |
| LN 18 | 45.520167° | 12.475611° | 3 | 3 | 3 | 3 | - |

## 4. Discussions

### 4.1 Monitoring of photosynthetic species diversity

Accurate species identification is essential for assessing biodiversity patterns and detecting changes in community composition over time. This is particularly challenging for photosynthetic organisms, whose identification may be hindered by phenotypic plasticity, simple morphology, cryptic diversity, and morphologically distinct life stages.

In this study, integrating morphological observations with DNA barcoding and species-delimitation analyses improved taxonomic resolution. Morphology guided the selection of suitable molecular markers and helped exclude biologically implausible sequence matches, while comparisons with GenBank and delimitation algorithms supported the identification of uncertain specimens. However, only slightly more than half of the samples yielded high-quality sequences, emphasizing the need to optimize gDNA extraction protocols for seaweeds and improve the primers available for marker amplification.

The identification of *Osmundea truncata* illustrates the value of an integrative approach. GenBank similarity searches could not reliably distinguish the samples from *O. oederi* and *O. pinnatifida*, two other species reported in the Venice Lagoon. This uncertainty was particularly relevant because *O. oederi* is non-indigenous (Sfriso *et al*., 2023). Phylogenetic and species-delimitation analyses consistently grouped the samples with *O. truncata*, clearly separating them from the other two taxa. Similarly, several sequences showed high similarity to multiple *Alsidium* species. Since only *A. corallinum* has previously been reported in the Venice Lagoon (Drigo *et al*., 2024), resolving their identity was important for detecting potential new introductions. Species-delimitation analyses consistently grouped all samples with *A. corallinum*, providing no evidence of additional *Alsidium* species in the surveyed sites. Another taxonomically challenging case involved samples showing similarly high sequence identity to *Caulacanthus okamurae* and *C. ustulatus*, differing by only one percentage point. *C. okamurae* is a non-indigenous species first recorded in the Venice Lagoon in the 21^st^ century (Sfriso *et al*., 2023; Drigo *et al*., 2024), whereas *C. ustulatus*, considered invasive in Europe, has been reported in the lagoon since the 19^th^ century (Roy *et al*., 2020; Drigo *et al*., 2024). The initial marker, phylogenetic reconstruction, and species-delimitation analyses did not resolve the ambiguity. Amplification of a second marker, *rbc*L, identified the samples as *C. okamurae*, supporting its presence in the Venice Lagoon (Supplementary Figure 1B).

Within Chlorophyta, several samples morphologically assigned to *Bryopsis* showed similar BLAST matches to multiple congeneric species. Phylogenetic and species-delimitation analyses resolved only sample VL003, which clustered with *B. muscosa* with maximum support (100/1/100; Figure 2). The remaining samples could be identified only as *Bryopsis* sp., reflecting the complex, often polyphyletic relationships reported within the genus (Leliaert & Lopez-Bautista, 2015). Another sequence showed high similarity to both *Gayralia oxysperma* (Ulotrichales order) and *Kornmannia leptoderma* (Ulvales order). Phylogenetic and delimitation analyses placed it within the *Gayralia* clade, close to *G. oxysperma* and clearly separated from Ulvales order. These results support its identification as *G. oxysperma* and suggest that the matching *K. leptoderma* GenBank sequences may require taxonomic reassessment.

In the case of higher plants, DNA barcoding did not provide unambiguous identification for samples of the genus *Triglochin*. Sequence alignment with GenBank returned high similarity scores with multiple species, particularly *T. palustris* and *T. maritima*, both previously reported in the Venice Lagoon during the 19^th^ and 20^th^ centuries, respectively (Drigo *et al*., 2024). Phylogenetic reconstruction and species-delimitation analyses resolved the taxonomic uncertainty, identifying the specimen as *T. maritima* and supporting the continued occurrence of this species in the Venice Lagoon during the 21^st^ century (Drigo *et al*., 2024). Together, these examples demonstrate the value of integrative taxonomy for accurately identifying seaweeds and vascular plants and improving the reliability of biodiversity monitoring (Magoga *et al*., 2021). As mentioned before, *Ruppia maritima* and *R. cirrhosa* in our dataset are not separated. However, their classification is not homogeneous in reference databases. In fact, according to POWO (2026), these two species are separated, but according to AlgaeBase (Giury M.D. and Giury G.M, 2026) and Ito and co-authors (2017), *R. cirrhosa* is a synonym of *R. maritima*. Thus, it is not possible to classify our species other than *Ruppia* sp.

Another dubious case is represented by *Salsola sp.* VL 143. The *rbc*L marker is not able to confidentially separate this sample from *Salsola soda*, the native species of the Venice Lagoon (Sfriso *et al*., 2009; Drigo *et al*., 2024), and *Soda foliosa*, distributed mainly in Central Asia (POWO, 2026) and never reported in the venetian lagoon. Despite the amplification of a second marker (*mat*K, Supplementary Figure 2B), the lack of reference sequences of *Soda foliosa* from GenBank did not allow us to identify the sample at the species level. Additional surveys and independent taxonomic validation are therefore required.

Seaweed diversity was dominated by Rhodophyta, followed by Chlorophyta and Heterokontophyta, whereas no Charophyta were detected. Since charophytes are primarily associated with freshwater and low-salinity habitats (Pukacz *et al*., 2024), their absence may reflect the historical reduction in freshwater inputs caused by human modifications to the Venice Lagoon (Drigo *et al*., 2024). These changes may have limited suitable habitats even in the less impacted Northern sector. The lower richness of Phaeophyceae relative to Rhodophyta may also suggest a decline in ecological quality, as several brown algal taxa are commonly associated with relatively unpolluted waters (Marzocchi *et al*., 2001). However, this interpretation should also consider differences in habitat availability, seasonality, and sampling effort. The non-detection of *Ulva australis* contrasts with recent surveys reporting its widespread occurrence in the Venice Lagoon and a concurrent decline in the native *U. rigida* (Sfriso *et al*., 2020, 2023). Conversely, numerous specimens were identified as *U. lacinulata*, a species not formally reported in the lagoon since the 19^th^ century (Drigo *et al*., 2024). This apparent discrepancy may reflect the complex taxonomy of *Ulva*, whose species often share highly similar morphological traits. Until recently, *U. lacinulata* was considered a synonym of *U. rigida* (Hughey *et al*., 2022); consequently, specimens collected during the 20^th^ century may have been assigned to *U. rigida*. These findings emphasize the importance of continuous monitoring supported by integrated morphological and molecular identification, while recognizing that the non-detection of a species during a survey does not necessarily demonstrate its absence from the lagoon. Related to new species introductions in the Venice lagoon, an interesting case is represented by *Vaucheria litorea*. Previous works reported the presence of other species of the same genus in the Venice lagoon (Sfriso *et al*., 2009; Drigo *et al*., 2024) and according to AlgaeBase (Guiry M. D. and Guiry G. M., 2026), *V. litorea* was not reported in Italy. The specimen was identified using two molecular markers, *COI-5P* and *rbc*L. However, morphological identification was not possible because it lacked the reproductive structures required for reliable anatomical classification (Brown, 1929; Entwisle, 1987). The species was collected in 2023 but not detected during the 2024 survey, suggesting that its occurrence may be localized or intermittent. Continued monitoring of the sites where it was initially recorded is therefore needed to clarify the distribution and persistence of *Vaucheria* in the Venice Lagoon, where the genus represents one of the few members of Xanthophyceae.

In terms of species that provide ecosystem services, some macroalgal species are considered habitat-forming, such as the case of *Gongolaria barbata* and *Fucus virsoides* (Bilajac *et al*., 2024; Descourvières *et al*., 2024; Pica *et al*., 2024; Gljušćić *et al*., 2025). However, marine heatwaves have been shown to endanger *G. barbata* survival (Bilajac *et al*., 2024) and *F. virsoides* declined significantly in the last decades (Gljušćić *et al*., 2025). Here, we recorded both species in the Southern and Central sectors of the lagoon, but no records of *F. virsoides* occurred in the Northern sector, whilst *G. barbata* was present. Unfortunately, the causes behind the decline of *F. virsoides* populations in recent decades remain unclear (Descourvières *et al*., 2024), making it difficult to explain its absence in this area of the lagoon. As the only endemic representative of the genus *Fucus* in the Mediterranean, understanding its eco-physiological responses is crucial to prevent its potential disappearance from the Venice Lagoon. These studies should consider the effects of marine heatwaves, which will likely increase in frequency in the future (Garrabou *et al*., 2022) and could greatly affect the Venice lagoon due to its shallow depth. The increase of heatwave frequency and intensity is a threat also for *G. barbata*, which is still widespread in the whole lagoon, but recent studies have shown an increase in mortality after exposure to temperatures higher than 34 °C (Gljušćić *et al*., 2025).

Another important component of the photosynthetic biodiversity is represented by seagrasses. These higher plants colonize soft substrata and reduce sediment resuspension through the *mat*, their intricate root system that compact and stabilize the soil (Macreadie *et al*., 2017; Sfriso *et al*., 2017), providing oxygen and nutrient to the ecosystem (Macreadie *et al*., 2017). This study confirmed the presence of the four seagrass species previously reported in the Venice Lagoon, with *Cymodocea nodosa* and *Zostera marina* being the most frequent, consistent with earlier monitoring (Rismondo *et al*., 2003). The rarest species belongs to the genus *Ruppia* which have been found only in the Southern and Central sectors, mainly in ponds inside salt marshes. Previous studies (Sfriso *et al*., 2009) reported a lower diffusion of *Ruppia* species, with a partial recovery in recent years, especially in the Northern sector (Sfriso *et al*., 2022). Continuous monitoring should be carried out to better assess actual diffusion of the *Ruppia* species in the lagoon.

In addition, also halophytes are crucial for the stability and preservation of salt marshes, which are primarily composed of medium-to-fine sediments (Taramelli *et al*., 2021). Similarly to seagrasses, their root systems help compact and stabilize the sediment (De Battisti *et al*., 2019). In terms of species richness, the highest number of halophyte species was recorded in the Southern sector, likely due to the proximity of these salt marshes to terrestrial areas. This might have favoured the arrival and establishment of seeds from terrestrial areas, since some of the sampled salt marshes are artificial and show higher elevation compared to others, creating areas of soil with lower salt content, that might have favoured their survival.

Most halophyte species occurred across all three sectors of the lagoon. Historical records document several *Limonium* species, although only *L. narbonense* had been reported in the 21^st^ century (Drigo *et al*., 2024) and was also detected in our survey. DNA barcoding additionally indicated the presence of *L. virgatum*. Within the genus *Juncus*, *J. maritimus* occurred mainly in the Central and Southern sectors. *Juncus acutus* and *J. gerardii* were also recorded in the Central sector, although neither had previously been reported in the lagoon during the 21^st^ century (Drigo *et al*., 2024).

Lastly, of the 35 non-indigenous species (NIS) previously reported in the Venice Lagoon (Sfriso *et al*., 2023; Drigo *et al*., 2024), the following were identified in this study: *Agardhiella subulata, Aglaothamnion halliae, Antithamnion hubbsii, Caulcanthus okamurae, Codium fragile, Hypnea cervicornis, Melanothamnus japonicus, Sargassum muticum* and *Spermothamnion cymosum*. The occurrence of NIS showed distinct spatial and seasonal patterns (Supplementary Table 7). *A. subulata* was recorded only in the Southern sector during spring, whereas *A. halliae* occurred in the Southern sector in spring and the Central sector in autumn. *A. hubbsii* was recorded in the Southern sector during autumn. *C. okamurae* occurred in the Central and Northern sectors in both spring and autumn, while *C. fragile* was detected in these sectors only during summer. *H. cervicornis* was found in the Southern and Central sectors during summer and in the Northern sector during autumn. *M. japonicus* occurred in the Southern sector, and *S. muticum* only in the Central sector during spring.

Overall, integrating morphological and molecular approaches improved taxonomic resolution and helped resolve several ambiguous identifications.

The surveys also provided the first records of *V. litorea* (Xantophyceae) for the Venice Lagoon. Further targeted monitoring and independent taxonomic validation are needed to confirm its occurrences and assess its persistence. Continued biodiversity surveys will also be essential for evaluating ecological change and ecosystem stability in this Mediterranean biodiversity hotspot.

### 4.2 Occurrence of *Sporobolus x townsendii* in the Venice lagoon

The first records of non-native *Sporobolus* taxa in the Venice Lagoon date to the early 2000s, when *S. x townsendii* was initially reported (Scarton *et al*., 2003; Ghirelli, 2004). Subsequent studies suggested that these records might instead represent *S. anglicus* (Cazzin *et al*., 2009; Wong *et al*., 2018), leaving the occurrence of *S. x townsendii* unresolved in this area.

The present study confirms the current presence of *S. x townsendii* in the Venice Lagoon. The morphological traits distinguishing it from related taxa were subtle and difficult to assess in the field. Reliable identification was therefore achieved by combining DNA barcoding with flow-cytometric genome-size measurements, demonstrating the importance of an integrative approach for this species complex. Among the *Sporobolus* taxa examined, *S. x townsendii* showed the most restricted distribution across the surveyed sites. A possible explanation is related to its reproductive behaviour. Being sterile (Ainouche *et al*., 2004; Ainouche *et al*., 2012), its growth and diffusion are vegetative, limiting its propagation in the environment. The spread of *S. x townsendii* may be further constrained by competition with *S. anglicus*, a strong competitor for resources (Ainouche *et al*., 2004). More importantly, the widespread occurrence of *S. anglicus* may threaten the persistence of the native *S. maritimus*. During our survey, *S. anglicus* was detected at every sampling site, whereas *S. maritimus* was not recorded at one site in the Southern sector (LS 1) and two sites in the Central sector (LC 7 and LC 10). Although non-detection does not demonstrate local absence, this pattern warrants continued monitoring. *S. anglicus* is reported to be a vigorous and aggressive perennial plant (Ainouche *et al*., 2004), widespread in other parts of continental Europe and UK (Baumel *et al*., 2001) and worldwide (Morgan & Systma, 2010; Maebara *et al*., 2020). Additionally, there are reports of ecosystem impacts in relation to *S. anglicus* spreading (Goss-Custard & Moser 1988). These characteristics make *S. anglicus* a NIS of the highest concern for native organisms. This pattern is particularly concerning in the Venice Lagoon, where anthropogenic pressures strongly affect biological communities. Further studies should quantify the relative abundance and distribution of *S. maritimus* and *S. anglicus*, particularly as increasingly frequent extreme climatic events may favour stress-tolerant taxa, such as *S. anglicus* (unpublished results).

## 5. Conclusions

This study shows that integrating morphological, molecular, phylogenetic, and cytometric methods improves the identification of photosynthetic species in the Venice Lagoon.

The survey confirmed the lagoon’s considerable diversity of seaweeds, seagrasses, and halophytes despite the strong pressures this environment is exposed to. The first record of *Vaucheria litorea* require further validation and monitoring. Particular attention should also be given to vulnerable habitat-forming species such as *Fucus virsoides* and *Gongolaria barbata*. The combined use of DNA barcoding and genome-size analysis confirmed the occurrence of *Sporobolus x townsendii*. This taxon showed a restricted distribution, whereas the non-indigenous *S. anglicus* occurred at every surveyed site. The native *S. maritimus* was not detected at three sites, raising concerns about its persistence. Overall, this study provides an updated taxonomic inventory and new reference sequences. These resources represent a valuable baseline for future biodiversity assessments. Long-term integrated monitoring will be essential for detecting new introductions and tracking ecological changes. Such efforts will support the conservation and sustainable management of the Venice Lagoon.

## Acknowledgments

We thank Cristina Breggion, Andrea Sambo and Vittoria Correale of the Chioggia Hydrobiological Station (Department of Biology, University of Padua, Italy - https://chioggia.biologia.unipd.it/stazione-idrobiologica/) for their support with the fieldwork.

## Author contributions

Conceptualization: FD, DDB, IM, CDP

Data Curation: FD

Formal Analysis: FD, DDB, CS, RT, CF

Funding Acquisition: CDP

Investigation: FD, RT, CF, CS, EM, DDB

Methodology: FD, RT, CS, DDB

Project Administration: CDP

Resources: CF, IM, CDP

Supervision: DDB, IM, CDP

Validation: IM, CDP

Visualization: FD, IM, CDP

Writing - Original Draft: FD

Writing - Review & Editing: FD, RT, CF, CS, EM, DDB, CDP, IM

## Data Availability

Molecular marker sequences generated in this study are available in GenBank (Supplementary Table 8), while species distribution data are accessible through the National Biodiversity Future Center (NBFC) Geoportal for Marine Biodiversity in Italy (https://www.biodiversitygateway.it/soluzioni-per-la-biodiversita/geoportale-biodiversita-marina-specie-habitat/).

## Supplementary material – brief legends

**Supplementary Figure 1. Phylogenetic reconstruction based on secondary markers of Chlorophyta and Rhodophyta from the Venice Lagoon.**

**Supplementary Figure 2. Phylogenetic reconstruction based on secondary markers of Tracheophyta from the Venice Lagoon.**

**Supplementary Figure 3. Conceptual workflow used to identify *Sporobolus* species in the Venice Lagoon.**

**Supplementary Figure 4. Herbarium specimen LN_17_9 of *Sporobolus* collected during the 2025 survey.**

**Supplementary Figure 5. Herbarium specimen LC_11_7 of *Sporobolus* collected during the 2025 survey.**

**Supplementary Figure 6. Herbarium specimen of LC_7_2 *Sporobolus* collected during the 2025 survey.**

**Supplementary Figure 7. Herbarium specimen LC_7_3 *of Sporobolus* collected during the 2025 survey.**

**Supplementary Figure 8. Herbarium specimen LC_7_1 of *Sporobolus* collected during the 2025 survey.**

**Supplementary Figure 9. Herbarium specimen LC_12_2 of *Sporobolus* collected during the 2025 survey.**

**Supplementary Figure 10. Herbarium specimen LC_12_4 of *Sporobolus* collected during the 2025 survey.**

**Supplementary Figure 11. Herbarium specimen LC_8_6 *and* LC_8_7 of *Sporobolus* collected during the 2025 survey.**

**Supplementary Figure 12. Herbarium specimen LS_5_4 of *Sporobolus* collected during the 2025 survey.**

**Supplementary Table 1. Sites sampled during the 2023 survey.**

**Supplementary Table 2. Sites sampled during the 2024 survey.**

**Supplementary Table 3. Sites sampled during the 2025 survey of *Sporobolus* species.**

**Supplementary Table 4. List of primers and PCR amplification profiles used for molecular marker amplification.**

**Supplementary Table 5. Best-fit model determined by ModelFinder.**

**Supplementary Table 6. GenBank accession numbers of reference sequences used for phylogenetic reconstruction.**

**Supplementary Table 7. Species identified during the 2023 and 2024 surveys and their distribution in the Venice Lagoon.**

**Supplementary Table 8. GenBank accession numbers of sequences generated during the monitoring surveys.**

**Supplementary Table 9. Molecular and flow-cytometric characterization of the collected *Sporobolus* specimens.**

## References

Ainouche, M.L., Baumel, A., Salmon, A., Yannic, G., 2004. Hybridization, polyploidy and speciation in Spartina (Poaceae). New Phytologist 161, 165–172. 10.1046/j.1469-8137.2003.00926.x

Ainouche, M., Chelaifa, H., Ferreira, J., Bellot, S., Ainouche, A., Salmon, A. 2012. Erratum From - Polyploid Evolution in Spartina: Dealing with Highly Redundant Hybrid Genomes. In: Soltis, P., Soltis, D. (eds) Polyploidy and Genome Evolution. Springer, Berlin, Heidelberg. 10.1007/978-3-642-31442-1_12

Arterburn, M., Kleinhofs, A., Murray, T., Jones, S., 2011. Polymorphic nuclear gene sequences indicate a novel genome donor in the polyploid genus Thinopyrum: Polymorphic sequences indicate a novel ancestor of Thinopyrum. Hereditas 148, 8–27. 10.1111/j.1601-5223.2010.02084.x

Bartolo, A.G., Zammit, G., Peters, A.F., Küpper, F.C., 2020. The current state of DNA barcoding of macroalgae in the Mediterranean Sea: presently lacking but urgently required. Botanica Marina 63, 253–272. 10.1515/bot-2019-0041

Baumel, A., Ainouche, M.L., Levasseur, J.E., 2001. Molecular investigations in populations of Spartina anglica C.E. Hubbard (Poaceae) invading coastal Brittany (France). Molecular Ecology 10, 1689–1701. 10.1046/j.1365-294X.2001.01299.x

Baumel, A., Ainouche, M.L., Misset, M.T., Gourret, J.-P., Bayer, R.J., 2003. Genetic evidence for hybridization between the native Spartina maritima and the introduced Spartina alterniflora (Poaceae) in South-West France: Spartina × neyrautii re-examined. Plant Systematics and Evolution 237, 87–97. 10.1007/s00606-002-0251-8

Bilajac, A., Gljušćić, E., Smith, S., Najdek, M., Iveša, L., 2024. Effects of extreme temperatures and recovery potential of Gongolaria barbata from a coastal lagoon in the northern Adriatic Sea: an ex situ approach. Ann Bot 134, 415–426. 10.1093/aob/mcae038

Brown, H.J., 1929. The Algal Family Vaucheriaceae. Transactions of the American Microscopical Society 48, 86. 10.2307/3222465

Bruni, I., De Mattia, F., Martellos, S., Galimberti, A., Savadori, P., Casiraghi, M., Nimis, P.L., Labra, M., 2012. DNA Barcoding as an Effective Tool in Improving a Digital Plant Identification System: A Case Study for the Area of Mt. Valerio, Trieste (NE Italy). PLoS ONE 7, e43256. 10.1371/journal.pone.0043256

Burgess, K.S., Fazekas, A.J., Kesanakurti, P.R., Graham, S.W., Husband, B.C., Newmaster, S.G., Percy, D.M., Hajibabaei, M., Barrett, S.C.H., 2011. Discriminating plant species in a local temperate flora using the *rbcL* + *matK* DNA barcode. Methods Ecol Evol 2, 333–340. 10.1111/j.2041-210X.2011.00092.x

Cazzin, M., Ghirelli, L., Mion, D., and Scarton, F. 2009. Completamento della cartografia della vegetazione e degli habitat della laguna di Venezia: anni 2005-2007. Lavori Soc. Ven. Sc. Nat. 34, 81–89.

Chelaifa, H., Monnier, A., Ainouche, M., 2010. Transcriptomic changes following recent natural hybridization and allopolyploidy in the salt marsh species Spartina × townsendii and Spartina anglica (Poaceae). New Phytologist 186, 161–174. 10.1111/j.1469-8137.2010.03179.x

Cuthbert, R.N., Pattison, Z., Taylor, N.G., Verbrugge, L., Diagne, C., Ahmed, D.A., Leroy, B., Angulo, E., Briski, E., Capinha, C., Catford, J.A., Dalu, T., Essl, F., Gozlan, R.E., Haubrock, P.J., Kourantidou, M., Kramer, A.M., Renault, D., Wasserman, R.J., Courchamp, F., 2021. Global economic costs of aquatic invasive alien species. Science of The Total Environment 775, 145238. 10.1016/j.scitotenv.2021.145238

De Battisti, D., Fowler, M.S., Jenkins, S.R., Skov, M.W., Rossi, M., Bouma, T.J., Neyland, P.J., Griffin, J.N., 2019. Intraspecific Root Trait Variability Along Environmental Gradients Affects Salt Marsh Resistance to Lateral Erosion. Front. Ecol. Evol. 7. 10.3389/fevo.2019.00150

De Castro, O., Di Maio, A., Lozada García, J.A., Piacenti, D., Vázquez-Torres, M., De Luca, P., 2013. Plastid DNA sequencing and nuclear SNP genotyping help resolve the puzzle of central American Platanus. Annals of Botany 112, 589–602. 10.1093/aob/mct134

De Vere, N., Rich, T.C.G., Trinder, S.A., Long, C., 2015. DNA Barcoding for Plants, in: Batley, J. (Ed.), Plant Genotyping, Methods in Molecular Biology. Springer New York, New York, NY, pp. 101–118. 10.1007/978-1-4939-1966-6_8

Descourvières, E., Mulas, M., Natale, S., de Pedro, R.S., Alboresi, A., Solidoro, C., Bandelj, V., Falace, A., 2024. Ecophysiological responses of Fucus virsoides (Phaeophyceae, Fucales) to past and present nutrient conditions in the northern Adriatic. Mar Biol 171, 205. 10.1007/s00227-024-04523-1

Drigo, F., Stefanelli, C., Moro, I., De Pittà, C., 2024. Evaluation of photosynthetic Taxa in the Venice Lagoon from the nineteenth century to present day. Front. Mar. Sci. 11, 1433680. 10.3389/fmars.2024.1433680

Elias, M., Hill, R.I., Willmott, K.R., Dasmahapatra, K.K., Brower, A.V.Z., Mallet, J., Jiggins, C.D., 2007. Limited performance of DNA barcoding in a diverse community of tropical butterflies. Proc. R. Soc. B. 274, 2881–2889. 10.1098/rspb.2007.1035

Entwisle, T.J., 1987. An evaluation of taxonomic characters in the subsection Sessiles, section Corniculatae, of Vaucheria (Vaucheriaceae, Chrysophyta). Phycologia 26, 297–231. 10.2216/i0031-8884-26-3-297.1

Flood, P.J., Duran, A., Barton, M., Mercado-Molina, A.E., Trexler, J.C., 2020. Invasion impacts on functions and services of aquatic ecosystems. Hydrobiologia 847, 1571– 1586. 10.1007/s10750-020-04211-3

Freshwater, D.W., Rueness, J., 1994. Phylogenetic relationships of some European Gelidium (Gelidiales, Rhodophyta) species, based on *rbcL* nucleotide sequence analysis. Phycologia 33, 187–194. 10.2216/i0031-8884-33-3-187.1

Gallardo, B., Clavero, M., Sánchez, M.I., Vilà, M., 2016. Global ecological impacts of invasive species in aquatic ecosystems. Global Change Biology 22, 151–163. 10.1111/gcb.13004

Garrabou, J., Gómez Gras, D., Medrano, A., Cerrano, C., Ponti, M., Schlegel, R., Bensoussan, N., Turicchia, E., Sini, M., Gerovasileiou, V., Teixido, N., Mirasole, A., Tamburello, L., Cebrian, E., Rilov, G., Ledoux, J., Souissi, J.B., Khamassi, F., Ghanem, R., Benabdi, M., Grimes, S., Ocaña, O., Bazairi, H., Hereu, B., Linares, C., Kersting, D.K., La Rovira, G., Ortega, J., Casals, D., Pagès Escolà, M., Margarit, N., Capdevila, P., Verdura, J., Ramos, A., Izquierdo, A., Barbera, C., Rubio Portillo, E., Anton, I., López Sendino, P., Díaz, D., Vázquez Luis, M., Duarte, C., Marbà, N., Aspillaga, E., Espinosa, F., Grech, D., Guala, I., Azzurro, E., Farina, S., Cristina Gambi, M., Chimienti, G., Montefalcone, M., Azzola, A., Mantas, T.P., Fraschetti, S., Ceccherelli, G., Kipson, S., Bakran Petricioli, T., Petricioli, D., Jimenez, C., Katsanevakis, S., Kizilkaya, I.T., Kizilkaya, Z., Sartoretto, S., Elodie, R., Ruitton, S., Comeau, S., Gattuso, J., Harmelin, J., 2022. Marine heatwaves drive recurrent mass mortalities in the Mediterranean Sea. Global Change Biology 28, 5708–5725. 10.1111/gcb.16301

Gavio, B., Fredericq, S., 2002. Grateloupia turuturu (Halymeniaceae, Rhodophyta) is the correct name of the non-native species in the Atlantic known as Grateloupia doryphora. Euro. J. Phycol. 37, 349–359. 10.1017/S0967026202003839

Ghirelli L., 2004. Spartina x townsendii H. et J. Groves. In: Segnalazioni floristiche italiane: 1115. Informatore Botanico Italiano, 36 (1): 84–85.

Guiry, M. D., and Guiry, G. M. (2026). AlgaeBase (World-wide electronic publication, University of Galway). Available at: https://www.algaebase.org.

Gljušćić, E., Bilajac, A., Ferrari, V., Iveša, L., 2025. Effects of increased temperatures during air exposure on the morphology and physiology of Fucus virsoides (Fucales, Phaeophyceae). Algae 40, 241–255. 10.4490/algae.2025.40.7.1

Goss-Custard, J.D., Moser, M.E., 1988. Rates of Change in the Numbers of Dunlin, Calidris alpina, Wintering in British Estuaries in Relation to the Spread of Spartina anglica. The Journal of Applied Ecology 25, 95. 10.2307/2403612

Hoagland, D.R. and Arnon, D.I., 1950. The water-culture method of growing plants without soil. Calif. Agr. Expt. Sta. Circ. 347.

Hoang, D.T., Chernomor, O., Von Haeseler, A., Minh, B.Q., Vinh, L.S., 2018. UFBoot2: Improving the Ultrafast Bootstrap Approximation. Molecular Biology and Evolution 35, 518–522. 10.1093/molbev/msx281

Huang, B., Teng, L., Ding, L., 2016. Morphological and molecular discrimination of green macroalgae Chaetomorpha aerea and C. linum. Acta Oceanol. Sin. 35, 118–123. 10.1007/s13131-016-0841-x

Hughey, J.R., Gabrielson, P.W., 2022. DNA Analysis of the Lectotype Specimen of Ulva nematoidea Bory (Ulvaceae, Chlorophyta) Determines Its Synonymy with Ulva Lactuca L. crya 43, 117–124. 10.5252/cryptogamie-algologie2022v43a7

Iamonico, D., De Castro, O., Di Iorio, E., Nicolella, G., Iberite, M., 2022. Taxonomy Complexity of Some Tyrrhenian Endemic Limonium Species Belonging to L. multiforme Group (Plumbaginaceae): New Insights from Molecular and Morphometric Analyses. Plants 11, 3163. 10.3390/plants11223163

Ito, Y., Ohi-Toma, T., Nepi, C., Santangelo, A., Stinca, A., Tanaka, N., Murata, J., 2017. Towards a better understanding of the *Ruppia maritima* complex (Ruppiaceae): Notes on the correct application and typification of the names *R. cirrhosa* and *R. spiralis*. TAXON 66, 167–171. 10.12705/661.11

Kalyaanamoorthy, S., Minh, B.Q., Wong, T.K.F., von Haeseler, A., Jermiin, L.S., 2017. ModelFinder: fast model selection for accurate phylogenetic estimates. Nat Methods 14, 587–589. 10.1038/nmeth.4285

Kang, J.C., Kim, M.S., 2013. A novel species Symphyocladia glabra sp. nov. (Rhodomelaceae, Rhodophyta) from Korea based on morphological and molecular analyses. ALGAE 28, 149–160. 10.4490/algae.2013.28.2.149

Katsanevakis, S., Coll, M., Piroddi, C., Steenbeek, J., Ben Rais Lasram, F., Zenetos, A., Cardoso, A.C., 2014. Invading the Mediterranean Sea: biodiversity patterns shaped by human activities. Front. Mar. Sci. 1. 10.3389/fmars.2014.00032

Kogame, K., Horiguchi, T., Masuda, M., 1999. Phylogeny of the order Scytosiphonales (Phaeophyceae) based on DNA sequences of *rbc* L, partial *rbc* S, and partial LSU nrDNA. Phycologia 38, 496–502. 10.2216/i0031-8884-38-6-496.1

Kowalska, Z., Pniewski, F., Latała, A., 2019. DNA barcoding – A new device in phycologist’s toolbox. Ecohydrology & Hydrobiology 19, 417–427. 10.1016/j.ecohyd.2019.01.002

Kumar, S., Stecher, G., Suleski, M., Sanderford, M., Sharma, S., and Tamura, K. 2024. MEGA12: Molecular Evolutionary Genetic Analysis Version 12 for Adaptive and Green Computing. Molecular Biology and Evolution, 41: 1–9. 10.1093/molbev/msae263

Leliaert, F., Lopez-Bautista, J.M., 2015. The chloroplast genomes of Bryopsis plumosa and Tydemania expeditiones (Bryopsidales, Chlorophyta): compact genomes and genes of bacterial origin. BMC Genomics 16, 204. 10.1186/s12864-015-1418-3

Leliaert, F., Verbruggen, H., Vanormelingen, P., Steen, F., López-Bautista, J.M., Zuccarello, G.C., De Clerck, O., 2014. DNA-based species delimitation in algae. European Journal of Phycology 49, 179–196. 10.1080/09670262.2014.904524

Letunic, I., Bork, P., 2024. Interactive Tree of Life (iTOL) v6: recent updates to the phylogenetic tree display and annotation tool. Nucleic Acids Research 52, W78–W82. 10.1093/nar/gkae268

Macreadie, P.I., Jarvis, J., Trevathan Tackett, S.M., Bellgrove, A., 2017. Seagrasses and Macroalgae: Importance, Vulnerability and Impacts, in: Phillips, B.F., Pérez Ramírez, M. (Eds.), Climate Change Impacts on Fisheries and Aquaculture. Wiley, pp. 729–770. 10.1002/9781119154051.ch22

Maebara, Y., Tamaoki, M., Iguchi, Y., Nakahama, N., Hanai, T., Nishino, A., Hayasaka, D., 2020. Genetic Diversity of Invasive Spartina alterniflora Loisel. (Poaceae) Introduced Unintentionally Into Japan and Its Invasion Pathway. Front. Plant Sci. 11, 556039. 10.3389/fpls.2020.556039

Magoga, G., Fontaneto, D., Montagna, M., 2021. Factors affecting the efficiency of molecular species delimitation in a species rich insect family. Molecular Ecology Resources 21, 1475–1489. 10.1111/1755-0998.13352

Marzocchi, M., Bellemo, G., Miotti, C., Curiel, D., and Scattolin, M. 2001. Le macroalghe dei substrati duri del canal grande (centro storico di venezia): prime considerazioni. Boll. Mus. Civ. St. Nat. 52, 25–39.

Michonneau, F. ‘Using GMYC for Species Delineation’. Zenodo, 2 August 2017. 10.5281/zenodo.838260.

Minh, B.Q., Nguyen, M.A.T., Von Haeseler, A., 2013. Ultrafast Approximation for Phylogenetic Bootstrap. Molecular Biology and Evolution 30, 1188–1195. 10.1093/molbev/mst024

Minh, B.Q., Schmidt, H.A., Chernomor, O., Schrempf, D., Woodhams, M.D., Von Haeseler, A., Lanfear, R., 2020. IQ-TREE 2: New Models and Efficient Methods for Phylogenetic Inference in the Genomic Era. Molecular Biology and Evolution 37, 1530–1534. 10.1093/molbev/msaa015

Morgan, V. H., and Systma, M. 2010. Alaska Spartina Prevention, Detection and Response Plan (Juneau, AK: National Marine Fisheries Service Alaska Region).

Moro, I., Trentin, R., Moschin, E., Dalla Vecchia, F., 2020. Morpho-physiological responses by Isochrysis galbana Parke to different concentrations of oxytetracycline. Environmental Pollution 262, 114273. 10.1016/j.envpol.2020.114273

Nguyen, L.-T., Schmidt, H.A., Von Haeseler, A., Minh, B.Q., 2015. IQ-TREE: A Fast and Effective Stochastic Algorithm for Estimating Maximum-Likelihood Phylogenies. Molecular Biology and Evolution 32, 268–274. 10.1093/molbev/msu300

Pica, M.L., Vitale, E., Donadio, R., Costanzo, G., Munari, M., Fabbrizzi, E., Fraschetti, S., Arena, C., 2024. Functional ecological traits in young and adult thalli of canopy-forming brown macroalga *Gongolaria barbata* (Phaeophyta) from a transitional water system. PeerJ 12, e17959. 10.7717/peerj.17959

Pignatti, S. 1982. Flora d’Italia. Bologna: Edagricole POWO, 2026. Plants of the World Online (Facilitated by the Royal Botanic Gardens, Kew). Available at: http://www.plantsoftheworldonline.org/.

Pukacz, A., Asaeda, T., Blindow, I., Bueno, N.C., Casanova, M.T., Herbst, A., Pełechaty, M., Ranawaka, V., Schubert, H., 2024. Ecology of Charophytes: Niche Dimensions of Pioneers and Habitat Engineers, in: Schubert, H., Blindow, I., Nat, E., Korsch, H., Gregor, T., Denys, L., Stewart, N., van de Weyer, K., Romanov, R., Casanova, M.T. (Eds.), Charophytes of Europe. Springer Nature Switzerland, Cham, pp. 125–152. 10.1007/978-3-031-31898-6_7

Rismondo, A., Curiel, D., Scarton, F., Mion, D., Caniglia, G., 2003. A New Seagrass Map for the Venice Lagoon. Proceedings of the Sixth International Conference on the Mediterranean Coastal Environment, MEDCOAST 03. Vol 2: 843–852.

Robuchon, M., Valero, M., Gey, D., Le Gall, L., 2015. How does molecular-assisted identification affect our estimation of α, β and γ biodiversity? An example from understory red seaweeds (Rhodophyta) of Laminaria kelp forests in Brittany, France. Genetica 143, 207–223. 10.1007/s10709-014-9796-z

Roy, D., Alderman, D., Anastasiu, P., Arianoutsou, M., Augustin, S., Bacher, S., et al. 2020. DAISIE - Inventory of alien invasive species in Europe. Version 1.7 (Research Institute for Nature and Forest (INBO) (Accessed March 8, 2024).

Rybalka, N., Andersen, R.A., Kostikov, I., Mohr, K.I., Massalski, A., Olech, M., Friedl, T., 2009. Testing for endemism, genotypic diversity and species concepts in Antarctic terrestrial microalgae of the *Tribonemataceae* ( *Stramenopiles*, Xanthophyceae ). Environmental Microbiology 11, 554–565. 10.1111/j.1462-2920.2008.01787.x

Rybalka, N., Epkes, S., Wegner, K.M., Michaelis, R., Reise, K., 2022. Invasive *Vaucheria* (Xanthophyceae) at the lower shore of the Wadden Sea. Phycologia 61, 274–283. 10.1080/00318884.2022.2035532

Saunders, G.W., McDevit, D.C., 2012. Methods for DNA Barcoding Photosynthetic Protists Emphasizing the Macroalgae and Diatoms, in: Kress, W.J., Erickson, D.L. (Eds.), DNA Barcodes, Methods in Molecular Biology. Humana Press, Totowa, NJ, pp. 207–222. 10.1007/978-1-61779-591-6_10

Scarton, F., Ghirelli, L., Curiel, D., and Rismondo, A. 2003. First data on Spartina× townsendii in the lagoon of Venice (Italy). In Proceedings of the sixth international conference on the Mediterranean coastal environment, MEDCOAST, Ravenna, Italy, vol. 3, pp. 787–792.

Sfriso, A., Curiel, D., and Rismondo, A. 2009. “The Lagoon of Venice,” in Flora and Vegetation of the Italian Transitional Water Systems, 1st ed. Eds. E. Cecere, A. Petrocelli, G. Izzo and A. Sfriso (Stampa “Multigraf”, Spinea (VE), 17–80.

Sfriso, A., Buosi, A., Facca, C., and Sfriso, A. A. 2017. Role of environmental factors in affecting macrophyte dominance in transitional environments: The Italian Lagoons as a study case. Mar. Ecol. 38, e12414. doi: 10.1111/maec.12414

Sfriso, A., Buosi, A., Wolf, M. A., Sfriso, A. A. 2020. Invasion of alien macroalgae in the Venice Lagoon, a pest or a resource? Aquatic Invasion. 15, 245–270. doi: 10.3391/ai.2020.15.2.03

Sfriso, A., Wolf, M. A., Buosi, A., Sciuto, K., and Sfriso, A. A. 2023. Alien macroalgal rearrangement in the soft substrata of the Venice lagoon (Italy): impacts, threats, time and future trends. Sustainability 15, 8256. doi: 10.3390/su15108256

Sherwood, A.R., Presting, G.G., 2007. Universal primers amplify a 23S rDNA plastid marker in eukaryotic algae and cyanobacteria. Journal of Phycology 43, 605–608. 10.1111/j.1529-8817.2007.00341.x

Taramelli, A., Valentini, E., Piedelobo, L., Righini, M., Cappucci, S., 2021. Assessment of State Transition Dynamics of Coastal Wetlands in Northern Venice Lagoon, Italy. Sustainability 13, 4102. 10.3390/su13084102

Trifinopoulos, J., Nguyen, L.-T., von Haeseler, A., Minh, B.Q., 2016. W-IQ-TREE: a fast online phylogenetic tool for maximum likelihood analysis. Nucleic Acids Res 44, W232–W235. 10.1093/nar/gkw256

Wolf, M. A., Moro, I. 2013. Guida alle alghe dell’alto Adriatico. Dryades project. From https://dryades.units.it/home/index.php?procedure=ext_key_home&key_type=var&key_id=14624

Wolf, M.A., Sciuto, K., Maggs, C.A., De Barros-Barreto, M.B.B., Andreoli, C., Moro, I., 2011. *Ceramium* Roth (Ceramiales, Rhodophyta) from Venice lagoon (Adriatic Sea, Italy): Comparative studies of Mediterranean and Atlantic taxa. TAXON 60, 1584– 1595. 10.1002/tax.606004

Wong, J.X.W., Costantini, F., Merloni, N., Savelli, L., Geelen, D., Airoldi, L., 2018. The widespread and overlooked replacement of Spartina maritima by non-indigenous S. anglica and S. townsendii in north-western Adriatic saltmarshes. Biol Invasions 20, 1687–1702. 10.1007/s10530-017-1654-3

Wong, T.K.F., Ly-Trong, N., Ren, H., Demotte, P., Baños, H., Roger, A.J., Susko, E., Bielow, C., Maio, N.D., Goldman, N., Hahn, M.W., 2026. IQ-TREE 3: phylogenomic inference software using complex evolutionary models. Molecular Biology and Evolution, 43(5), p.msag117

Ye, J., Coulouris, G., Zaretskaya, I., Cutcutache, I., Rozen, S., & Madden, T. L. (2012). Primer-BLAST: A tool to design target-specific primers for polymerase chain reaction. BMC Bioinformatics, 13(1), 134. 10.1186/1471-2105-13-134

