## Supplementary figures and images for "Integrating morphological and molecular approaches to assess photosynthetic species diversity in the Venice Lagoon"

### Supplementary Figure 1

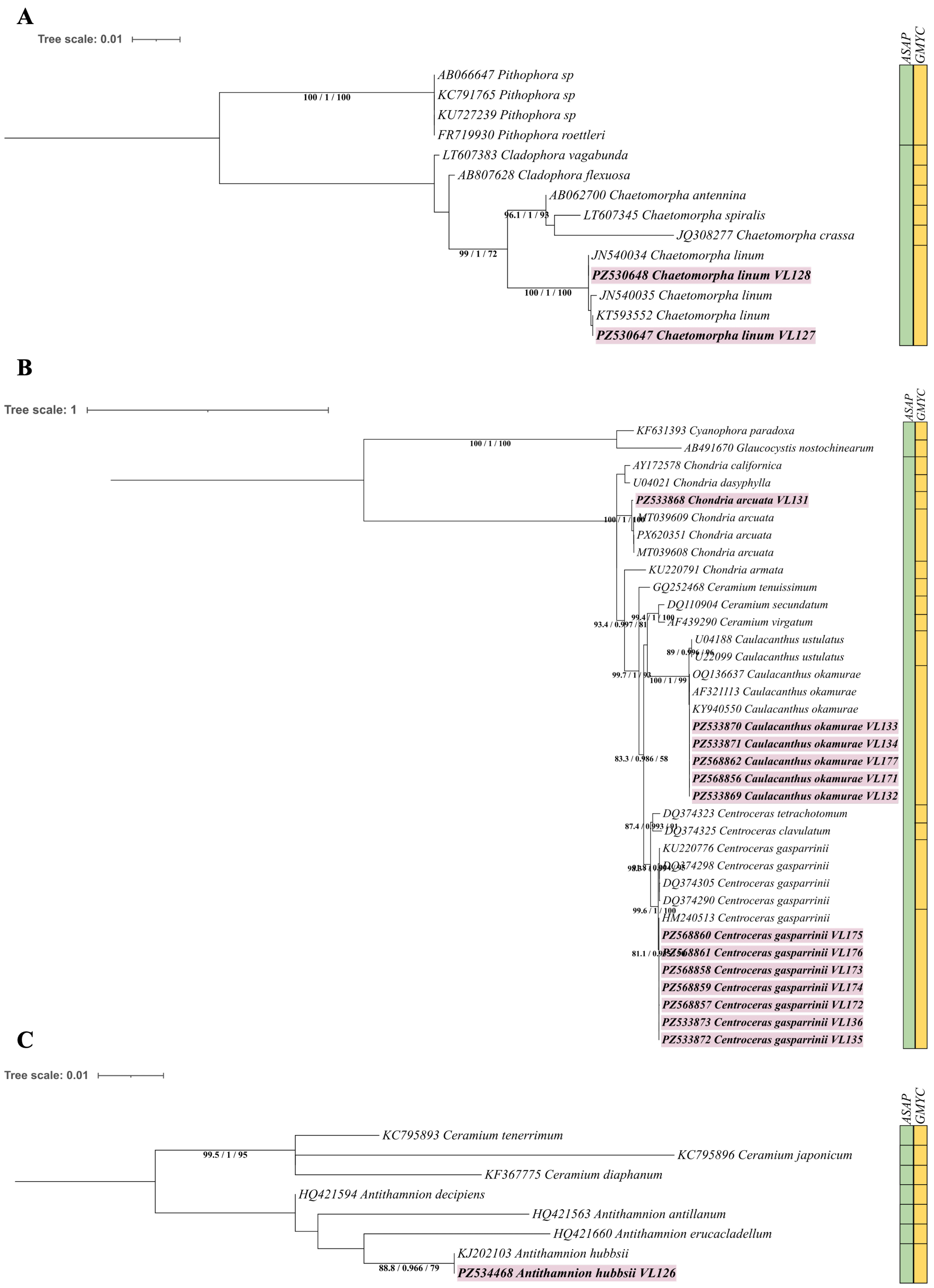

### Supplementary Figure 2

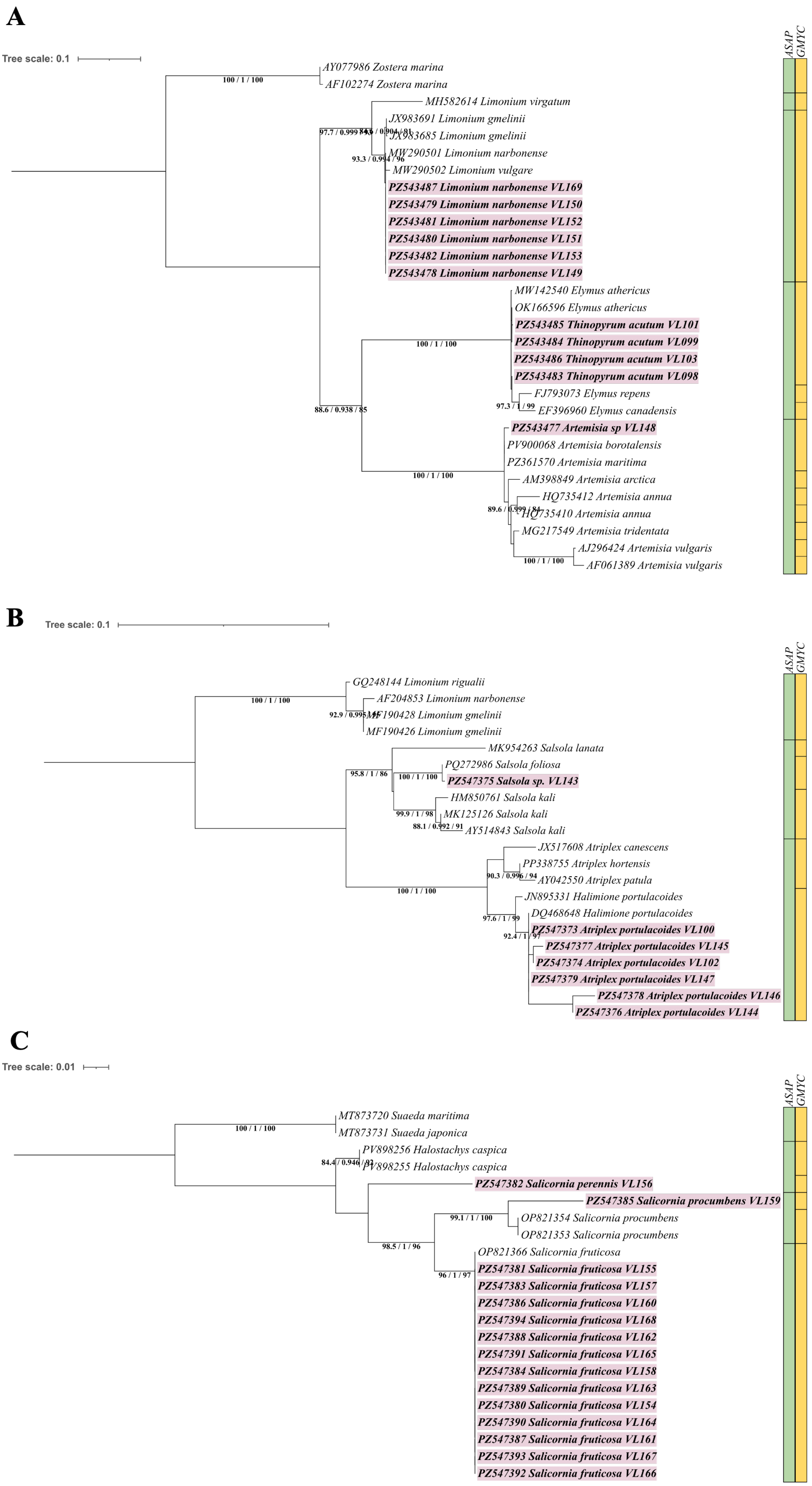
