## Supplementary Figures (1 to 12) and Tables (1 to 9) for "Integrating morphological and molecular approaches to assess photosynthetic species diversity in the Venice Lagoon": Supplementary material.docx

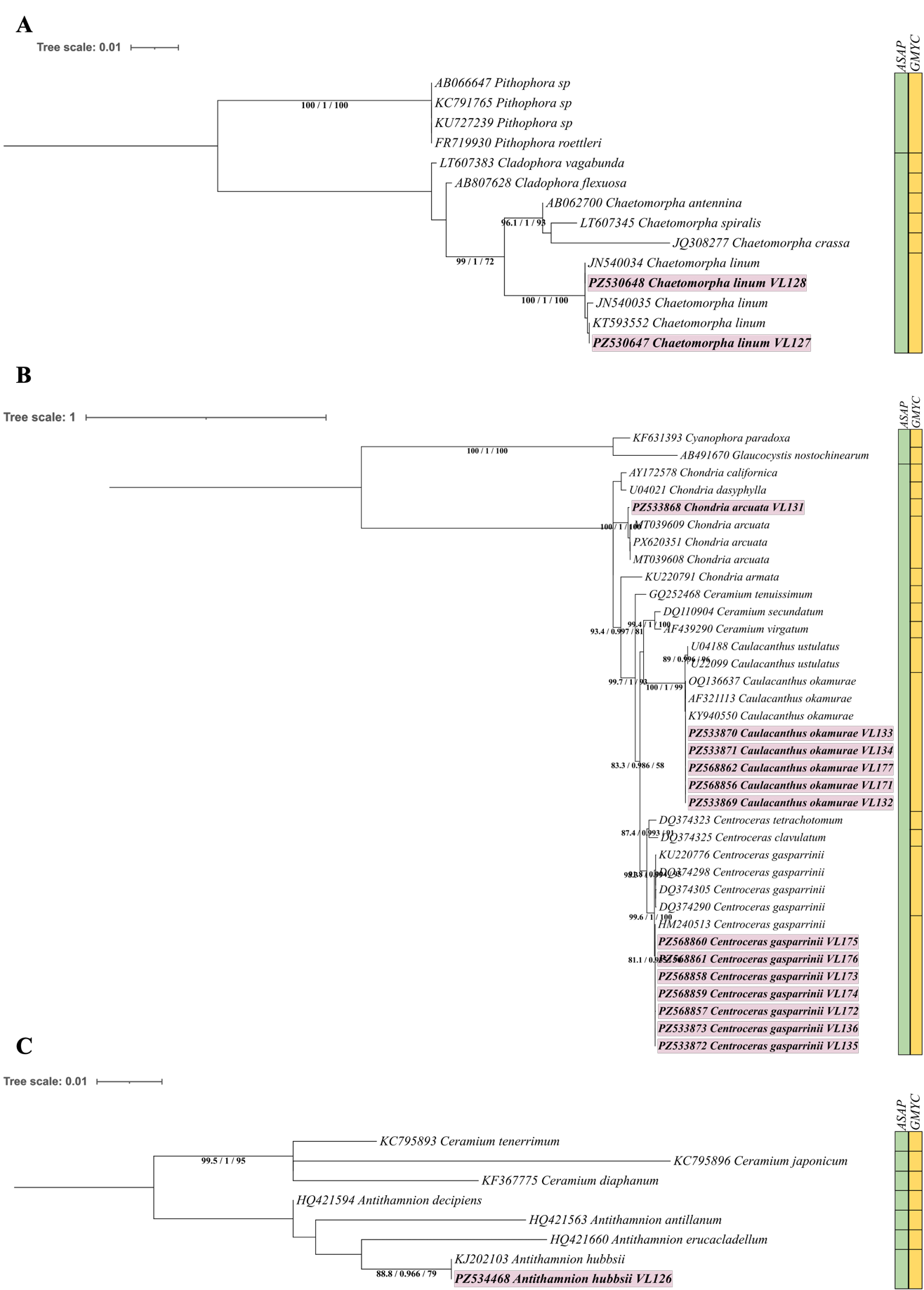

**Supplementary Figure 1. Phylogenetic reconstruction based on secondary markers of Chlorophyta and Rhodophyta from the Venice Lagoon**. Phylogenetic trees were inferred using IQ-TREE from 18S rRNA (**A**), rbcL (**B**), and UPA (**C**) sequences. Node values indicate SH-aLRT/aBayes/UFBoot support, with SH-aLRT values shown only when ≥80%. ASAP and GMYC species-delimitation results are represented by green and yellow outer circles, respectively. Survey samples are shown in bold and highlighted in light magenta, while GenBank sequences are labelled with accession numbers and species names (Supplementary Table 6).

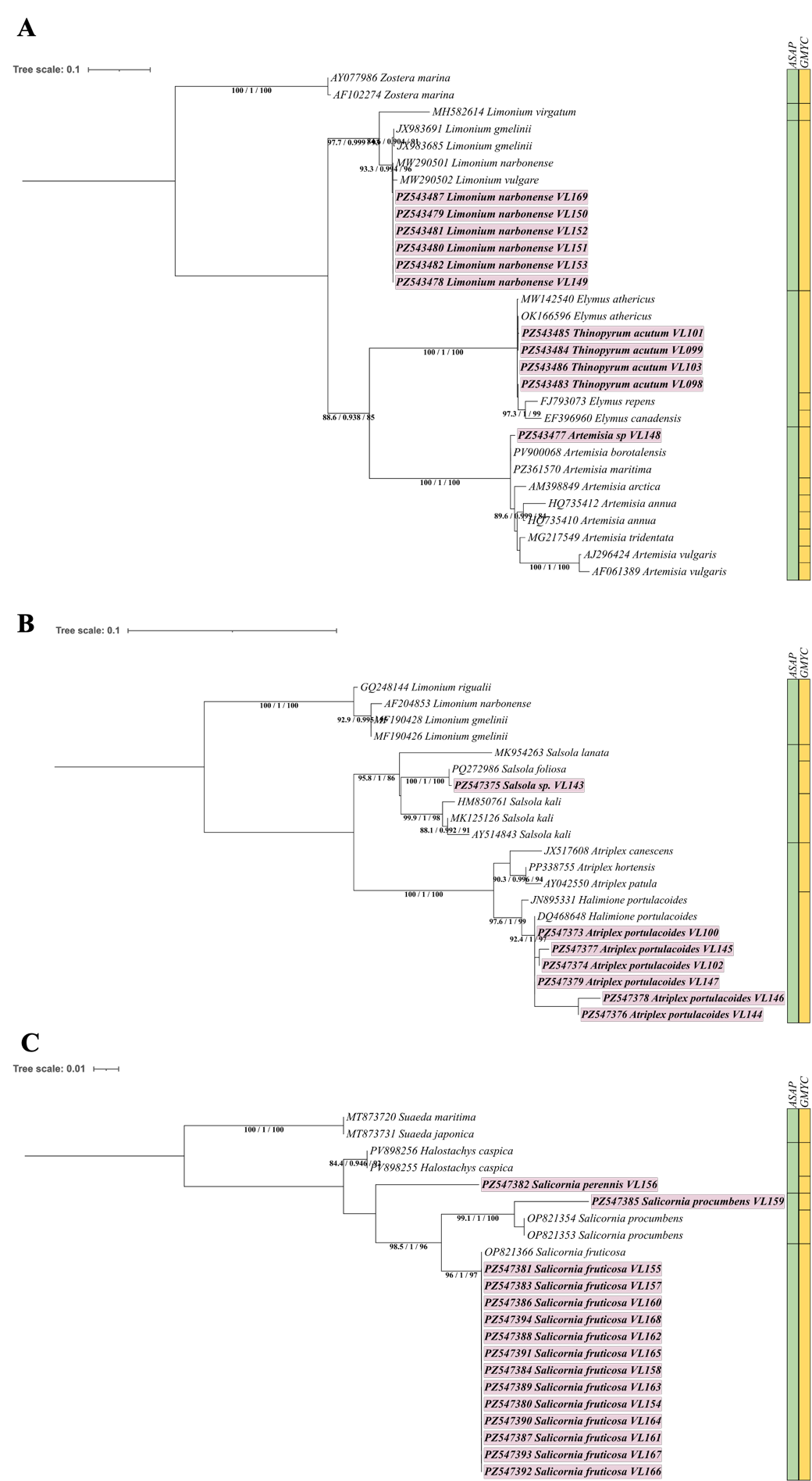

**Supplementary Figure 2. Phylogenetic reconstruction based on secondary markers of Tracheophyta from the Venice Lagoon**. Phylogenetic trees were inferred using IQ-TREE from ITS rRNA (**A**), matK (**B**), and psbA–trnH intergenic spacer (**C**) sequences. Node values indicate SH-aLRT/aBayes/UFBoot support, with SH-aLRT values shown only when ≥80%. ASAP and GMYC species-delimitation results are represented by green and yellow outer circles, respectively. Survey samples are shown in bold and highlighted in light magenta, while GenBank sequences are labelled with accession numbers and species names (Supplementary Table 6).

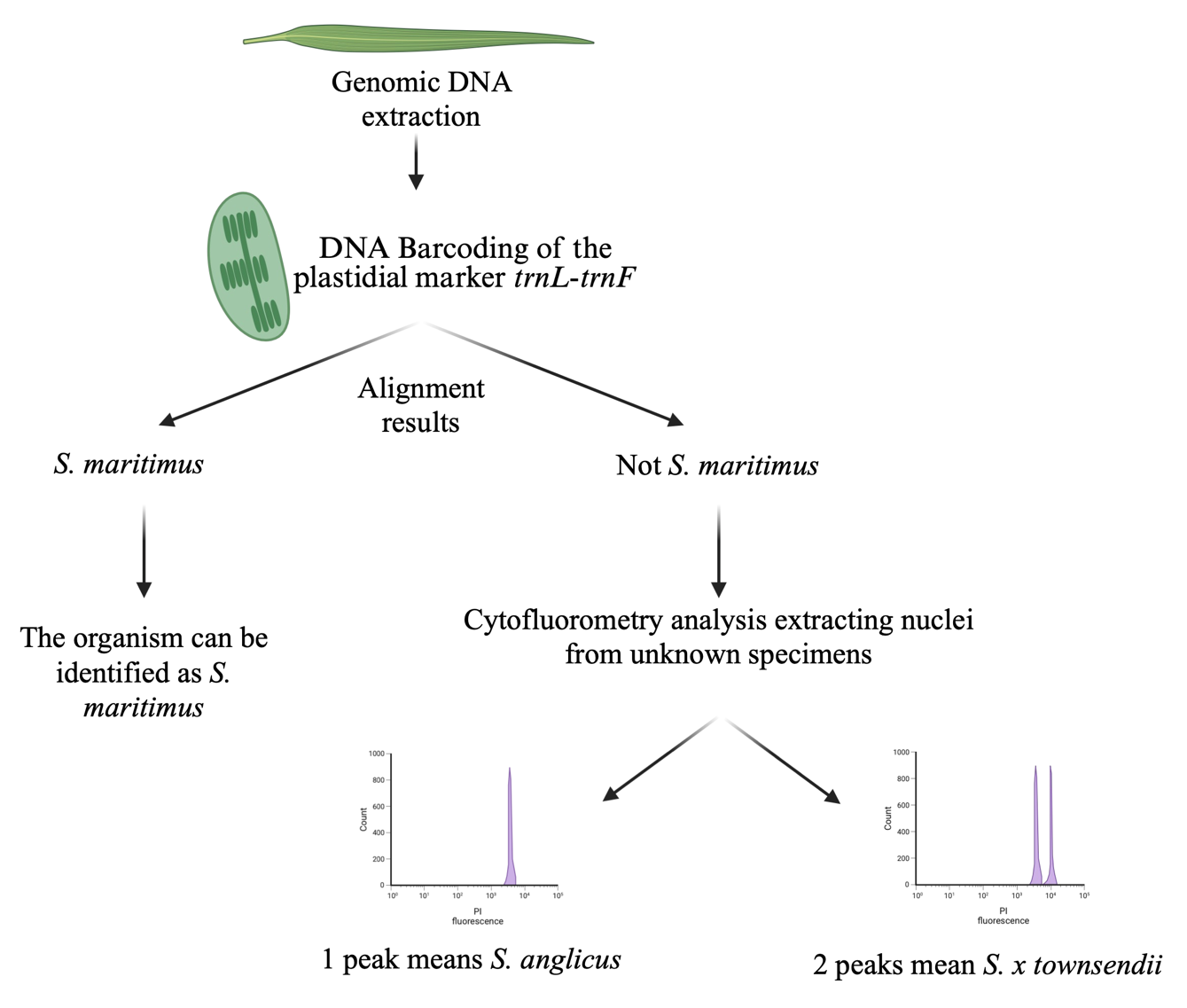

**Supplementary Figure 3. Conceptual workflow used to identify Sporobolus species in the Venice Lagoon.** The resulting sequences were compared with GenBank references: specimens matching S. maritimus were directly identified, whereas those matching the S. alterniflorus/S. anglicus lineage underwent genome-size analysis. Nuclei from each unresolved specimen were analysed together with those of a reference S. anglicus specimen by flow cytometry. A single peak indicated S. anglicus, whereas two distinct peaks identified S. x townsendii, whose genome is smaller than that of S. anglicus. Created with BioRender (<https://app.biorender.com/illustrations/67bc8292dc1daea8bb1c995c>).

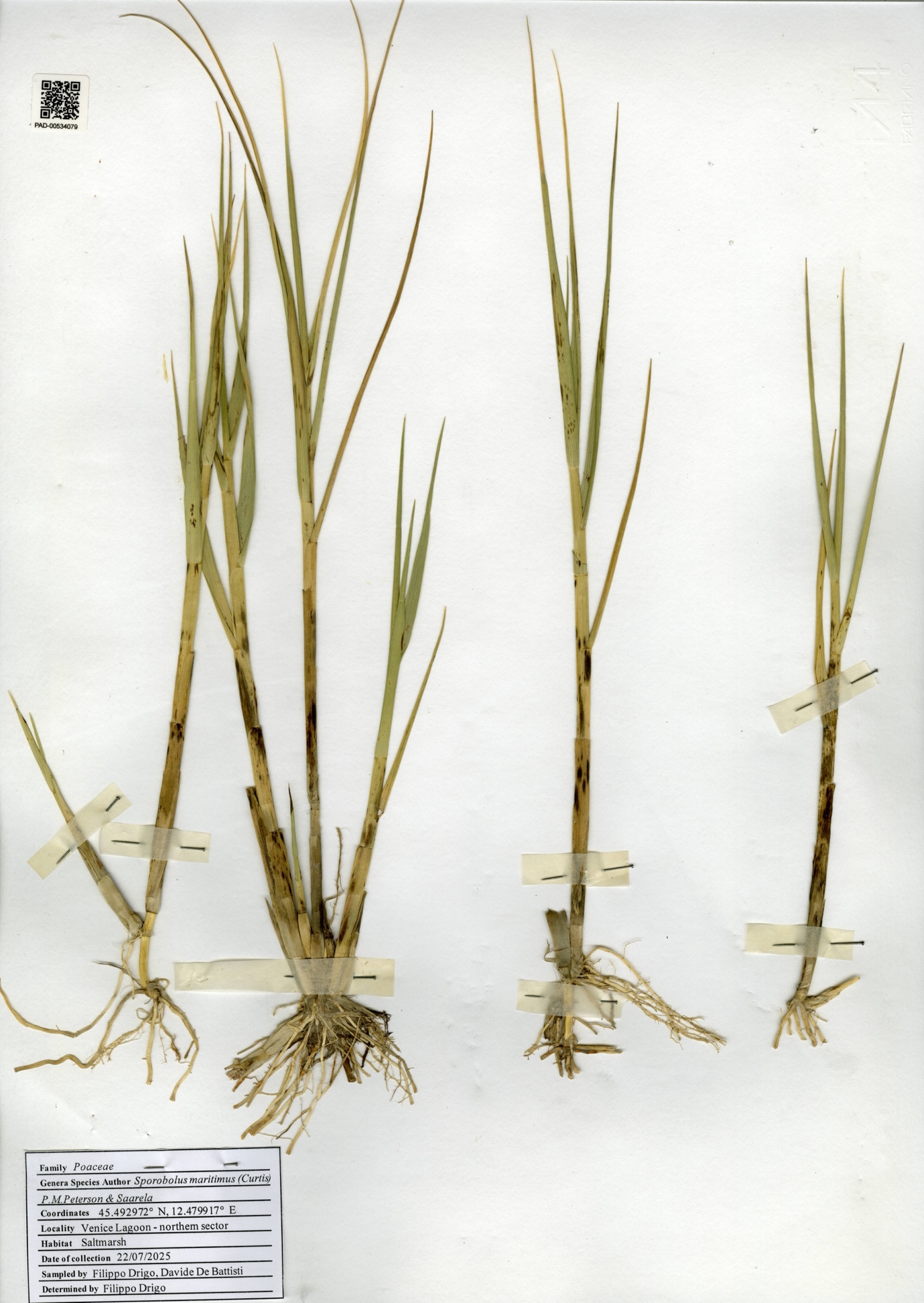

**Supplementary Figure 4. Herbarium specimen LN_17_9 of Sporobolus collected during the 2025 survey.** Herbarium sheet of specimen LN_17_9 (accession number PAD-00534079), identified as S. maritimus and deposited in the Herbarium at the Botanical Museum of the University of Padova.

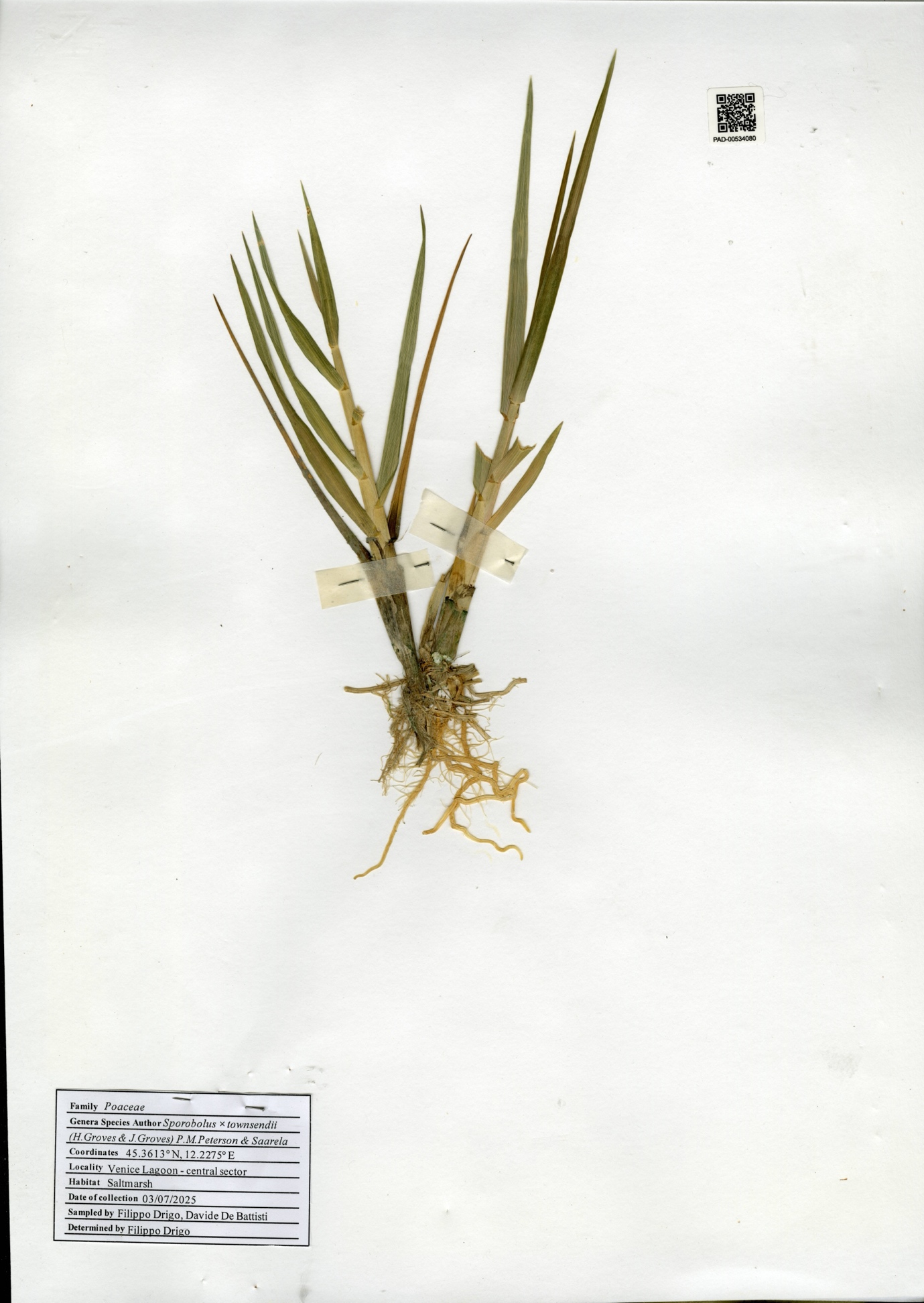

**Supplementary Figure 5. Herbarium specimen LC_11_7** **of Sporobolus collected during the 2025 survey.** Herbarium sheet of specimen LC_11_7 (accession number PAD-00534080), identified as S. x townsendii and deposited in the Herbarium at the Botanical Museum of the University of Padova.

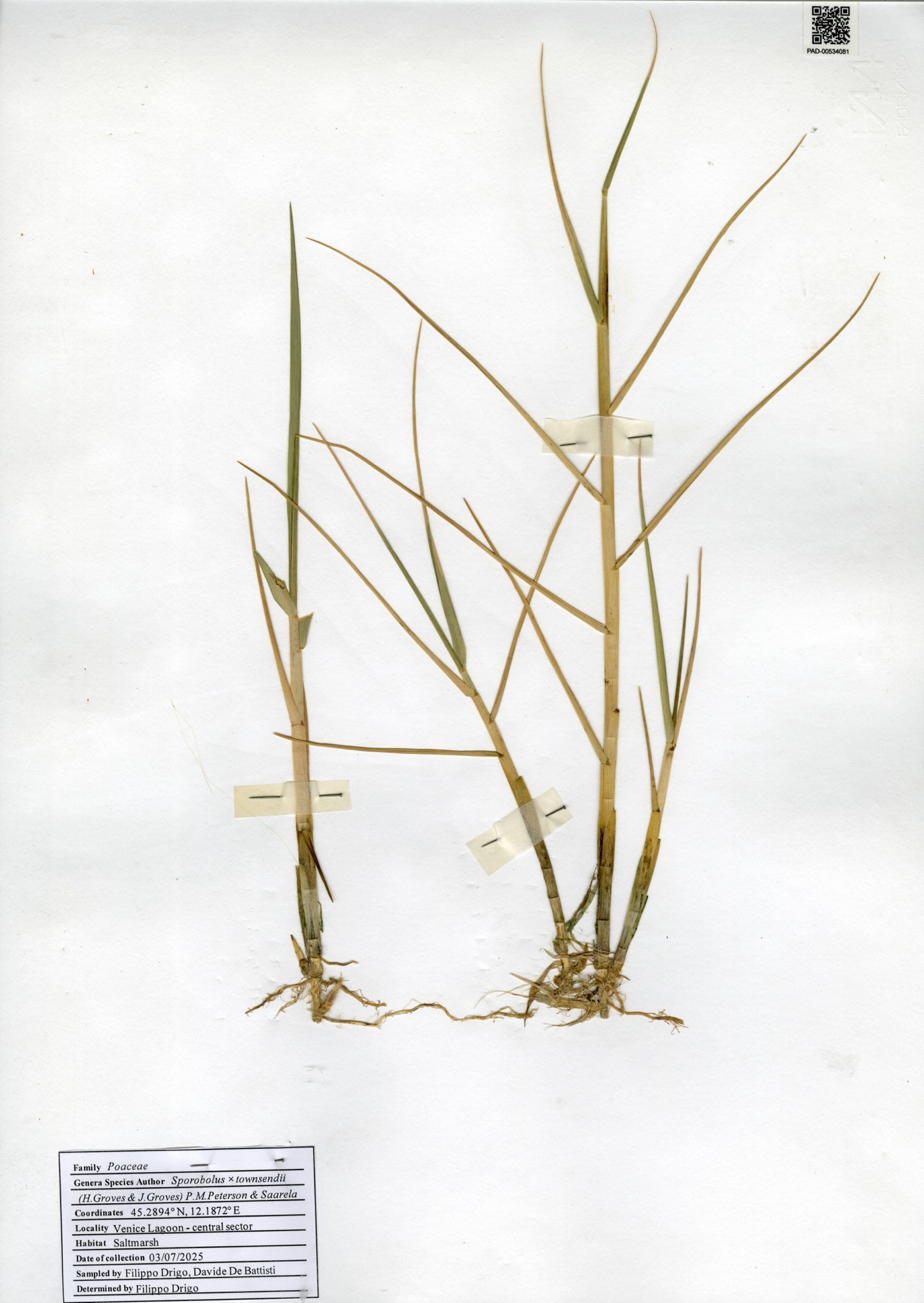

**Supplementary Figure 6. Herbarium specimen LC_7_2 of Sporobolus collected during the 2025 survey.** Herbarium sheet of specimen LC_7_2 (accession number PAD-00534081), identified as S. x townsendii and deposited in the Herbarium at the Botanical Museum of the University of Padova.

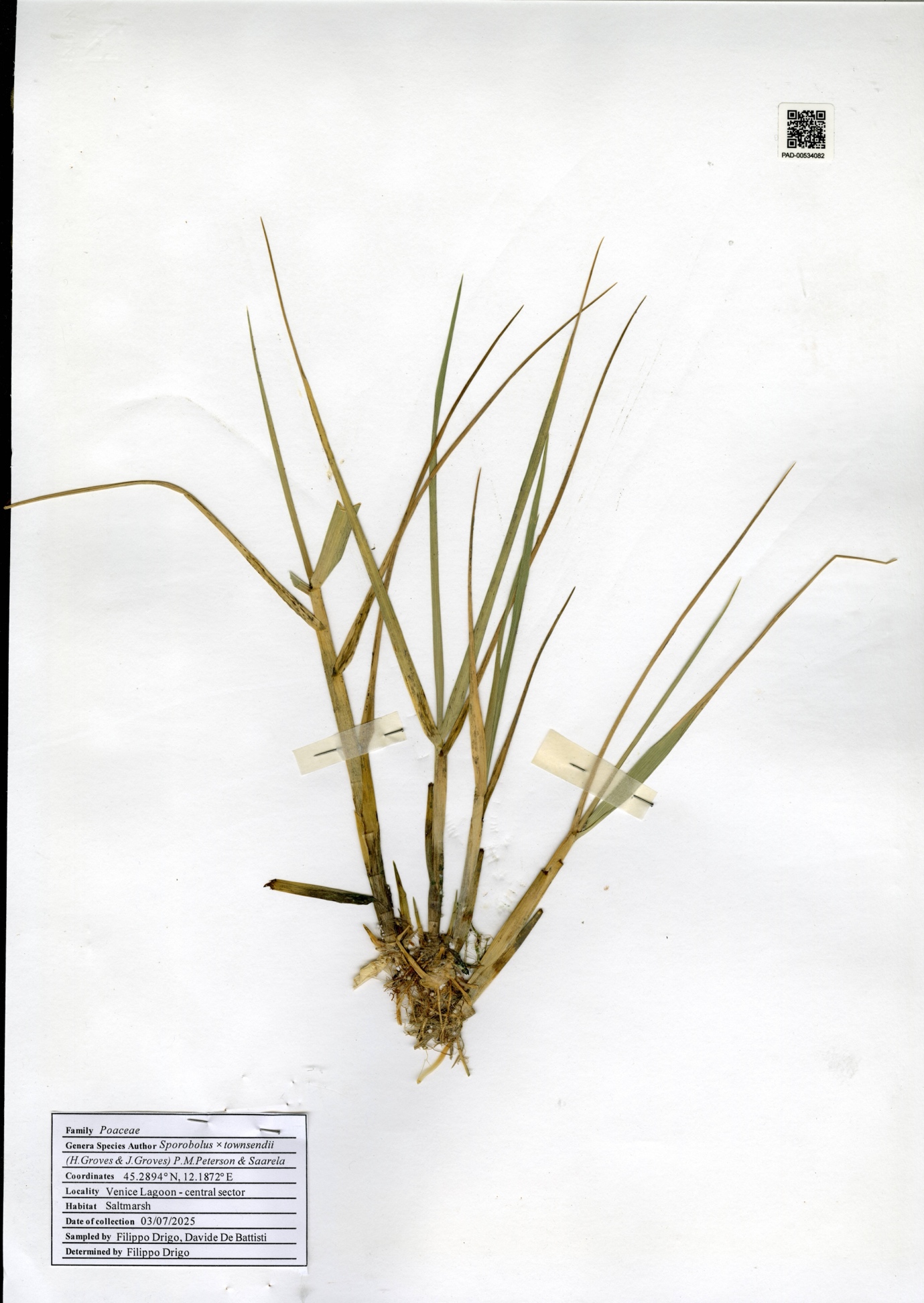

**Supplementary Figure 7. Herbarium specimen LC_7_3 of Sporobolus collected during the 2025 survey.** Herbarium sheet of specimen LC_7_3 (accession number PAD-00534082), identified as S. x townsendii and deposited in the Herbarium at the Botanical Museum of the University of Padova.

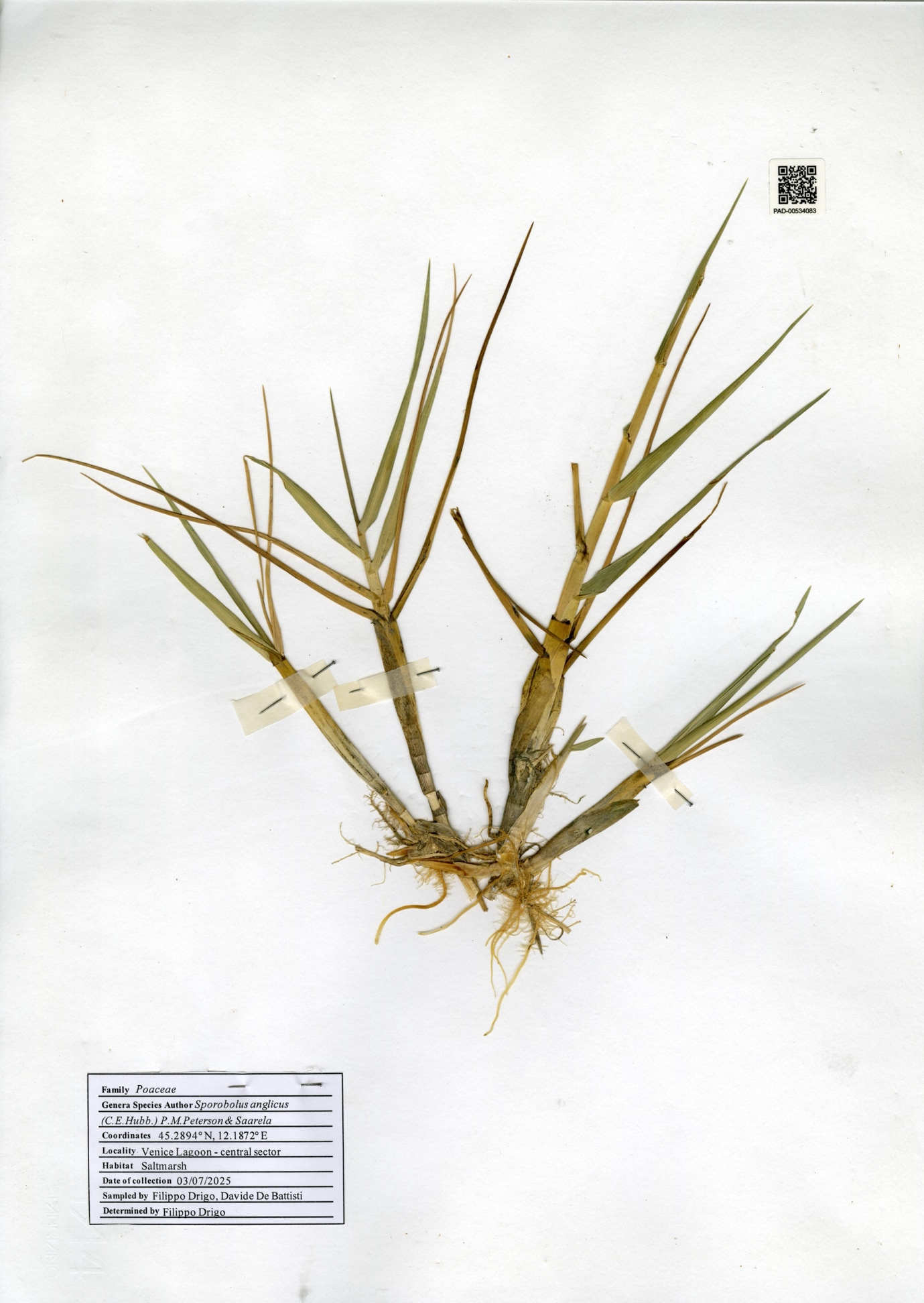

**Supplementary Figure 8. Herbarium specimen LC_7_1** **of Sporobolus collected during the 2025 survey.** Herbarium sheet of specimen LC_7_1 (accession number PAD-00534083), identified as S. anglicus and deposited in the Herbarium at the Botanical Museum of the University of Padova.

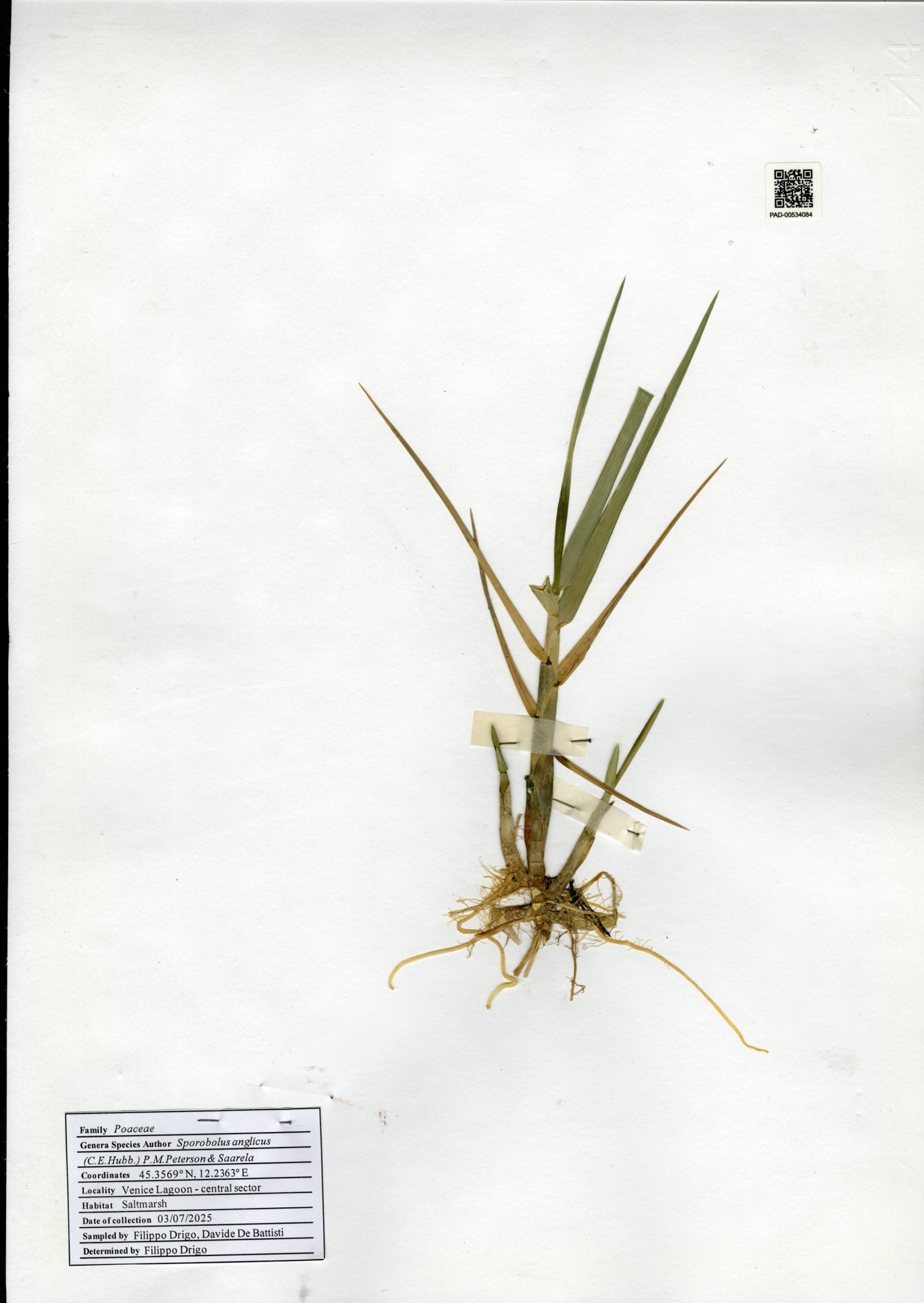

**Supplementary Figure 9. Herbarium specimen LC_12_2 of Sporobolus collected during the 2025 survey.** Herbarium sheet of specimen LC_12_2 (accession number PAD-00534084), identified as S. anglicus and deposited in the Herbarium at the Botanical Museum of the University of Padova.

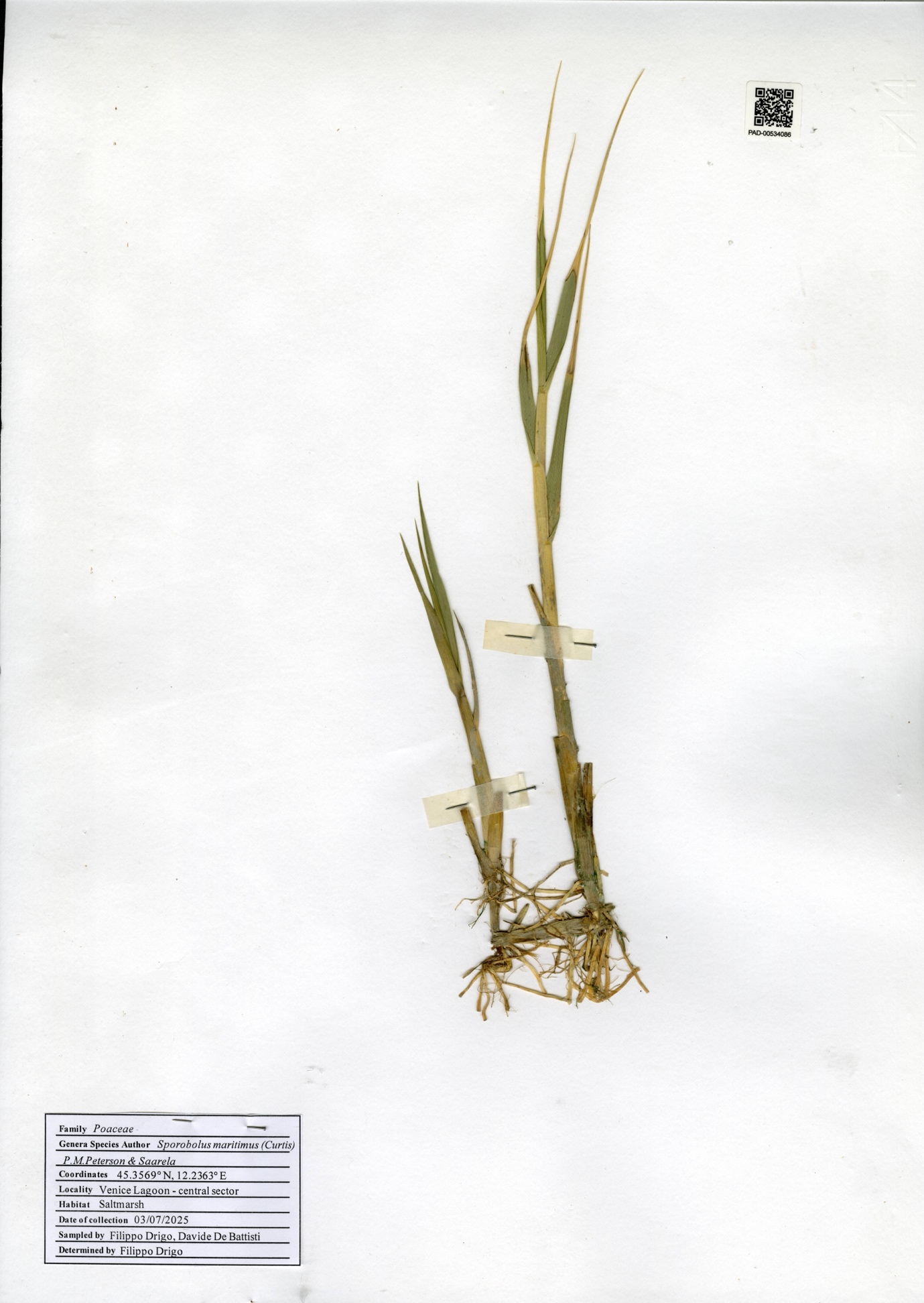

**Supplementary Figure 10. Herbarium specimen LC_12_4 of Sporobolus collected during the 2025 survey.** Herbarium sheet of specimen LC_12_4 specimen (accession number PAD-00534086), identified as S. maritimus and deposited in the Herbarium at the Botanical Museum of the University of Padova.

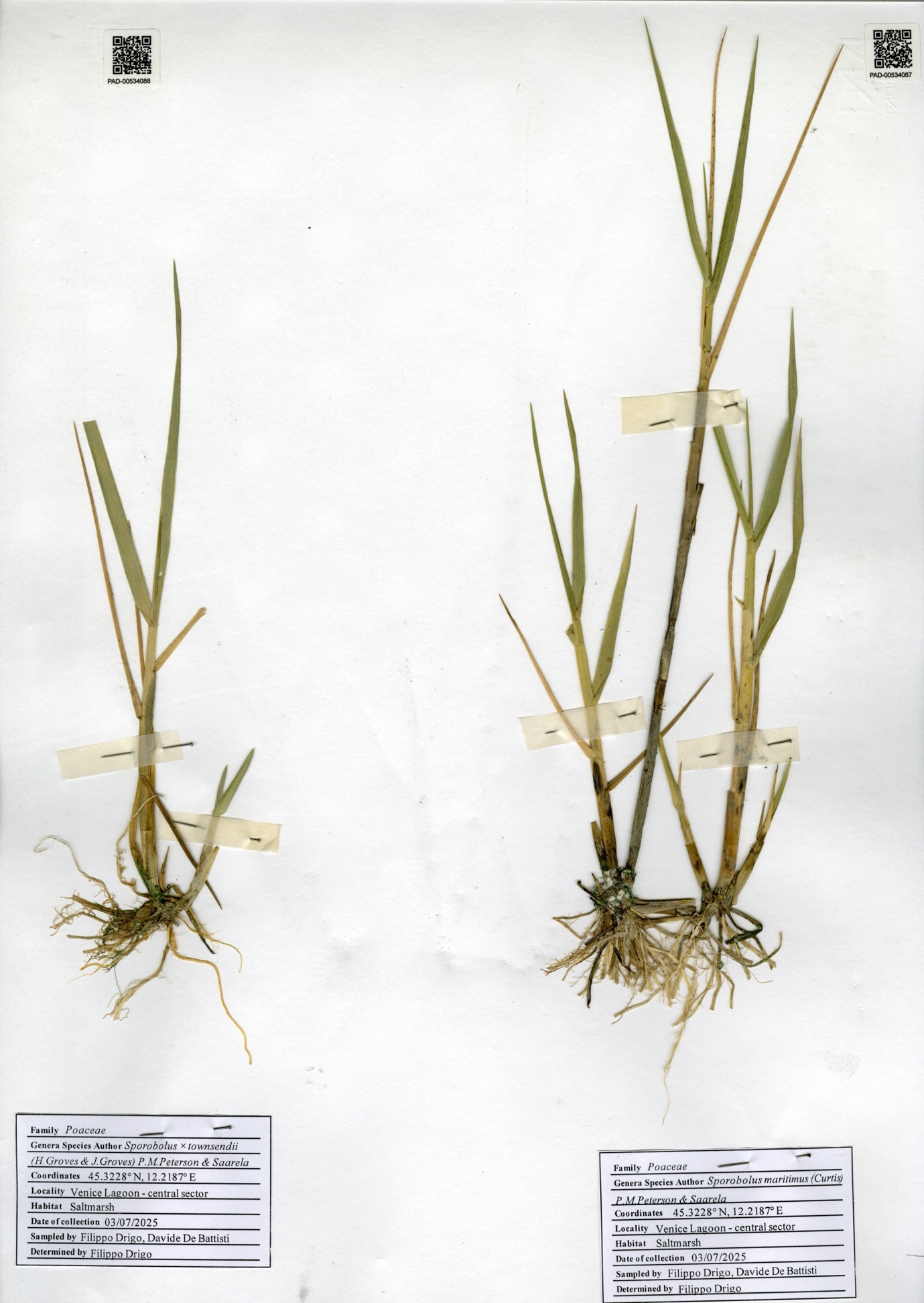

**Supplementary Figure 11. Herbarium specimen LC_8_6 and LC_8_7 of Sporobolus collected during the 2025 survey.** Herbarium sheet of specimen LC_8_6 (accession number PAD-00534088, left), identified as S. x townsendii and specimen LC_8_7 (accession number PAD-00534087, right), identified as S. maritimus and deposited in the Herbarium at the Botanical Museum of the University of Padova.

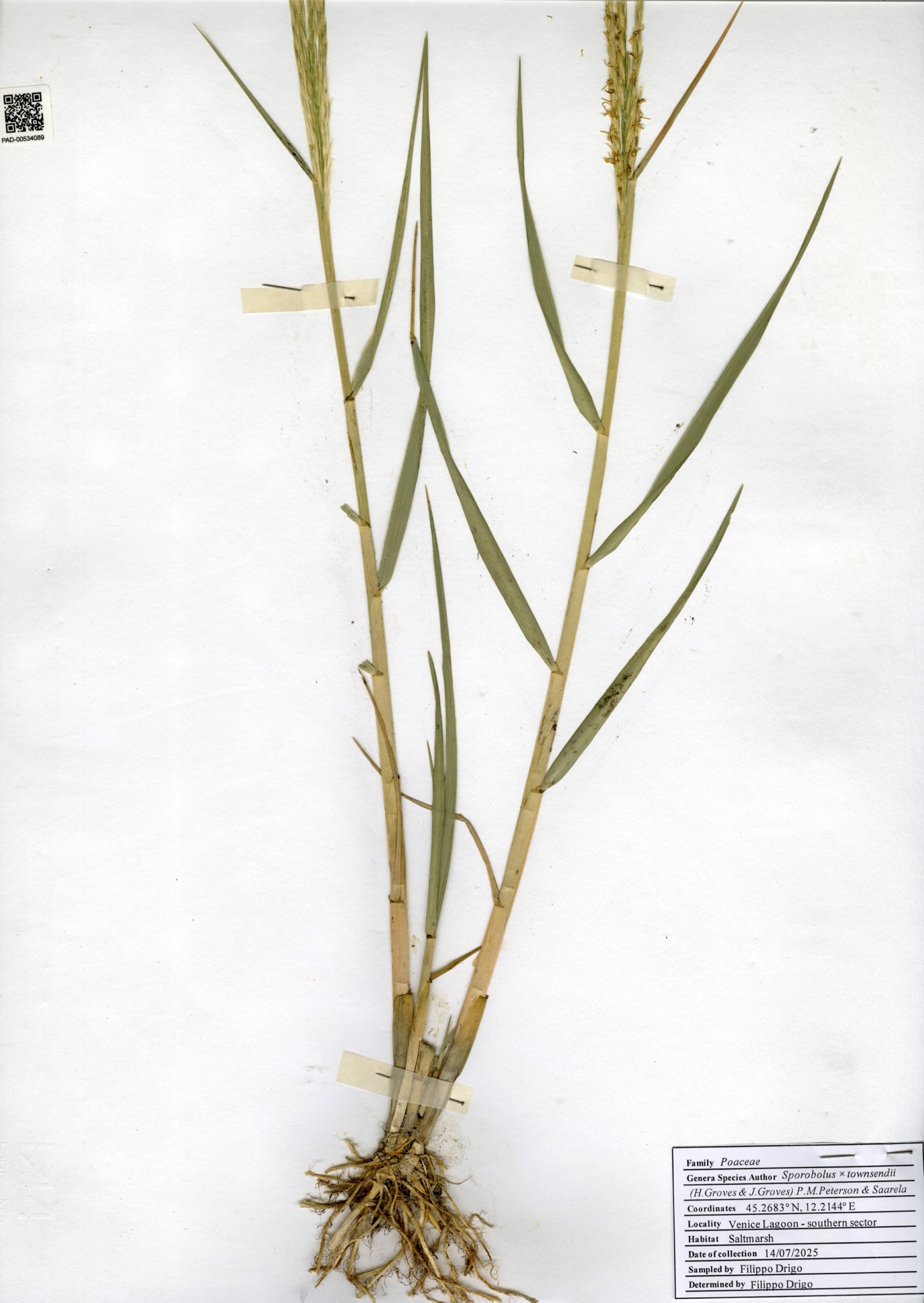

**Supplementary Figure 12. Herbarium specimen LS_5_4** **of Sporobolus collected during the 2025 survey.** Herbarium sheet of specimen LS_5_4 (accession number PAD-00534089), identified as S. x townsendii and deposited in the Herbarium at the Botanical Museum of the University of Padova.

**Supplementary Table 1. Sites sampled during the 2023 survey**. For each site, the table reports the site ID, geographical coordinates (latitude and longitude), habitat type, and sector of the Venice Lagoon.

| Site ID | Latitude (°N) | Longitude (°E) | Typology | Sector |
| --- | --- | --- | --- | --- |
| 2023_SM_1 | 45.2401 | 12.28805 | Salt marsh | South |
| 2023_WP_2 | 45.22582908 | 12.2823494 | Wood pole |  |
| 2023_WP_3 | 45.22489863 | 12.27273869 | Wood pole |  |
| 2023_WP_4 | 45.21215914 | 12.25840513 | Wood pole |  |
| 2023_WP_5 | 45.19825137 | 12.26621685 | Wood pole |  |
| 2023_WP_6 | 45.19004986 | 12.26959521 | Wood pole |  |
| 2023_SM_7 | 45.24379 | 12.22616 | Salt marsh |  |
| 2023_SM_8 | 45.36118 | 12.22743 | Salt marsh | Central |
| 2023_SM_9 | 45.35636 | 12.22974 | Salt marsh |  |
| 2023_SM_10 | 45.33247 | 12.22502 | Salt marsh |  |
| 2023_SM_11 | 45.34333 | 12.22419 | Salt marsh |  |
| 2023_SM_12 | 45.32449 | 12.21641 | Salt marsh |  |
| 2023_WP_13 | 45.2977833 | 12.1889667 | Wood pole |  |
| 2023_WP_14 | 45.2932 | 12.21805 | Wood pole |  |
| 2023_WP_15 | 45.2929167 | 12.2537667 | Wood pole |  |
| 2023_WP_16 | 45.2722 | 12.2549833 | Wood pole |  |
| 2023_WP_17 | 45.3477667 | 12.2431 | Wood pole |  |
| 2023_WP_18 | 45.3400833 | 12.2319167 | Wood pole |  |
| 2023_WP_19 | 45.3362333 | 12.2171167 | Wood pole |  |
| 2023_WP_20 | 45.33585 | 12.19975 | Wood pole |  |
| 2023_WP_21 | 45.3437333 | 12.2567333 | Wood pole |  |
| 2023_WP_22 | 45.2914167 | 12.3036833 | Wood pole |  |
| 2023_WP_23 | 45.3068667 | 12.3091667 | Wood pole |  |
| 2023_WP_24 | 45.3179833 | 12.3152 | Wood pole |  |
| 2023_WP_25 | 45.3411 | 12.28855 | Wood pole |  |
| 2023_WP_26 | 45.3307833 | 12.2829167 | Wood pole |  |
| 2023_WP_27 | 45.31945 | 12.2828833 | Wood pole |  |
| 2023_WP_28 | 45.3095 | 12.2689333 | Wood pole |  |
| 2023_WP_29 | 45.28605 | 12.2541167 | Wood pole |  |
| 2023_SM_30 | 45.27572 | 12.22481 | Salt marsh |  |
| 2023_SM_31 | 45.30602 | 12.22133 | Salt marsh |  |
| 2023_SM_32 | 45.32856587 | 12.22436254 | Salt marsh |  |

**Supplementary Table 2.** **Sites sampled during the 2024 survey.** For each site, the table reports the site ID, geographical coordinates (latitude and longitude), habitat type, and sector of the Venice Lagoon.

| Site ID | Latitude (°N) | Longitude (°E) | Typology | Sector |
| --- | --- | --- | --- | --- |
| VL_S_SM_1 | 45.19015 | 12.26905 | Salt marsh | South |
| VL_S_SM_2 | 45.19281 | 12.23346 |  |  |
| VL_S_SM_3 | 45.2261652 | 12.2308818 |  |  |
| VL_S_SM_4 | 45.20256 | 12.26267 |  |  |
| VL_S_WP_1 | 45.225817 | 12.281733 | Wood Poles |  |
| VL_S_WP_2 | 45.21705 | 12.270467 |  |  |
| VL_S_WP_3 | 45.192383 | 12.266417 |  |  |
| VL_S_WP_4 | 45.2123 | 12.258317 |  |  |
| VL_S_WP_5 | 45.219967 | 12.2524 |  |  |
| VL_C_SM_1 | 45.33036 | 12.22047 | Salt marsh | Central |
| VL_C_SM_2 | 45.342933 | 12.223733 |  |  |
| VL_C_SM_3 | 45.32 | 12.21975 |  |  |
| VL_C_SM_4 | 45.3566 | 12.22915 |  |  |
| VL_C_WP_1 | 45.32335 | 12.216933 | Wood Poles |  |
| VL_C_WP_2 | 45.328767 | 12.218183 |  |  |
| VL_C_WP_3 | 45.340083 | 12.231917 |  |  |
| VL_C_WP_4 | 45.351167 | 12.2621 |  |  |
| VL_C_WP_5 | 45.32785 | 12.316267 |  |  |
| VL_N_SM_1 | 45.493783 | 12.477423 | Salt marsh | North |
| VL_N_SM_2 | 45.482278 | 12.448805 |  |  |
| VL_N_SM_3 | 45.51935 | 12.475288 |  |  |
| VL_N_SM_4 | 45.48972 | 12.439348 |  |  |
| VL_N_WP_1 | 45.496436 | 12.482668 | Wood Poles |  |
| VL_N_WP_2 | 45.497865 | 12.483248 |  |  |
| VL_N_WP_3 | 45.521496 | 12.52988 |  |  |
| VL_N_WP_4 | 45.503388 | 12.526335 |  |  |
| VL_N_WP_5 | 45.504174 | 12.510027 |  |  |

**Supplementary Table 3. Sites sampled during the 2025 survey of Sporobolus species**. For each site, the table reports the site ID (LS, Southern Lagoon; LC, Central Lagoon; LN, Northern Lagoon), geographical coordinates (latitude and longitude), sampling date (DD-MM-YYYY), and sector of the Venice Lagoon.

| Site ID | Latitude (°N) | Longitude (°E) | Collection date | Sector |
| --- | --- | --- | --- | --- |
| LS 1 | 45.185389° | 12.253889° | 14-07-2025 | Southern |
| LS 2 | 45.2238° | 12.2244° | 14-07-2025 |  |
| LS 3 | 45.2258° | 12.2301° | 14-07-2025 |  |
| LS 4 | 45.2488° | 12.2219° | 24-07-2025 |  |
| LS 5 | 45.2683° | 12.2144° | 14-07-2025 |  |
| LS 6 | 45.2602° | 12.1955° | 24-07-2025 |  |
| LC 7 | 45.2894° | 12.1872° | 04-07-2025 | Central |
| LC 8 | 45.3228° | 12.2187° | 04-07-2025 |  |
| LC 9 | 45.32866° | 12.22448° | 04-07-2025 |  |
| LC10 | 45.342933° | 12.223733° | 04-07-2025 |  |
| LC 11 | 45.3613° | 12.2275° | 04-07-2025 |  |
| LC 12 | 45.3569° | 12.2363° | 04-07-2025 |  |
| LN 13 | 45.4494° | 12.4072° | 22-07-2025 | Northern |
| LN 14 | 45.4925° | 12.4077° | 22-07-2025 |  |
| LN 15 | 45.490806° | 12.439889° | 22-07-2025 |  |
| LN 16 | 45.483139° | 12.449194° | 22-07-2025 |  |
| LN 17 | 45.492972° | 12.479917° | 22-07-2025 |  |
| LN 18 | 45.520167° | 12.475611° | 22-07-2025 |  |

**Supplementary Table 4. List of primers and PCR amplification profiles used for molecular marker amplification.** Overview of the amplification profiles for each taxonomic group, the primer pair used, their sequence, and the amplification profile with the source article.

| Taxa | Gene marker | Primer Name | F/R | Primer sequence (5`-3`) | PCR conditions | | | Reference |
| --- | --- | --- | --- | --- | --- | --- | --- | --- |
|  |  |  |  |  | Denaturation | Cycles | Final extension |  |
| Chlorophyta | *tuf*A | tufGF4 | F | GGNGCNGCNCAAATGGAYGG | 94 °C for 4 min | 38 cycles of 94 °C for 1 min, 45 °C for 30 sec, 72 °C for 1 min | 72 °C for 7 min | Saunders & McDevit, 2012 |
|  |  | tufAR | R | CCTTCNCGAATMGCRAAWCGC |  |  |  |  |
| Heterokontophyta and Rhodophyta | *COI-5P* | GazF2 | F | CCAACCAYAAAGATATWGGTAC | 95 °C for 2 min | 5 cycles of 95 °C for 30 sec, 45 °C for 30 sec, 72 °C for 1 min, plus 35 cycles of 95 °C for 30 sec, 46.5 °C for 30 sec, 72 °C for 1 min | 72 °C for 7 min |  |
|  |  | GazR2 | R | GGATGACCAAARAACCAAAA |  |  |  |  |
|  |  | GWSFn | F | TCAACAAAYCAYAAAGATATYGG |  |  |  |  |
|  |  | GWSFa | F | CAAAYCAYAARGATATYGGAAC |  |  |  |  |
|  |  | GWSFt | F | CAAAYCAYAARGATATYGGTAC |  |  |  |  |
|  |  | GWSFi | F | CAAAYCAYAARGATATYGGIAC |  |  |  |  |
|  |  | GWSRx | R | ACTTCTGGRTGICCRAARAAYCA |  |  |  |  |
|  |  | GWSRi | R | GGRTGICCRAARAAYCARAA |  |  |  |  |
| Tracheophyta | *rbc*L | rbcLa-F | F | ATGTCACCACAAACAGAGACTAAAGC | 95 °C for 2 min | 45 cycles 95 °C for 30 sec, 50 °C for 1:30 min, 72 °C for 40 sec | 72 °C for 5 min | De Vere *et al.,* 2015 |
|  |  | rbcLr590 | R | AGTCCACCGCGTAGACATTCAT |  |  |  |  |
| *Scytosiphon* |  | PRB-F3 | F | TGTAAATGGATGCGTATGTG | 94 °C for 3 min | 35 cycles 94 °C for 45 sec, 55 °C for 1 min, 72 °C for 1 min | 72 °C for 5 min | Kogame *et al.,* 1999 |
|  |  | RSPR | R | AATAAAGGAAGACCCCATAATTCCCA |  |  |  |  |
| *Chondria* |  | F7 | F | AACTCTGTAGAACGNACAAG | 94 °C for 5 min | 35 cycles 94 °C for 1 min, 52 °C for 1 min, 72 °C for 2 min | 72 °C for 7 min | Gavio & Fredericq, 2002;  Kang & Kim, 2013 |
|  |  | JNR1 | R | CAATACGCATCCATTTACAA |  |  |  |  |
| *Vaucheria* |  | Spacer2F | F | GTACAAGAGCGTACTCGAAT | 94 °C for 5 min | 35 cycles 94 °C for 1 min, 52 °C for 1 min, 72 °C for 2:30 min | 72 °C for 5 min | Rybalka *et al.,* 2009 |
|  |  | Va1DPrbcL76 | R | AAACAGCCTTGTGTAACTCTC |  |  |  |  |
| *Centroceras* |  | F57 | F | GTAATTCCATATGCTAATGGG | 94 °C for 2 min | 30 cycles 94 °C for 40 sec, 50 °C for 40 sec, 72 °C for 40 sec | 72 °C for 5 min | Freshwater and Rueness*,* 1994; Wolf *et al.,* 2011 |
|  |  | R753 | R | GCTCTTTCATACATATCTTCC |  |  |  |  |
| *Limonium* | *ITS* rRNA | JK14 | F | GGAGAAGTCGTAACAAGGTTTCCG | 95 °C for 3 min | 35 cycles 95 °C for 30 sec, 50 °C for 1 min, 72 °C for 1 min | 72 °C for 5 min | De Castro *et al.,* 2013; Iamonico *et al.,* 2022 |
|  |  | SN3 | R | TTCGCTCGCCGTTACTAAGGG |  |  |  |  |
| *Thinopyrum* |  | ITS_F | F | TCGTAACAAGGTTTCCGTAGGTG | 94 °C for 3 min | 35 cycles 94 °C for 30 sec, 56 °C for 30 sec, 72 °C for 1:30 min | 72 °C for 5 min | Arterburn *et al.,* 2011 |
|  |  | ITS_R | R | TCCTCCGNCTTATTGATATGC |  |  |  |  |
| *Sporobolus* | *trnL-trnF* | trnLtrnF-F | F | ATAGATCCTGACATAGCAAACGAT | 95 °C for 3 min | 30 cycles of 95 °C for 30 sec, 48 °C for 30 sec, 72 °C for 1:15 min | 72 °C for 6 min | This study |
|  |  | trnLtrnF-R | R | TCATCCTAGTAGAGTATTTGCATCC |  |  |  |  |
| *Chaetomorpha* | *18S* rRNA | 18S-F | F | AATGGCTCGGTAAATCAGTT | 94 °C for 2 min | 30 cycles 94 °C for 30 sec, 55 °C for 1 min, 72 °C for 1 min | 72 °C for 10 min | Huang *et al.,* 2016 |
|  |  | 18S-R | R | AGTTGATGACTCGCGCTTAC |  |  |  |  |
| *Antithamnion* | *UPA* | p23SrV_f1 | F | GGACAGAAAGACCCTATGAA | 94 °C for 2 min | 35 cycles 94 °C for 20 sec, 55 °C for 30 sec, 72 °C for 30 sec | 72 °C for 10 min | Sherwood & Presting, 2007 |
|  |  | p23SrV_r1 | R | TCAGCCTGTTATCCCTAGAG |  |  |  |  |
| *Atriplex* | *mat*K | 1R_KIM-f | F | CGTACAGTACTTTTGTGTTTACGAG | 95 °C for 1 min | 35 cycles 95 °C for 30 sec, 50 °C for 1 min, 72 °C for 1 min | 72 °C for 10 min | Burgess *et al.,* 2011 |
|  |  | 3F_KIM-r | R | ACCCAGTCCATCTGGAAATCTTGGTTC |  |  |  |  |
| *Salicornia* | *psbA-trnH* | psbA-trnH_F | F | ATGTAAAGGGGCAGTGCTGG | 94 °C for 5 min | 35 cycles 94 °C for 45 sec, 53 °C for 30 sec, 72 °C for 1 min | 72 °C for 7 min | Bruni *et al.,* 2012; This study |
|  |  | psbA-trnH_R | R | ATCCACTTGGCTACATCCGC |  |  |  |  |

**Supplementary Table 5. Best-fit model determined by ModelFinder.** The table reports the best-fit model determined by ModelFinder for each phylogenetic reconstruction performed and the figure where the resulting phylogenetic tree is represented.

| **Taxa** | **Gene marker** | **Best-fit model (ModelFinder)** | **Figure** |
| --- | --- | --- | --- |
| Chlorophyta | *tuf*A | TIM3+F+I+G4 | Figure 2 |
|  | *18S* rRNA | TN+I | Supplementary Figure 1A |
| Xantophyceae | *COI-5P* | K3Pu+F+R2 | Figure 3A |
|  | *rbc*L | GTR+F+G4 | Figure 3B |
| Phaeophyceae | *COI-5P* | TN+F+G4 | Figure 4 |
| Rhodophyta | *COI-5P* | TIM+F+I+R4 | Figure 5 |
|  | *rbc*L | GTR+F+G4 | Supplementary Figure 1B |
|  | *UPA* | TPM2+I | Supplementary Figure 1C |
| Tracheophyta | *rbc*L | HKY+F+R3 | Figure 6 |
|  | *ITS* rRNA | TIM3e+I | Supplementary Figure 2A |
|  | *mat*K | TVM+F+G4 | Supplementary Figure 2B |
|  | *psbA-trnH* | K3Pu+F | Supplementary Figure 2C |

**Supplementary Table 6. GenBank accession numbers of reference sequences used for phylogenetic reconstruction.** For each sequence, the table reports the accession number, species name, taxonomic group, and molecular marker.

| **Accession** | **Species name** | **For which analysis used** | **Gene marker** |
| --- | --- | --- | --- |
| KT593552 | *Chaetomorpha linum* | Chlorophyta | *18S* rRNA |
| AB062700 | *Chaetomorpha antennina* | Chlorophyta | *18S* rRNA |
| LT607383 | *Cladophora vagabunda* | Chlorophyta | *18S* rRNA |
| AB807628 | *Cladophora flexuosa* | Chlorophyta | *18S* rRNA |
| JQ308277 | *Chaetomorpha crassa* | Chlorophyta | *18S* rRNA |
| LT607345 | *Chaetomorpha spiralis* | Chlorophyta | *18S* rRNA |
| FR719930 | *Pithophora roettleri* | Chlorophyta | *18S* rRNA |
| AB066647 | *Pithophora sp* | Chlorophyta | *18S* rRNA |
| KC791765 | *Pithophora sp* | Chlorophyta | *18S* rRNA |
| KU727239 | *Pithophora sp* | Chlorophyta | *18S* rRNA |
| JN540034 | *Chaetomorpha linum* | Chlorophyta | *18S* rRNA |
| JN540035 | *Chaetomorpha linum* | Chlorophyta | *18S* rRNA |
| KT290281 | *Blidingia minima* | Chlorophyta | *tuf*A |
| HQ610239 | *Blidingia chadefaudii* | Chlorophyta | *tuf*A |
| MH475464 | *Blidingia marginata* | Chlorophyta | *tuf*A |
| MN258036 | *Blidingia marginata* | Chlorophyta | *tuf*A |
| MH475463 | *Blidingia sp* | Chlorophyta | *tuf*A |
| HQ610252 | *Gayralia oxysperma* | Chlorophyta | *tuf*A |
| LK022443 | *Bryopsis muscosa* | Chlorophyta | *tuf*A |
| LK022441 | *Bryopsis corymbosa* | Chlorophyta | *tuf*A |
| LK022442 | *Bryopsis foliosa* | Chlorophyta | *tuf*A |
| LK022444 | *Bryopsis myosuroides* | Chlorophyta | *tuf*A |
| FJ432653 | *Bryopsis plumosa* | Chlorophyta | *tuf*A |
| AB008009 | *Bryopsis maxima* | Chlorophyta | *tuf*A |
| U09427 | *Codium fragile* | Chlorophyta | *tuf*A |
| FJ432662 | *Codium lucasii* | Chlorophyta | *tuf*A |
| KP685764 | *Codium bursa* | Chlorophyta | *tuf*A |
| KP685766 | *Codium bursa* | Chlorophyta | *tuf*A |
| MT160674 | *Ulva australis* | Chlorophyta | *tuf*A |
| MT076211 | *Ulva intestinalis* | Chlorophyta | *tuf*A |
| HQ610295 | *Ulva compressa* | Chlorophyta | *tuf*A |
| MW543061 | *Ulva lacinulata lectotype* | Chlorophyta | *tuf*A |
| MW543060 | *Ulva rigida holotype* | Chlorophyta | *tuf*A |
| OL421407 | *Ulva sp capillata holotype* | Chlorophyta | *tuf*A |
| OR723828 | *Ulva californica* | Chlorophyta | *tuf*A |
| OL421110 | *Ulva intestinalis* | Chlorophyta | *tuf*A |
| FJ535856 | *Codium platylobium* | Chlorophyta | *tuf*A |
| AB561084 | *Prasinococcus capsulatus* | Chlorophyta | *tuf*A |
| AB561083 | *Prasinophyceae sp* | Chlorophyta | *tuf*A |
| OL421504 | *Kornmannia leptoderma* | Chlorophyta | *tuf*A |
| OL421090 | *Kornmannia leptoderma* | Chlorophyta | *tuf*A |
| OL421134 | *Kornmannia leptoderma* | Chlorophyta | *tuf*A |
| LC877949 | *Gayralia oxysperma* | Chlorophyta | *tuf*A |
| MH538555 | *Kornmannia leptoderma* | Chlorophyta | *tuf*A |
| MF441478 | *Kornmannia leptoderma* | Chlorophyta | *tuf*A |
| MW242795 | *Gayralia brasiliensis* | Chlorophyta | *tuf*A |
| OR597303 | *Gayralia brasiliensis* | Chlorophyta | *tuf*A |
| KX845585 | *Codium fragile* | Chlorophyta | *tuf*A |
| LN877867 | *Codium fragile* | Chlorophyta | *tuf*A |
| MN853876 | *Codium fragile* | Chlorophyta | *tuf*A |
| LC768901 | *Bryopsis sp* | Chlorophyta | *tuf*A |
| NC_026795 | *Bryopsis plumosa* | Chlorophyta | *tuf*A |
| NC_013359 | *Bryopsis hypnoides* | Chlorophyta | *tuf*A |
| PQ652170 | *Bryopsis hypnoides* | Chlorophyta | *tuf*A |
| OR860994 | *Bryopsis pennata* | Chlorophyta | *tuf*A |
| OR860995 | *Bryopsis pennata* | Chlorophyta | *tuf*A |
| MT324546 | *Bryopsis pennata* | Chlorophyta | *tuf*A |
| JQ755423 | *Bryopsis plumosa* | Chlorophyta | *tuf*A |
| PQ733683 | *Ulva australis* | Chlorophyta | *tuf*A |
| PQ733682 | *Ulva australis* | Chlorophyta | *tuf*A |
| PQ733669 | *Ulva australis* | Chlorophyta | *tuf*A |
| PQ733666 | *Ulva australis* | Chlorophyta | *tuf*A |
| PQ733687 | *Ulva californica* | Chlorophyta | *tuf*A |
| PQ733702 | *Ulva capillata* | Chlorophyta | *tuf*A |
| PQ733696 | *Ulva californica* | Chlorophyta | *tuf*A |
| PQ733701 | *Ulva capillata* | Chlorophyta | *tuf*A |
| PQ733741 | *Ulva compressa* | Chlorophyta | *tuf*A |
| PQ733730 | *Ulva compressa* | Chlorophyta | *tuf*A |
| PQ733805 | *Ulva intestinalis* | Chlorophyta | *tuf*A |
| PQ733813 | *Ulva intestinalis* | Chlorophyta | *tuf*A |
| PQ733832 | *Ulva lacinulata* | Chlorophyta | *tuf*A |
| PQ733829 | *Ulva lacinulata* | Chlorophyta | *tuf*A |
| PQ733866 | *Ulva linza* | Chlorophyta | *tuf*A |
| PQ733869 | *Ulva linza* | Chlorophyta | *tuf*A |
| PQ733872 | *Ulva linza* | Chlorophyta | *tuf*A |
| MT160657 | *Ulva lacinulata* | Chlorophyta | *tuf*A |
| MW543061 | *Ulva lacinulata* | Chlorophyta | *tuf*A |
| OR734311 | *Ulva flexuosa subsp flexuosa* | Chlorophyta | *tuf*A |
| MT160670 | *Ulva rigida* | Chlorophyta | *tuf*A |
| MT078951 | *Ulva rigida* | Chlorophyta | *tuf*A |
| MT894983 | *Ulva rigida* | Chlorophyta | *tuf*A |
| MW543060 | *Ulva rigida* | Chlorophyta | *tuf*A |
| MK456404 | *Ulva fenestrata type* | Chlorophyta | *tuf*A |
| OR427287 | *Ulva compressa type* | Chlorophyta | *tuf*A |
| MT078955 | *Ulva lacinulata type* | Chlorophyta | *tuf*A |
| MH731007 | *Ulva expansa type* | Chlorophyta | *tuf*A |
| OL604521 | *Ulva australis* | Chlorophyta | *tuf*A |
| OL604520 | *Ulva australis* | Chlorophyta | *tuf*A |
| OL604523 | *Ulva australis* | Chlorophyta | *tuf*A |
| OL604522 | *Ulva australis* | Chlorophyta | *tuf*A |
| PDO0021 | *Codium lucasii* | Chlorophyta | *tuf*A |
| KZN2K4-24 | *Codium platylobium* | Chlorophyta | *tuf*A |
| BDA0059 | *Bryopsis pennata* | Chlorophyta | *tuf*A |
| GWS039110 | *Bryopsis hypnoides* | Chlorophyta | *tuf*A |
| GWS018364 | *Bryopsis plumosa* | Chlorophyta | *tuf*A |
| GWS004830 | *Gayralia oxysperma* | Chlorophyta | *tuf*A |
| NAT046 | *Kornmannia leptoderma* | Chlorophyta | *tuf*A |
| GWS043159 | *Blidingia chadefaudii* | Chlorophyta | *tuf*A |
| 0722_OST018 | *Blidingia minima* | Chlorophyta | *tuf*A |
| GWS021396 | *Blidingia sp* | Chlorophyta | *tuf*A |
| GWS004837 | *Blidingia marginata* | Chlorophyta | *tuf*A |
| 0722_GOS017 | *Ulva capillata* | Chlorophyta | *tuf*A |
| L3960382 | *Ulva flexuosa* | Chlorophyta | *tuf*A |
| LAK49 | *Ulva californica* | Chlorophyta | *tuf*A |
| GWS024562 | *Ulva lacinulata* | Chlorophyta | *tuf*A |
| GWS017830 | *Ulva intestinalis* | Chlorophyta | *tuf*A |
| L3960381 | *Ulva compressa* | Chlorophyta | *tuf*A |
| 02UA | *Ulva expansa* | Chlorophyta | *tuf*A |
| GWS014779 | *Ulva fenestrata* | Chlorophyta | *tuf*A |
| GWS023392 | *Ulva australis* | Chlorophyta | *tuf*A |
| MF768075 | *Gongolaria barbata* | Heterokontopphyta | *COI-5P* |
| MF768051 | *Ericaria amentacea* | Heterokontopphyta | *COI-5P* |
| MF768052 | *Ericaria amentacea* | Heterokontopphyta | *COI-5P* |
| MF768074 | *Gongolaria barbata* | Heterokontopphyta | *COI-5P* |
| EU681404 | *Dictyopteris polypodioides* | Heterokontopphyta | *COI-5P* |
| EU681424 | *Scytosiphon lomentaria* | Heterokontopphyta | *COI-5P* |
| KP101267 | *Sargassum siliquosum* | Heterokontopphyta | *COI-5P* |
| OP270186 | *Sargassum siliquosum* | Heterokontopphyta | *COI-5P* |
| KF281312 | *Scytosiphon lomentaria* | Heterokontopphyta | *COI-5P* |
| KF281380 | *Scytosiphon promiscuus holotype* | Heterokontopphyta | *COI-5P* |
| MN927527 | *Phaeocystis globosa* | Heterokontopphyta | *COI-5P* |
| MN927543 | *Phaeocystis globosa* | Heterokontopphyta | *COI-5P* |
| MT760761 | *Phaeocystis globosa* | Heterokontopphyta | *COI-5P* |
| KC782903 | *Sargassum muticum* | Heterokontopphyta | *COI-5P* |
| KY682972 | *Sargassum muticum* | Heterokontopphyta | *COI-5P* |
| MT760703 | *Phaeocystis globosa* | Heterokontopphyta | *COI-5P* |
| GWS017948 | *Scytosiphon lomentaria* | Heterokontopphyta | *COI-5P* |
| GWS011886 | *Scytosiphon promiscuus* | Heterokontopphyta | *COI-5P* |
| PTM1863 | *Sargassum muticum* | Heterokontopphyta | *COI-5P* |
| KX265516 | *Melanothamnus japonicus* | Rhodophyta | *COI-5P* |
| KM894012 | *Melanothamnus akkeshiensis* | Rhodophyta | *COI-5P* |
| KM894022 | *Melanothamnus akkeshiensis* | Rhodophyta | *COI-5P* |
| KJ960856 | *Melanothamnus harveyi* | Rhodophyta | *COI-5P* |
| KM893997 | *Melanothamnus harveyi* | Rhodophyta | *COI-5P* |
| KX265519 | *Melanothamnus sphaerocarpus* | Rhodophyta | *COI-5P* |
| KX265523 | *Melanothamnus sphaerocarpus* | Rhodophyta | *COI-5P* |
| KX265543 | *Melanothamnus yendoi* | Rhodophyta | *COI-5P* |
| KX265550 | *Melanothamnus yendoi* | Rhodophyta | *COI-5P* |
| KM894048 | *Melanothamnus japonicus* | Rhodophyta | *COI-5P* |
| KM894042 | *Melanothamnus japonicus* | Rhodophyta | *COI-5P* |
| KF714838 | *Agardhiella subulata* | Rhodophyta | *COI-5P* |
| KF714844 | *Agardhiella subulata* | Rhodophyta | *COI-5P* |
| KF714846 | *Agardhiella subulata* | Rhodophyta | *COI-5P* |
| KP210178 | *Gracilaria flabelliformis* | Rhodophyta | *COI-5P* |
| KF492773 | *Palisada perforata* | Rhodophyta | *COI-5P* |
| KJ202080 | *Solieria filiformis* | Rhodophyta | *COI-5P* |
| HM915934 | *Solieria robusta* | Rhodophyta | *COI-5P* |
| PV553641 | *Aglaothamnion scopula* | Rhodophyta | *COI-5P* |
| MH388695 | *Alsidium corallinum* | Rhodophyta | *COI-5P* |
| MH388698 | *Chondria baileyana* | Rhodophyta | *COI-5P* |
| MT683781 | *Chondria sp* | Rhodophyta | *COI-5P* |
| MT783832 | *Chondria sp* | Rhodophyta | *COI-5P* |
| HQ423044 | *Chondria arcuata* | Rhodophyta | *COI-5P* |
| OP800339 | *Chondria baileyana* | Rhodophyta | *COI-5P* |
| KU564345 | *Chondria baileyana* | Rhodophyta | *COI-5P* |
| HM573506 | *Polysiphonia echinata* | Rhodophyta | *COI-5P* |
| MF093996 | *Carradoriella elongata* | Rhodophyta | *COI-5P* |
| MF093998 | *Kapraunia schneideri* | Rhodophyta | *COI-5P* |
| MF093999 | *Polysiphonia scopulorum* | Rhodophyta | *COI-5P* |
| MF094000 | *Polysiphonia sertularioides* | Rhodophyta | *COI-5P* |
| MF094004 | *Polysiphonia stricta* | Rhodophyta | *COI-5P* |
| MN165073 | *Alsidium oliveiranum* | Rhodophyta | *COI-5P* |
| MN165074 | *Alsidium oliveiranum* | Rhodophyta | *COI-5P* |
| MF094017 | *Alsidium corallinum* | Rhodophyta | *COI-5P* |
| MN165075 | *Alsidium corallinum* | Rhodophyta | *COI-5P* |
| MF093968 | *Alsidium seaforthii* | Rhodophyta | *COI-5P* |
| MN165076 | *Alsidium seaforthii* | Rhodophyta | *COI-5P* |
| MN165082 | *Alsidium triquetrum* | Rhodophyta | *COI-5P* |
| MN165084 | *Alsidium triquetrum* | Rhodophyta | *COI-5P* |
| KX258833 | *Osmundea ramosissima* | Rhodophyta | *COI-5P* |
| KU566532 | *Osmundea oederi* | Rhodophyta | *COI-5P* |
| KM254320 | *Osmundea spectabilis* | Rhodophyta | *COI-5P* |
| KX258840 | *Palisada crustiformans* | Rhodophyta | *COI-5P* |
| KF492772 | *Palisada flagellifera* | Rhodophyta | *COI-5P* |
| GU223895 | *Palisada parvipapillata* | Rhodophyta | *COI-5P* |
| MF544099 | *Palisada tenerrima* | Rhodophyta | *COI-5P* |
| MN184283 | *Osmundea oederi* | Rhodophyta | *COI-5P* |
| OK209883 | *Palisada perforata* | Rhodophyta | *COI-5P* |
| JX096542 | *Gelidium crinale* | Rhodophyta | *COI-5P* |
| KX423473 | *Gelidium galapagense* | Rhodophyta | *COI-5P* |
| OQ656734 | *Gelidium crinale* | Rhodophyta | *COI-5P* |
| OQ656730 | *Gelidium crinale* | Rhodophyta | *COI-5P* |
| KU958465 | *Gelidium crinale* | Rhodophyta | *COI-5P* |
| OQ656729 | *Gelidium crinale* | Rhodophyta | *COI-5P* |
| KT208040 | *Gelidium calidum* | Rhodophyta | *COI-5P* |
| MW760396 | *Gelidium americanum* | Rhodophyta | *COI-5P* |
| JQ843331 | *Gracilaria gracilis* | Rhodophyta | *COI-5P* |
| MW465693 | *Gracilaria gracilis* | Rhodophyta | *COI-5P* |
| KF714847 | *Gracilaria gracilis* | Rhodophyta | *COI-5P* |
| FJ499513 | *Gracilaria pacifica* | Rhodophyta | *COI-5P* |
| KM254246 | *Gracilaria pacifica* | Rhodophyta | *COI-5P* |
| MG199570 | *Gracilaria arcuata* | Rhodophyta | *COI-5P* |
| NC_058681 | *Gracilaria multipartita* | Rhodophyta | *COI-5P* |
| NC_058682 | *Gracilaria ornata* | Rhodophyta | *COI-5P* |
| OL690498 | *Gracilaria parva* | Rhodophyta | *COI-5P* |
| OL690495 | *Gracilaria parva* | Rhodophyta | *COI-5P* |
| HQ322059 | *Gracilaria vermiculophylla* | Rhodophyta | *COI-5P* |
| HQ322046 | *Gracilaria vermiculophylla* | Rhodophyta | *COI-5P* |
| HQ322032 | *Gracilaria parvispora* | Rhodophyta | *COI-5P* |
| FJ499509 | *Gracilaria gracilis* | Rhodophyta | *COI-5P* |
| KM214578 | *Hypnea spinella* | Rhodophyta | *COI-5P* |
| KU905111 | *Hypnea spinella* | Rhodophyta | *COI-5P* |
| KU905089 | *Hypnea cervicornis* | Rhodophyta | *COI-5P* |
| HQ422674 | *Hypnea cervicornis* | Rhodophyta | *COI-5P* |
| EF136606 | *Hypnea flexicaulis* | Rhodophyta | *COI-5P* |
| KP725277 | *Hypnea musciformis* | Rhodophyta | *COI-5P* |
| KP725276 | *Hypnea musciformis* | Rhodophyta | *COI-5P* |
| GQ141881 | *Hypnea musciformis* | Rhodophyta | *COI-5P* |
| HQ422612 | *Hypnea musciformis* | Rhodophyta | *COI-5P* |
| KU566543 | *Osmundea truncata* | Rhodophyta | *COI-5P* |
| KU566542 | *Osmundea truncata* | Rhodophyta | *COI-5P* |
| KU566544 | *Osmundea truncata* | Rhodophyta | *COI-5P* |
| MF120730 | *Carradoriella elongata* | Rhodophyta | *COI-5P* |
| MF409425 | *Carradoriella elongata* | Rhodophyta | *COI-5P* |
| MF409433 | *Carradoriella elongata* | Rhodophyta | *COI-5P* |
| MF120623 | *Carradoriella virgata* | Rhodophyta | *COI-5P* |
| MF120783 | *Carradoriella virgata* | Rhodophyta | *COI-5P* |
| MF543366 | *Kapraunia schneideri* | Rhodophyta | *COI-5P* |
| MF120708 | *Kapraunia schneideri* | Rhodophyta | *COI-5P* |
| MF120766 | *Polysiphonia morrowii* | Rhodophyta | *COI-5P* |
| MF120685 | *Polysiphonia morrowii* | Rhodophyta | *COI-5P* |
| HM573514 | *Kapraunia schneideri* | Rhodophyta | *COI-5P* |
| HM573516 | *Kapraunia schneideri* | Rhodophyta | *COI-5P* |
| HM573515 | *Kapraunia schneideri* | Rhodophyta | *COI-5P* |
| MG030803 | *Spermothamnion cymosum* | Rhodophyta | *COI-5P* |
| MG030804 | *Spermothamnion cymosum* | Rhodophyta | *COI-5P* |
| MF597664 | *Spermothamnion cymosum* | Rhodophyta | *COI-5P* |
| AB491672 | *Cyanophora paradoxa* | Rhodophyta | *COI-5P* |
| KF631389 | *Cyanophora paradoxa* | Rhodophyta | *COI-5P* |
| KF631391 | *Cyanophora tetracyanea* | Rhodophyta | *COI-5P* |
| KF631390 | *Cyanophora biloba* | Rhodophyta | *COI-5P* |
| AB491671 | *Glaucocystis nostochinearum* | Rhodophyta | *COI-5P* |
| PQ826726 | *Caulacanthus okamurae* | Rhodophyta | *COI-5P* |
| KJ960508 | *Caulacanthus ustulatus* | Rhodophyta | *COI-5P* |
| OQ174559 | *Caulacanthus ustulatus* | Rhodophyta | *COI-5P* |
| OQ174562 | *Caulacanthus ustulatus* | Rhodophyta | *COI-5P* |
| NC_047435 | *Caulacanthus okamurae* | Rhodophyta | *COI-5P* |
| MT193839 | *Caulacanthus okamurae* | Rhodophyta | *COI-5P* |
| MT996228 | *Palisada sp* | Rhodophyta | *COI-5P* |
| KX258838 | *Palisada sp* | Rhodophyta | *COI-5P* |
| OP297574 | *Palisada perforata* | Rhodophyta | *COI-5P* |
| MH388710 | *Palisada perforata* | Rhodophyta | *COI-5P* |
| LC820980 | *Palisada intermedia* | Rhodophyta | *COI-5P* |
| KX258835 | *Palisada sp* | Rhodophyta | *COI-5P* |
| MG030782 | *Palisada tenerrima* | Rhodophyta | *COI-5P* |
| PP974343 | *Palisada corallopsis* | Rhodophyta | *COI-5P* |
| PP974342 | *Palisada corallopsis* | Rhodophyta | *COI-5P* |
| EU194956 | *Aglaothamnion halliae* | Rhodophyta | *COI-5P* |
| EU194959 | *Aglaothamnion sepositum* | Rhodophyta | *COI-5P* |
| EU194958 | *Aglaothamnion roseum* | Rhodophyta | *COI-5P* |
| EU194960 | *Callithamnion tetragonum* | Rhodophyta | *COI-5P* |
| PP735967 | *Callithamnion acutum* | Rhodophyta | *COI-5P* |
| LC820911 | *Antithamnion nipponicum* | Rhodophyta | *COI-5P* |
| HQ4227951 | *Antithamnion antillanum* | Rhodophyta | *COI-5P* |
| HQ423091 | *Antithamnion antillanum* | Rhodophyta | *COI-5P* |
| HQ423063 | *Antithamnion decipiens* | Rhodophyta | *COI-5P* |
| HQ423013 | *Antithamnion sp* | Rhodophyta | *COI-5P* |
| HQ423104 | *Antithamnion sp* | Rhodophyta | *COI-5P* |
| HQ412555 | *Antithamnion hubbsii* | Rhodophyta | *COI-5P* |
| KJ202083 | *Antithamnion hubbsii* | Rhodophyta | *COI-5P* |
| KM254403 | *Antithamnion sp* | Rhodophyta | *COI-5P* |
| GWS020296 | *Callithamnion acutum* | Rhodophyta | *COI-5P* |
| GWS005753 | *Aglaothamnion halliae* | Rhodophyta | *COI-5P* |
| GWS015092 | *Spermothamnion cymosum* | Rhodophyta | *COI-5P* |
| MT676065 | *Polysiphonia scopulorum* | Rhodophyta | *COI-5P* |
| KM609436 | *Polysiphonia echinata* | Rhodophyta | *COI-5P* |
| GWS032169 | *Chondria baileyana* | Rhodophyta | *COI-5P* |
| GWS021223 | *Osmundea spectabilis* | Rhodophyta | *COI-5P* |
| SP365646 | *Hypnea cervicornis* | Rhodophyta | *COI-5P* |
| KP708623 | *Hypnea flexicaulis* | Rhodophyta | *COI-5P* |
| GWS001078 | *Solieria filiformis* | Rhodophyta | *COI-5P* |
| GWS001590 | *Solieria robusta* | Rhodophyta | *COI-5P* |
| GWS017856 | *Agardhiella subulata* | Rhodophyta | *COI-5P* |
| MN165079 | *Alsidium seaforthii* | Rhodophyta | *COI-5P* |
| GWS018610 | *Gracilaria vermiculophylla* | Rhodophyta | *COI-5P* |
| GWS022199 | *Gracilaria pacifica* | Rhodophyta | *COI-5P* |
| GWS014616 | *Gracilaria gracilis* | Rhodophyta | *COI-5P* |
| GWS017314 | *Gracilaria arcuata* | Rhodophyta | *COI-5P* |
| ON527807 | *Gracilaria parvispora* | Rhodophyta | *COI-5P* |
| PHYKOS-4519 | *Gracilaria parva* | Rhodophyta | *COI-5P* |
| TFP08_0072 | *Gracilaria flabelliformis* | Rhodophyta | *COI-5P* |
| MZ336090 | *Gracilaria multipartita* | Rhodophyta | *COI-5P* |
| GWS012474 | *Melanothamnus harveyi* | Rhodophyta | *COI-5P* |
| U04021 | *Chondria dasyphylla* | Rhodophyta | *rbc*L |
| AY172578 | *Chondria californica* | Rhodophyta | *rbc*L |
| KU220791 | *Chondria armata* | Rhodophyta | *rbc*L |
| MT039608 | *Chondria arcuata* | Rhodophyta | *rbc*L |
| MT039609 | *Chondria arcuata* | Rhodophyta | *rbc*L |
| PX620351 | *Chondria arcuata* | Rhodophyta | *rbc*L |
| AF321113 | *Caulacanthus okamurae* | Rhodophyta | *rbc*L |
| OQ136637 | *Caulacanthus okamurae* | Rhodophyta | *rbc*L |
| KY940550 | *Caulacanthus okamurae* | Rhodophyta | *rbc*L |
| U22099 | *Caulacanthus ustulatus* | Rhodophyta | *rbc*L |
| U04188 | *Caulacanthus ustulatus* | Rhodophyta | *rbc*L |
| DQ374290 | *Centroceras gasparrinii* | Rhodophyta | *rbc*L |
| DQ374298 | *Centroceras gasparrinii* | Rhodophyta | *rbc*L |
| DQ374305 | *Centroceras gasparrinii* | Rhodophyta | *rbc*L |
| HM240513 | *Centroceras gasparrinii* | Rhodophyta | *rbc*L |
| KU220776 | *Centroceras gasparrinii* | Rhodophyta | *rbc*L |
| DQ374325 | *Centroceras clavulatum* | Rhodophyta | *rbc*L |
| DQ374323 | *Centroceras tetrachotomum* | Rhodophyta | *rbc*L |
| GQ252468 | *Ceramium tenuissimum* | Rhodophyta | *rbc*L |
| AF439290 | *Ceramium virgatum* | Rhodophyta | *rbc*L |
| DQ110904 | *Ceramium secundatum* | Rhodophyta | *rbc*L |
| AB491670 | *Glaucocystis nostochinearum* | Rhodophyta | *rbc*L |
| KF631393 | *Cyanophora paradoxa* | Rhodophyta | *rbc*L |
| KJ202103 | *Antithamnion hubbsii* | Rhodophyta | *UPA* |
| HQ421563 | *Antithamnion antillanum* | Rhodophyta | *UPA* |
| HQ421594 | *Antithamnion decipiens* | Rhodophyta | *UPA* |
| HQ421660 | *Antithamnion erucacladellum* | Rhodophyta | *UPA* |
| KF367775 | *Ceramium diaphanum* | Rhodophyta | *UPA* |
| KC795893 | *Ceramium tenerrimum* | Rhodophyta | *UPA* |
| KC795896 | *Ceramium japonicum* | Rhodophyta | *UPA* |
| MW142540 | *Elymus athericus* | Tracheophyta | *ITS* rRNA |
| OK166596 | *Elymus athericus* | Tracheophyta | *ITS* rRNA |
| EF396960 | *Elymus canadensis* | Tracheophyta | *ITS* rRNA |
| FJ793073 | *Elymus repens* | Tracheophyta | *ITS* rRNA |
| JX983685 | *Limonium gmelinii* | Tracheophyta | *ITS* rRNA |
| JX983691 | *Limonium gmelinii* | Tracheophyta | *ITS* rRNA |
| MH582614 | *Limonium virgatum* | Tracheophyta | *ITS* rRNA |
| MW290502 | *Limonium vulgare* | Tracheophyta | *ITS* rRNA |
| MW290501 | *Limoniumrbonense* | Tracheophyta | *ITS* rRNA |
| AF102274 | *Zostera marina* | Tracheophyta | *ITS* rRNA |
| AY077986 | *Zostera marina* | Tracheophyta | *ITS* rRNA |
| PZ361570 | *Artemisia maritima* | Tracheophyta | *ITS* rRNA |
| PV900068 | *Artemisia borotalensis* | Tracheophyta | *ITS* rRNA |
| AF061389 | *Artemisia vulgaris* | Tracheophyta | *ITS* rRNA |
| AJ296424 | *Artemisia vulgaris* | Tracheophyta | *ITS* rRNA |
| HQ735410 | *Artemisia annua* | Tracheophyta | *ITS* rRNA |
| HQ735412 | *Artemisia annua* | Tracheophyta | *ITS* rRNA |
| MG217549 | *Artemisia tridentata* | Tracheophyta | *ITS* rRNA |
| AM398849 | *Artemisia arctica* | Tracheophyta | *ITS* rRNA |
| JN895331 | *Halimione portulacoides* | Tracheophyta | *mat*K |
| DQ468648 | *Halimione portulacoides* | Tracheophyta | *mat*K |
| AY042550 | *Atriplex patula* | Tracheophyta | *mat*K |
| PP338755 | *Atriplex hortensis* | Tracheophyta | *mat*K |
| JX517608 | *Atriplex canescens* | Tracheophyta | *mat*K |
| PQ272986 | *Salsola foliosa* | Tracheophyta | *mat*K |
| MK954263 | *Salsola lanata* | Tracheophyta | *mat*K |
| AY514843 | *Salsola kali* | Tracheophyta | *mat*K |
| HM850761 | *Salsola kali* | Tracheophyta | *mat*K |
| MK125126 | *Salsola kali* | Tracheophyta | *mat*K |
| MF190426 | *Limonium gmelinii* | Tracheophyta | *mat*K |
| MF190428 | *Limonium gmelinii* | Tracheophyta | *mat*K |
| AF204853 | *Limoniumrbonense* | Tracheophyta | *mat*K |
| GQ248144 | *Limonium rigualii* | Tracheophyta | *mat*K |
| OP821366 | *Salicornia fruticosa* | Tracheophyta | *psbA-trnH* |
| OP821353 | *Salicornia procumbens* | Tracheophyta | *psbA-trnH* |
| OP821354 | *Salicornia procumbens* | Tracheophyta | *psbA-trnH* |
| MT873731 | *Suaeda japonica* | Tracheophyta | *psbA-trnH* |
| MT873720 | *Suaeda maritima* | Tracheophyta | *psbA-trnH* |
| PV898255 | *Halostachys caspica* | Tracheophyta | *psbA-trnH* |
| PV898256 | *Halostachys caspica* | Tracheophyta | *psbA-trnH* |
| KR263134 | *Ruppia cirrhosa* | Tracheophyta | *rbc*L |
| MG227122 | *Ruppia cirrhosa* | Tracheophyta | *rbc*L |
| DQ859175 | *Ruppia cirrhosa* | Tracheophyta | *rbc*L |
| JN225335 | *Cymodocea serrulata* | Tracheophyta | *rbc*L |
| JN225334 | *Cymodocea rotundata* | Tracheophyta | *rbc*L |
| KF488487 | *Cymodocea nodosa* | Tracheophyta | *rbc*L |
| KF488486 | *Cymodocea nodosa* | Tracheophyta | *rbc*L |
| HM587582 | *Atriplex australasica* | Tracheophyta | *rbc*L |
| HM587583 | *Atriplex centralasiatica* | Tracheophyta | *rbc*L |
| JN225350 | *Zostera noltii* | Tracheophyta | *rbc*L |
| U80733 | *Zostera noltii* | Tracheophyta | *rbc*L |
| U80734 | *Zostera marina* | Tracheophyta | *rbc*L |
| JN225353 | *Zostera marina* | Tracheophyta | *rbc*L |
| KF488490 | *Cymodocea rotundata* | Tracheophyta | *rbc*L |
| KF488491 | *Cymodocea serrulata* | Tracheophyta | *rbc*L |
| JN113277 | *Ruppia cirrhosa* | Tracheophyta | *rbc*L |
| MZ466378 | *Ruppia cirrhosa* | Tracheophyta | *rbc*L |
| JN113278 | *Ruppia maritima* | Tracheophyta | *rbc*L |
| U03729 | *Ruppia maritima* | Tracheophyta | *rbc*L |
| HQ901576 | *Ruppia maritima* | Tracheophyta | *rbc*L |
| GU724242 | *Artemisia scoparia* | Tracheophyta | *rbc*L |
| KX582024 | *Artemisia scoparia* | Tracheophyta | *rbc*L |
| KX282552 | *Artemisia scoparia* | Tracheophyta | *rbc*L |
| KX282551 | *Artemisia scoparia* | Tracheophyta | *rbc*L |
| KX582020 | *Artemisia pubescens* | Tracheophyta | *rbc*L |
| MN185069 | *Artemisia japonica* | Tracheophyta | *rbc*L |
| MG223735 | *Artemisia glomerata* | Tracheophyta | *rbc*L |
| MK348958 | *Artemisia absinthium* | Tracheophyta | *rbc*L |
| HM587584 | *Atriplex glauca* | Tracheophyta | *rbc*L |
| KM360659 | *Atriplex portulacoides* | Tracheophyta | *rbc*L |
| MN810509 | *Atriplex portulacoides* | Tracheophyta | *rbc*L |
| JN891002 | *Halimione portulacoides* | Tracheophyta | *rbc*L |
| OQ832174 | *Atriplex gardneri* | Tracheophyta | *rbc*L |
| MN810488 | *Atriplex clivicola* | Tracheophyta | *rbc*L |
| MN810501 | *Atriplex peruviana* | Tracheophyta | *rbc*L |
| AY216629 | *Juncus maritimus* | Tracheophyta | *rbc*L |
| AY216613 | *Juncus gerardii* | Tracheophyta | *rbc*L |
| JN891642 | *Juncus maritimus* | Tracheophyta | *rbc*L |
| AM999842 | *Juncus acutus* | Tracheophyta | *rbc*L |
| JN891719 | *Juncus acutus* | Tracheophyta | *rbc*L |
| MG227439 | *Juncus gerardii* | Tracheophyta | *rbc*L |
| MG226178 | *Juncus gerardii* | Tracheophyta | *rbc*L |
| MG226269 | *Juncus gerardii* | Tracheophyta | *rbc*L |
| KX678617 | *Juncus gerardii* | Tracheophyta | *rbc*L |
| JN891720 | *Juncus acutus* | Tracheophyta | *rbc*L |
| JN893492 | *Juncus acutus* | Tracheophyta | *rbc*L |
| OQ851353 | *Juncus acutus* | Tracheophyta | *rbc*L |
| AJ286364 | *Limoniumrbonense* | Tracheophyta | *rbc*L |
| KJ608053 | *Limonium virgatum* | Tracheophyta | *rbc*L |
| MH582735 | *Limonium vulgare* | Tracheophyta | *rbc*L |
| MT984450 | *Limonium vulgare* | Tracheophyta | *rbc*L |
| Y16904 | *Limonium vulgare* | Tracheophyta | *rbc*L |
| AB917056 | *Limonium virgatum* | Tracheophyta | *rbc*L |
| MW289886 | *Limoniumrbonense* | Tracheophyta | *rbc*L |
| NC_066030 | *Salicornia fruticosa* | Tracheophyta | *rbc*L |
| ON684503 | *Salicornia fruticosa* | Tracheophyta | *rbc*L |
| JQ412418 | *Sarcocornia perennis* | Tracheophyta | *rbc*L |
| HM630105 | *Salsola soda* | Tracheophyta | *rbc*L |
| PP754489 | *Soda foliosa* | Tracheophyta | *rbc*L |
| PQ272990 | *Soda foliosa* | Tracheophyta | *rbc*L |
| AY270129 | *Salsola kali* | Tracheophyta | *rbc*L |
| HM850332 | *Salsola kali* | Tracheophyta | *rbc*L |
| JN893159 | *Salsola kali* | Tracheophyta | *rbc*L |
| JN893640 | *Salsola kali* | Tracheophyta | *rbc*L |
| KX789408 | *Salsola kali* | Tracheophyta | *rbc*L |
| MG247392 | *Salsola kali* | Tracheophyta | *rbc*L |
| MH598919 | *Salsola kali* | Tracheophyta | *rbc*L |
| MG248444 | *Spergularia media* | Tracheophyta | *rbc*L |
| OY804563 | *Spergularia marina* | Tracheophyta | *rbc*L |
| OP353921 | *Spergularia marina* | Tracheophyta | *rbc*L |
| OY734961 | *Spergularia bocconei* | Tracheophyta | *rbc*L |
| OQ172709 | *Spergularia bocconei* | Tracheophyta | *rbc*L |
| JN891830 | *Spergularia media* | Tracheophyta | *rbc*L |
| JN891963 | *Spergularia media* | Tracheophyta | *rbc*L |
| HM850380 | *Spergularia bocconei* | Tracheophyta | *rbc*L |
| OQ172710 | *Spergularia bocconei* | Tracheophyta | *rbc*L |
| PP129849 | *Spergularia bocconei* | Tracheophyta | *rbc*L |
| KC483744 | *Puccinellia distans* | Tracheophyta | *rbc*L |
| MG226113 | *Puccinellia distans* | Tracheophyta | *rbc*L |
| MG228057 | *Puccinellia distans* | Tracheophyta | *rbc*L |
| MG227498 | *Phragmites australis* | Tracheophyta | *rbc*L |
| MT886091 | *Phragmites australis* | Tracheophyta | *rbc*L |
| KM360930 | *Phragmites australis* | Tracheophyta | *rbc*L |
| JX985731 | *Suaeda fruticosa* | Tracheophyta | *rbc*L |
| KF679785 | *Suaeda fruticosa* | Tracheophyta | *rbc*L |
| KF679786 | *Suaeda fruticosa* | Tracheophyta | *rbc*L |
| JN891221 | *Suaeda maritima* | Tracheophyta | *rbc*L |
| JX997825 | *Suaeda maritima* | Tracheophyta | *rbc*L |
| JN893488 | *Suaeda maritima* | Tracheophyta | *rbc*L |
| JN891222 | *Suaeda maritima* | Tracheophyta | *rbc*L |
| JN893487 | *Suaeda vera* | Tracheophyta | *rbc*L |
| JN891228 | *Suaeda vera* | Tracheophyta | *rbc*L |
| KR057197 | *Suaeda maritima* | Tracheophyta | *rbc*L |
| MG247562 | *Suaeda maritima* | Tracheophyta | *rbc*L |
| GQ452331 | *Triglochin barrelieri* | Tracheophyta | *rbc*L |
| GQ452332 | *Triglochin elongata* | Tracheophyta | *rbc*L |
| GQ452333 | *Triglochin maritima* | Tracheophyta | *rbc*L |
| GQ452334 | *Triglochin palustris* | Tracheophyta | *rbc*L |
| GQ452336 | *Triglochin striata* | Tracheophyta | *rbc*L |
| JX848473 | *Triglochin maritima* | Tracheophyta | *rbc*L |
| HQ901578 | *Triglochin maritima* | Tracheophyta | *rbc*L |
| AB088811 | *Triglochin maritima* | Tracheophyta | *rbc*L |
| DQ859176 | *Triglochin palustris* | Tracheophyta | *rbc*L |
| OZ078213 | *Triglochin palustris* | Tracheophyta | *rbc*L |
| PP234593 | *Triglochin palustris* | Tracheophyta | *rbc*L |
| NC_088567 | *Tripolium pannonicum* | Tracheophyta | *rbc*L |
| LC778122 | *Tripolium pannonicum* | Tracheophyta | *rbc*L |
| MF963323 | *Euthamia occidentalis* | Tracheophyta | *rbc*L |
| KM212122 | *Euthamia graminifolia* | Tracheophyta | *rbc*L |
| DQ845682 | *Anthoceros agrestis* | Tracheophyta | *rbc*L |
| AY463053 | *Anthoceros laminiferus* | Tracheophyta | *rbc*L |
| DQ845681 | *Anthoceros punctatus* | Tracheophyta | *rbc*L |
| U87063 | *Anthoceros punctatus* | Tracheophyta | *rbc*L |
| DQ845679 | *Anthoceros lamellatus* | Tracheophyta | *rbc*L |
| JN893641 | *Inula crithmoides* | Tracheophyta | *rbc*L |
| NC_066031 | *Limbarda crithmoides* | Tracheophyta | *rbc*L |
| NMW3388 | *juncus acutus* | Tracheophyta | *rbc*L |
| NMW3418 | *juncus maritimus* | Tracheophyta | *rbc*L |
| EDNA22-0062645 | *juncus gerardii* | Tracheophyta | *rbc*L |
| BGE_00278 A10 | *Zostera marina* | Tracheophyta | *rbc*L |
| BGE_00278 A03 | *Zostera noltii* | Tracheophyta | *rbc*L |
| KF488492 | *Cymodocea serrulata* | Tracheophyta | *rbc*L |
| EDNA21-0061572 | *Ruppia maritima* | Tracheophyta | *rbc*L |
| EDNA21-0061102 | *Triglochin palustris* | Tracheophyta | *rbc*L |
| EDNA21-0061462 | *Triglochin maritima* | Tracheophyta | *rbc*L |
| CCDB-22990-C10 | *Euthamia occidentalis* | Tracheophyta | *rbc*L |
| CCDB-18322-E10 | *Euthamia graminifolia* | Tracheophyta | *rbc*L |
| EDNA21-0061451 | *Tripolium pannonicum* | Tracheophyta | *rbc*L |
| BPTPS107 | *Limbarda crithmoides* | Tracheophyta | *rbc*L |
| EDNA21-0061571 | *Halimione portulacoides* | Tracheophyta | *rbc*L |
| HM587581 | *Atriplex aucherii* | Tracheophyta | *rbc*L |
| CCDB-18328-D10 | *Atriplex gardneri* | Tracheophyta | *rbc*L |
| CCDB-26262-H05 | *Atriplex glauca* | Tracheophyta | *rbc*L |
| CCDB-26262-E02 | *Atriplex centralasiatica* | Tracheophyta | *rbc*L |
| EDNA21-0061526 | *Salsola kali* | Tracheophyta | *rbc*L |
| EDNA20-0060189 | *Suaeda vera* | Tracheophyta | *rbc*L |
| KF500483 | *Suaeda heterophylla* | Tracheophyta | *rbc*L |
| BABY-09666 | *Suaeda maritima* | Tracheophyta | *rbc*L |
| BPTPS106 | *Salicornia fruticosa* | Tracheophyta | *rbc*L |
| OQ851357 | *Salicornia perennis* | Tracheophyta | *rbc*L |
| MG249663 | *Salicornia perennis* | Tracheophyta | *rbc*L |
| GQ452335 | *Triglochin rheophila* | Tracheophyta | *rbc*L |
| JN890621 | *Elymus athericus* | Tracheophyta | *rbc*L |
| JN891933 | *Elymus athericus* | Tracheophyta | *rbc*L |
| EF672515 | *Vaucheria litorea* | Xantophyceae | *COI* |
| AB000212 | *Vaucheria sessilis* | Xantophyceae | *COI* |
| AB000204 | *Botrydium granulatum* | Xantophyceae | *COI* |
| AB000207 | *Monodus sp* | Xantophyceae | *COI* |
| AB000210 | *Ophiocytium majus* | Xantophyceae | *COI* |
| AB000206 | *Heterococcus caespitosus* | Xantophyceae | *COI* |
| AB000211 | *Tribonema aequale* | Xantophyceae | *COI* |
| KF281380 | *Scytosiphon promiscuus holotype* | Xantophyceae | *COI* |
| GWS011886 | *Scytosiphon promiscuus* | Xantophyceae | *COI* |
| AF015589 | *Vaucheria bursata* | Xantophyceae | *rbc*L |
| AF476937 | *Vaucheria bursata* | Xantophyceae | *rbc*L |
| AF207527 | *Vaucheria litorea* | Xantophyceae | *rbc*L |
| AF476935 | *Vaucheria litorea* | Xantophyceae | *rbc*L |
| AF476943 | *Vaucheria aversa* | Xantophyceae | *rbc*L |
| AF064743 | *Botrydium stoloniferum* | Xantophyceae | *rbc*L |
| AJ874699 | *Ophiocytium majus* | Xantophyceae | *rbc*L |
| MK482707 | *Ophiocytium majus* | Xantophyceae | *rbc*L |
| AF084610 | *Heterococcus caespitosus* | Xantophyceae | *rbc*L |
| EF455943 | *Heterococcus caespitosus* | Xantophyceae | *rbc*L |
| AF084611 | *Tribonema aequale* | Xantophyceae | *rbc*L |
| AF465709 | *Tribonema intermixum* | Xantophyceae | *rbc*L |
| AF207811 | *Scytosiphon lomentaria* | Xantophyceae | *rbc*L |
| AY372954 | *Scytosiphon lomentaria* | Xantophyceae | *rbc*L |

**Supplementary Table 7. Species identified during the 2023 and 2024 surveys and their distribution in the Venice Lagoon.** For each species, the table reports the group (R = Rhodophyta, H = Halophytes, C = Chlorophyta, P = Phaeophyceae, X = Xantophyceae, S = Seagrasses), the lagoon sectors (southern, central, and northern) and sampling seasons (spring, summer, and autumn) in which it was recorded.

| **Species** | **Group** | **Spring** | | | **Summer** | | | **Autumn** | | |
| --- | --- | --- | --- | --- | --- | --- | --- | --- | --- | --- |
|  |  | **Southern** | **Central** | **Northern** | **Southern** | **Central** | **Northern** | **Southern** | **Central** | **Northern** |
| *Agardhiella subulata* | R | x |  |  |  |  |  |  |  |  |
| *Aglaothamnion halliae* | R |  |  |  |  |  |  |  | x |  |
| *Aglaothamnion sp.* | R | x |  |  |  |  |  |  |  |  |
| *Alsidium corallinum* | R |  |  |  | x | x |  | x | x | x |
| *Antithamnion hubbsii* | R |  |  |  |  |  |  | x |  |  |
| *Antithamnion sp.* | R |  |  |  |  |  |  |  | x |  |
| *Artemisia sp.* | H |  |  |  |  |  |  | x |  | x |
| *Atriplex portulacoides* | H | x | x |  | x | x | x | x | x |  |
| *Blidingia marginata* | C |  | x |  |  | x |  |  |  |  |
| *Bryopsis muscosa* | C |  |  |  |  |  |  |  |  | x |
| *Bryopsis sp.* | C | x | x | x | x |  | x |  |  | x |
| *Carradoriella elongata* | R |  | x |  |  |  |  |  |  |  |
| *Caulacanthus okamurae* | R |  | x |  |  |  |  |  |  | x |
| *Centroceras gasparrinii* | R |  |  |  | x | x |  | x | x |  |
| *Chaetomorpha linum* | C |  |  |  |  |  |  |  |  |  |
| *Chaetomorpha sp.* | C | x | x | x | x |  | x | x | x | x |
| *Chondria arcuata* | R |  |  |  |  |  | x |  |  |  |
| *Chondria sp.* | R |  |  |  |  |  |  |  |  | x |
| *Cladophora sp.* | C | x | x |  | x |  | x |  | x |  |
| *Codium fragile* | C |  |  |  |  | x | x |  |  |  |
| *Cymodocea nodosa* | S | x | x |  | x | x |  | x | x | x |
| *Cystoseira sp.* | P | x | x |  |  |  |  |  |  |  |
| *Dasya sp.* | P | x | x |  | x |  |  |  |  |  |
| *Dictyopteris polypodioides* | P |  | x |  |  | x |  |  | x |  |
| *Dictyota dichotoma* | P | x | x |  |  |  |  | x | x |  |
| *Dictyota fascicola* | P | x | x |  | x |  |  |  |  |  |
| *Dictyota* sp. | P |  | x |  |  |  |  |  |  |  |
| *Ectocarpus siliculosus* | P | x | x | x |  |  |  |  | x |  |
| *Fucus virsoides* | P | x |  |  |  | x |  |  |  |  |
| *Gayralia oxysperma* | C | x |  |  |  |  |  |  |  |  |
| *Gelidium crinale* | R |  |  |  |  | x |  |  |  |  |
| *Gongolaria barbata* | P | x | x |  |  | x |  |  |  | x |
| *Gracilaria gracilis* | R |  |  |  |  |  |  | x |  |  |
| *Gracilaria multipartita* | R | x |  |  |  |  |  |  |  |  |
| *Heterosiphonia* sp. | R | x |  |  | x |  |  |  |  |  |
| *Hypnea cervicornis* | R |  |  |  | x | x | x | x | x |  |
| *Hypnea musciformis* | R |  |  |  |  |  |  |  |  | x |
| *Juncus acutus* | H |  |  |  | x |  |  |  |  |  |
| *Juncus gerardii* | H |  |  |  | x |  |  |  |  |  |
| *Juncus maritimus* | H | x | x |  | x |  |  | x |  | x |
| *Kapraunia schneideri* | R |  |  |  |  |  |  | x | x |  |
| *Limbarda crithmoides* | H | x | x |  | x | x | x | x | x | x |
| *Limonium* sp. | H | x | x |  | x | x | x | x | x | x |
| *Limonium narbonense* | H | x | x |  |  | x |  |  |  |  |
| *Limonium virgatum* | H |  |  |  |  |  |  | x |  |  |
| *Melanothamnus japonicus* | R |  |  |  | x |  |  |  |  |  |
| *Nitophyllum punctatum* | R |  |  |  |  |  |  |  | x |  |
| *Osmundea truncata* | R |  | x | x |  |  |  |  |  |  |
| *Palisada* sp. | R |  |  |  |  | x |  |  | x |  |
| *Phragmites australis* | H |  |  |  | x |  |  |  |  |  |
| *Phyllophora sicula* | R |  | x |  | x | x | x |  |  | x |
| *Polysiphonia* sp. | R |  |  |  |  | x | x | x | x |  |
| *Puccinellia distans* | H | x | x |  | x | x | x | x | x | x |
| *Pyropia* sp. | R |  | x | x |  |  |  |  |  |  |
| *Radicilingua* sp. | R | x | x | x |  |  |  | x | x |  |
| *Ruppia* sp. | S |  |  |  |  | x |  | x |  |  |
| *Salicornia perennis* | H | x |  |  |  |  |  |  |  |  |
| *Salicornia procumbens* | H |  |  |  |  |  |  |  | x |  |
| *Salicornia fruticosa* | H | x | x |  | x |  |  | x |  | x |
| *Salicornia* sp. | H | x | x | x | x | x | x | x | x | x |
| *Salsola* sp. | H |  |  |  |  |  |  | x |  |  |
| *Sargassum muticum* | P |  | x |  |  |  |  |  |  |  |
| *Scytosiphon lomentaria* | P | x | x |  |  |  |  |  |  |  |
| *Spergularia media* | H |  | x |  | x |  |  | x |  |  |
| *Spermothamnion cymosum* | R |  |  |  |  | x |  |  |  |  |
| *Sporobolus* sp. | H | x | x |  | x | x | x | x | x | x |
| *Suaeda maritima* | H |  |  |  | x |  |  | x | x |  |
| *Thinopyrum acutum* | H | x |  |  | x |  |  | x |  |  |
| *Triglochin maritima* | H |  |  |  | x |  |  |  |  |  |
| *Tripolium pannonicum* | H | x | x |  | x |  | x | x | x | x |
| *Ulva compressa* | C | x | x | x |  | x |  | x | x | x |
| *Ulva flexuosa* | C |  |  |  |  |  |  |  |  |  |
| *Ulva lacinulata* | C | x | x | x | x | x |  | x | x | x |
| *Ulva rigida* | C |  |  |  |  |  |  |  |  |  |
| *Ulva* sp. | C | x |  |  | x | x | x |  |  |  |
| *Vaucheria litorea* | X |  |  |  |  |  |  |  |  |  |
| *Zostera marina* | S | x | x |  |  | x | x | x | x |  |
| *Zostera noltii* | S | x | x | x |  |  |  |  |  | x |

**Supplementary Table 8. GenBank accession numbers of sequences generated during the monitoring surveys.** For each identified species, the table reports the corresponding GenBank accession number and molecular marker.

| **Species** | **Accession numbers** | **Gene marker** | | | | | | | | |
| --- | --- | --- | --- | --- | --- | --- | --- | --- | --- | --- |
|  |  | *COI*  *5P* | *UPA* | *ITS* rRNA | *rbc*L | *mat*K | *tuf*A | *18S* rRNA | *psbA-trnH* | *trnL-trnF* |
| *Agardhiella subulata* | PZ305589, PZ305590 | x |  |  |  |  |  |  |  |  |
| *Aglaothamnion halliae* | PZ305577 | x |  |  |  |  |  |  |  |  |
| *Alsidium corallinum* | PZ305591, PZ305592, PZ305593, PZ305594, PZ305595, PZ305596, PZ305597, PZ305598 | x |  |  |  |  |  |  |  |  |
| *Antithamnion hubbsii* | PZ561528 | x |  |  |  |  |  |  |  |  |
|  | PZ534468 |  | x |  |  |  |  |  |  |  |
| *Artemisia* sp. | PZ543477 |  |  | x |  |  |  |  |  |  |
|  | PZ547424 |  |  |  | x |  |  |  |  |  |
| *Atriplex portulacoides* | PZ547373, PZ547374, PZ547376, PZ547377, PZ547378, PZ547379 |  |  |  |  | x |  |  |  |  |
|  | PZ318371, PZ547430, PZ547431, PZ547438, PZ547439, PZ547440, PZ547441 |  |  |  | x |  |  |  |  |  |
| *Blidingia marginata* | PZ237727, PZ237728 |  |  |  |  |  | x |  |  |  |
| *Bryopsis muscosa* | PZ237725 |  |  |  |  |  | x |  |  |  |
| *Bryopsis* sp. 1 | PZ547454, PZ547455, PZ547458 |  |  |  |  |  | x |  |  |  |
| *Bryopsis* sp. 2 | PZ547456 |  |  |  |  |  | x |  |  |  |
| *Bryopsis* sp. 3 | PZ547457 |  |  |  |  |  | x |  |  |  |
| *Carradoriella elongata* | PZ305580 | x |  |  |  |  |  |  |  |  |
| *Caulacanthus okamurae* | PZ548663, PZ561529, PZ548664 | x |  |  |  |  |  |  |  |  |
|  | PZ533869, PZ533870, PZ533871, PZ568856, PZ568862 |  |  |  | x |  |  |  |  |  |
| *Centroceras gasparrinii* | PZ533872, PZ533873, PZ568857, PZ568858, PZ568859, PZ568860, PZ568861 |  |  |  | x |  |  |  |  |  |
| *Chaetomorpha linum* | PZ530647, PZ530648 |  |  |  |  |  |  | x |  |  |
|  | PZ547452, PZ547453 |  |  |  |  |  | x |  |  |  |
| *Chondria arcuata* | PZ548662 | x |  |  |  |  |  |  |  |  |
|  | PZ533868 |  |  |  | x |  |  |  |  |  |
| *Codium fragile* | PZ237723, PZ237724 |  |  |  |  |  | x |  |  |  |
| *Cymodocea nodosa* | PZ318347, PZ318348, PZ318349, PZ318350, PZ318351 |  |  |  | x |  |  |  |  |  |
| *Dictyopteris polypodioides* | PZ305570 | x |  |  |  |  |  |  |  |  |
| *Elymus athericus* | PZ547419, PZ547420, PZ547421, PZ547422 |  |  |  | x |  |  |  |  |  |
| *Gayralia oxysperma* | PZ237726 |  |  |  |  |  | x |  |  |  |
| *Gelidium crinale* | PZ305599 | x |  |  |  |  |  |  |  |  |
| *Gongolaria barbata* | PZ305571, PZ305572, PZ305573, PZ305574 | x |  |  |  |  |  |  |  |  |
| *Gracilaria gracilis* | PZ305600 | x |  |  |  |  |  |  |  |  |
| *Gracilaria multipartita* | PZ305601, PZ305602 | x |  |  |  |  |  |  |  |  |
| *Hypnea cervicornis* | PZ305585, PZ305586, PZ305587, PZ305588 | x |  |  |  |  |  |  |  |  |
| *Hypnea musciformis* | PZ305583, PZ305584 | x |  |  |  |  |  |  |  |  |
| *Juncus acutus* | PZ547399, PZ547400, PZ547401 |  |  |  | x |  |  |  |  |  |
| *Juncus gerardii* | PZ547398 |  |  |  | x |  |  |  |  |  |
| *Juncus maritimus* | PZ547402, PZ547403, PZ547404, PZ547405, PZ547406, PZ547407 |  |  |  | x |  |  |  |  |  |
| *Kapraunia schneideri* | PZ305579 | x |  |  |  |  |  |  |  |  |
| *Limbarda crithmoides* | PZ318364, PZ318365, PZ318366, PZ318367 |  |  |  | x |  |  |  |  |  |
| *Limonium narbonense* | PZ543478, PZ543479, PZ543480, PZ543481, PZ543482, PZ543487 |  |  | x |  |  |  |  |  |  |
|  | PZ547425, PZ547426, PZ547427, PZ547428, PZ547429, PZ547451 |  |  |  | x |  |  |  |  |  |
| *Limonium virgatum* | PZ318368 |  |  |  | x |  |  |  |  |  |
| *Melanothamnus japonicus* | PZ547459 | x |  |  |  |  |  |  |  |  |
| *Osmundea truncata* | PZ305581, PZ305582 | x |  |  |  |  |  |  |  |  |
| *Palisada* sp. | PZ548665 | x |  |  |  |  |  |  |  |  |
| *Phragmites australis* | PZ318354 |  |  |  | x |  |  |  |  |  |
| *Puccinellia distans* | PZ547408, PZ547409, PZ547410, PZ547411, PZ547412, PZ547413, PZ547414, PZ547415, PZ547416, PZ547417, PZ547418 |  |  |  | x |  |  |  |  |  |
| *Pyropia* sp. | PZ629021 |  |  |  |  |  | x |  |  |  |
| *Ruppia* sp. | PZ547395, PZ547396, PZ547397 |  |  |  | x |  |  |  |  |  |
| *Salicornia fruticosa* | PZ547380, PZ547381, PZ547383, PZ547384, PZ547386, PZ547387, PZ547388, PZ547389, PZ547390, PZ547391, PZ547392, PZ547393, PZ547394 |  |  |  |  |  |  |  | x |  |
|  | PZ547432, PZ547433, PZ547435, PZ547436, PZ547442, PZ547443, PZ547444, PZ547445, PZ547446, PZ547447, PZ547448, PZ547449, PZ547450 |  |  |  | x |  |  |  |  |  |
| *Salicornia perennis* | PZ547382 |  |  |  |  |  |  |  | x |  |
|  | PZ547434 |  |  |  | x |  |  |  |  |  |
| *Salicornia procumbens* | PZ547385 |  |  |  |  |  |  |  | x |  |
|  | PZ547437 |  |  |  | x |  |  |  |  |  |
| *Sargassum muticum* | PZ305575, PZ305576 | x |  |  |  |  |  |  |  |  |
| *Scytosiphon lomentaria* | PZ548661 | x |  |  |  |  |  |  |  |  |
|  | PZ533867 |  |  |  | x |  |  |  |  |  |
| *Salsola* sp. | PZ547375 |  |  |  |  | x |  |  |  |  |
|  | PZ547423 |  |  |  | x |  |  |  |  |  |
| *Spergularia media* | PZ318369, PZ318370 |  |  |  | x |  |  |  |  |  |
| *Spermothamnion cymosum* | PZ305578 | x |  |  |  |  |  |  |  |  |
| *Sporobolus anglicus* | PZ166190, PZ166193, PZ166194, PZ166199, PZ166200, PZ166201, PZ166202, PZ166203, PZ166204, PZ166209, PZ166210, PZ166211, PZ166214, PZ166215, PZ166216, PZ166217, PZ166223, PZ166225, PZ166226, PZ166227, PZ166228, PZ166229, PZ166230, PZ166231, PZ166232, PZ166237, PZ166238, PZ166239, PZ166240, PZ166241, PZ166242, PZ166243, PZ166244, PZ166245, PZ166246, PZ166247, PZ166251, PZ166252, PZ166253, PZ166254, PZ166258, PZ166259, PZ166260, PZ166261, PZ166262, PZ166265, PZ166266, PZ166267, PZ166268, PZ166271, PZ166272, PZ166273, PZ166274, PZ166278, PZ166279, PZ166283, PZ166284, PZ166285, PZ166289, PZ166290, PZ166291 |  |  |  |  |  |  |  |  | x |
| *Sporobolus maritimus* | PZ166195, PZ166196, PZ166197, PZ166198, PZ166205, PZ166206, PZ166208, PZ166212, PZ166213, PZ166218, PZ166219, PZ166221, PZ166222, PZ166233, PZ166234, PZ166235, PZ166236, PZ166248, PZ166249, PZ166250, PZ166255, PZ166256, PZ166257, PZ166263, PZ166264, PZ166269, PZ166270, PZ166275, PZ166276, PZ166277, PZ166280, PZ166281, PZ166282, PZ166286, PZ166287, PZ166288, PZ166292, PZ166293, PZ166294 |  |  |  |  |  |  |  |  | x |
| *Sporobolus x townsendii* | PZ166191, PZ166192, PZ166207, PZ166220, PZ166224 |  |  |  |  |  |  |  |  | x |
| *Suaeda maritima* | PZ318372, PZ318373, PZ318374, PZ318375, PZ318376 |  |  |  | x |  |  |  |  |  |
| *Thinopyrum acutum* | PZ543483, PZ543484, PZ543485, PZ543486 |  |  | x |  |  |  |  |  |  |
| *Triglochin maritima* | PZ318352, PZ318353 |  |  |  | x |  |  |  |  |  |
| *Tripolium pannonicum* | PZ318355, PZ318356, PZ318357, PZ318358, PZ318359, PZ318360, PZ318361, PZ318362, PZ318363 |  |  |  | x |  |  |  |  |  |
| *Ulva compressa* | PZ237729, PZ237730, PZ305550, PZ305551, PZ305552, PZ305553, PZ305554 |  |  |  |  |  | x |  |  |  |
| *Ulva flexuosa* | PZ305556, PZ305557 |  |  |  |  |  | x |  |  |  |
| *Ulva lacinulata* | PZ305558, PZ305559, PZ305560, PZ305561, PZ305562, PZ305563, PZ305564, PZ305565, PZ305566, PZ305567, PZ305568, PZ305569 |  |  |  |  |  | x |  |  |  |
| *Ulva rigida* | PZ305555 |  |  |  |  |  | x |  |  |  |
| *Vaucheria litorea* | PZ548660 | x |  |  |  |  |  |  |  |  |
|  | PZ533866 |  |  |  | x |  |  |  |  |  |
| *Zostera marina* | PZ318343, PZ318344 |  |  |  | x |  |  |  |  |  |
| *Zostera noltii* | PZ318345, PZ318346 |  |  |  | x |  |  |  |  |  |

**Supplementary Table 9. Molecular and flow-cytometric characterization of the collected Sporobolus specimens.** For each site (LS, Southern Lagoon; LC, Central Lagoon; LN, Northern Lagoon), the table reports the number of plants collected, the trnL–trnF sequence-alignment result, the number of peaks detected by flow cytometry using S. anglicus as a reference, and the final taxonomic identification based on the combined evidence. A single peak indicates a genome size consistent with that of S. anglicus, whereas two peaks indicate the presence of nuclei with different genome sizes. Flow-cytometric analysis was not performed when the chloroplast marker identified the specimen as S. maritimus.

| Site | Plant | Chloroplast marker | Peaks at the cytofluorometer analysis | Identification |
| --- | --- | --- | --- | --- |
| LS 1 | 1 | *S. anglicus/S. alterniflorus* | 1 | *S. anglicus* |
|  | 2 | *S. anglicus/S. alterniflorus* | 1 | *S. anglicus* |
| LS 2 | 3 | *S. anglicus/S. alterniflorus* | 1 | *S. anglicus* |
|  | 4 | *S. anglicus/S. alterniflorus* | 1 | *S. anglicus* |
|  | 5 | *S. anglicus/S. alterniflorus* | 1 | *S. anglicus* |
|  | 6 | *S. maritimus* | \ | *S. maritimus* |
|  | 7 | *S. maritimus* | \ | *S. maritimus* |
|  | 8 | *S. maritimus* | \ | *S. maritimus* |
| LS 3 | 9 | *S. anglicus/S. alterniflorus* | 1 | *S. anglicus* |
|  | 10 | *S. anglicus/S. alterniflorus* | 1 | *S. anglicus* |
|  | 11 | *S. anglicus/S. alterniflorus* | 1 | *S. anglicus* |
|  | 12 | *S. anglicus/S. alterniflorus* | 1 | *S. anglicus* |
|  | 13 | *S. anglicus/S. alterniflorus* | 1 | *S. anglicus* |
|  | 14 | *S. anglicus/S. alterniflorus* | 1 | *S. anglicus* |
| LS 4 | 15 | *S. anglicus/S. alterniflorus* | 1 | *S. anglicus* |
|  | 16 | *S. anglicus/S. alterniflorus* | 1 | *S. anglicus* |
|  | 17 | *S. anglicus/S. alterniflorus* | 1 | *S. anglicus* |
|  | 18 | *S. anglicus/S. alterniflorus* | 1 | *S. anglicus* |
|  | 19 | *S. anglicus/S. alterniflorus* | 1 | *S. anglicus* |
|  | 20 | *S. maritimus* | \ | *S. maritimus* |
|  | 21 | *S. maritimus* | \ | *S. maritimus* |
| LS 5 | 22 | *S. maritimus* | \ | *S. maritimus* |
|  | 23 | *S. maritimus* | \ | *S. maritimus* |
|  | 24 | *S. anglicus/S. alterniflorus* | 1 | *S. anglicus* |
|  | 25 | *S. anglicus/S. alterniflorus* | 2 | *S. x townsendii* |
|  | 26 | *S. anglicus/S. alterniflorus* | 1 | *S. anglicus* |
|  | 27 | *S. anglicus/S. alterniflorus* | 1 | *S. anglicus* |
| LS 6 | 28 | *S. anglicus/S. alterniflorus* | 1 | *S. anglicus* |
|  | 29 | *S. anglicus/S. alterniflorus* | 1 | *S. anglicus* |
|  | 30 | *S. anglicus/S. alterniflorus* | 1 | *S. anglicus* |
|  | 31 | *S. anglicus/S. alterniflorus* | 1 | *S. anglicus* |
|  | 32 | *S. maritimus* | \ | *S. maritimus* |
|  | 33 | *S. maritimus* | \ | *S. maritimus* |
|  | 34 | *S. maritimus* | \ | *S. maritimus* |
| LC 7 | 35 | *S. anglicus/S. alterniflorus* | 1 | *S. anglicus* |
|  | 36 | *S. anglicus/S. alterniflorus* | 2 | *S. x townsendii* |
|  | 37 | *S. anglicus/S. alterniflorus* | 2 | *S. x townsendii* |
| LC 8 | 38 | *S. anglicus/S. alterniflorus* | 1 | *S. anglicus* |
|  | 39 | *S. anglicus/S. alterniflorus* | 1 | *S. anglicus* |
|  | 40 | *S. anglicus/S. alterniflorus* | 1 | *S. anglicus* |
|  | 41 | *S. maritimus* | \ | *S. maritimus* |
|  | 42 | *S. maritimus* | \ | *S. maritimus* |
|  | 43 | *S. maritimus* | \ | *S. maritimus* |
|  | 44 | *S. anglicus/S. alterniflorus* | 2 | *S. x townsendii* |
| LC 9 | 45 | *S. anglicus/S. alterniflorus* | 1 | *S. anglicus* |
|  | 46 | *S. anglicus/S. alterniflorus* | 1 | *S. anglicus* |
|  | 47 | *S. anglicus/S. alterniflorus* | 1 | *S. anglicus* |
|  | 48 | *S. maritimus* | \ | *S. maritimus* |
|  | 49 | *S. maritimus* | \ | *S. maritimus* |
| LC10 | 50 | *S. anglicus/S. alterniflorus* | 1 | *S. anglicus* |
|  | 51 | *S. anglicus/S. alterniflorus* | 1 | *S. anglicus* |
|  | 52 | *S. anglicus/S. alterniflorus* | 1 | *S. anglicus* |
| LC 11 | 53 | *S. anglicus/S. alterniflorus* | 1 | *S. anglicus* |
|  | 54 | *S. anglicus/S. alterniflorus* | 1 | *S. anglicus* |
|  | 55 | *S. anglicus/S. alterniflorus* | 1 | *S. anglicus* |
|  | 56 | *S. anglicus/S. alterniflorus* | 1 | *S. anglicus* |
|  | 57 | *S. maritimus* | \ | *S. maritimus* |
|  | 58 | *S. maritimus* | \ | *S. maritimus* |
|  | 59 | *S. anglicus/S. alterniflorus* | 2 | *S. x townsendii* |
| LC 12 | 60 | *S. anglicus/S. alterniflorus* | 1 | *S. anglicus* |
|  | 61 | *S. anglicus/S. alterniflorus* | 1 | *S. anglicus* |
|  | 62 | *S. maritimus* | \ | *S. maritimus* |
|  | 63 | *S. maritimus* | \ | *S. maritimus* |
|  | 64 | *S. maritimus* | \ | *S. maritimus* |
|  | 65 | *S. maritimus* | \ | *S. maritimus* |
| LN 13 | 66 | *S. anglicus/S. alterniflorus* | 1 | *S. anglicus* |
|  | 67 | *S. anglicus/S. alterniflorus* | 1 | *S. anglicus* |
|  | 68 | *S. anglicus/S. alterniflorus* | 1 | *S. anglicus* |
|  | 69 | *S. anglicus/S. alterniflorus* | 1 | *S. anglicus* |
|  | 70 | *S. maritimus* | \ | *S. maritimus* |
|  | 71 | *S. maritimus* | \ | *S. maritimus* |
| LN 14 | 72 | *S. anglicus/S. alterniflorus* | 1 | *S. anglicus* |
|  | 73 | *S. anglicus/S. alterniflorus* | 1 | *S. anglicus* |
|  | 74 | *S. anglicus/S. alterniflorus* | 1 | *S. anglicus* |
|  | 75 | *S. anglicus/S. alterniflorus* | 1 | *S. anglicus* |
|  | 76 | *S. maritimus* | \ | *S. maritimus* |
|  | 77 | *S. maritimus* | \ | *S. maritimus* |
|  | 78 | *S. maritimus* | \ | *S. maritimus* |
| LN 15 | 79 | *S. anglicus/S. alterniflorus* | 1 | *S. anglicus* |
|  | 80 | *S. anglicus/S. alterniflorus* | 1 | *S. anglicus* |
|  | 81 | *S. maritimus* | \ | *S. maritimus* |
|  | 82 | *S. maritimus* | \ | *S. maritimus* |
|  | 83 | *S. maritimus* | \ | *S. maritimus* |
| LN 16 | 84 | *S. anglicus/S. alterniflorus* | 1 | *S. anglicus* |
|  | 85 | *S. anglicus/S. alterniflorus* | 1 | *S. anglicus* |
|  | 86 | *S. anglicus/S. alterniflorus* | 1 | *S. anglicus* |
|  | 87 | *S. maritimus* | \ | *S. maritimus* |
|  | 88 | *S. maritimus* | \ | *S. maritimus* |
|  | 89 | *S. maritimus* | \ | *S. maritimus* |
| LN 17 | 90 | *S. anglicus/S. alterniflorus* | 1 | *S. anglicus* |
|  | 91 | *S. maritimus* | \ | *S. maritimus* |
|  | 92 | *S. anglicus/S. alterniflorus* | 1 | *S. anglicus* |
|  | 93 | *S. anglicus/S. alterniflorus* | 1 | *S. anglicus* |
|  | 94 | *S. anglicus/S. alterniflorus* | 1 | *S. anglicus* |
|  | 95 | *S. anglicus/S. alterniflorus* | 1 | *S. anglicus* |
|  | 96 | *S. anglicus/S. alterniflorus* | 1 | *S. anglicus* |
|  | 97 | *S. maritimus* | \ | *S. maritimus* |
|  | 98 | *S. maritimus* | \ | *S. maritimus* |
|  | 99 | *S. maritimus* | \ | *S. maritimus* |
| LN 18 | 100 | *S. anglicus/S. alterniflorus* | 1 | *S. anglicus* |
|  | 101 | *S. anglicus/S. alterniflorus* | 1 | *S. anglicus* |
|  | 102 | *S. anglicus/S. alterniflorus* | 1 | *S. anglicus* |
|  | 103 | *S. maritimus* | \ | *S. maritimus* |
|  | 104 | *S. maritimus* | \ | *S. maritimus* |
|  | 105 | *S. maritimus* | \ | *S. maritimus* |
